# Physioxia-modulated mesenchymal stem cells secretome has higher capacity to preserve neuronal network and translation processes in hypoxic-ischemic encephalopathy in vitro model

**DOI:** 10.1101/2024.11.26.625525

**Authors:** Inês Caramelo, Sandra I. Anjo, Vera M. Mendes, Ivan L. Salazar, Alexandra Dinis, Carla M.P. Cardoso, Carlos B. Duarte, Mário Grãos, Bruno Manadas

## Abstract

Hypoxic-ischemic encephalopathy (HIE) is one of the leading causes of child death worldwide. Most of the survivors develop various neurological diseases, such as cerebral palsy, seizures, and/or motor and behavioral problems. HIE is caused by an episode of perinatal asphyxia, which interrupts the blood supply to the brain. Due to its high energy demands, this interruption initiates glutamate excitotoxic pathways, leading to cell death. Umbilical cord mesenchymal stem cells (UC-MSCs) are gaining attention as a promising complement to the current clinical approach, based on therapeutic hypothermia, which has shown limited efficacy. Previous data have shown that priming MSCs under physiological culture conditions, namely soft platforms (3kPa) – mechanomodulated – or physiological oxygen levels (5% O_2_) – physioxia – leads to changes in the cellular proteome and their secretome. To evaluate how exposing MSCs’ to these culture conditions could impact their therapeutic potential, physiologically primed UC-MSCs or their secretome were added to an *in vitro* HIE model using cortical neurons primary cultures subjected to oxygen and glucose deprivation (OGD) insult. By comparing the neuronal proteome of sham, OGD insulted, and OGD-treated neurons, it was possible to identify proteins whose levels were restored in the presence of UC-MSCs or their secretome. Despite the different approaches that differentially altered UC-MSCs’ proteome and secretome, the effects converged on the re-establishment of the levels of proteins involved in translation mechanisms (such as the 40S and 60s ribosomal subunits), possibly stabilizing proteostasis, which is known to be essential for neuronal recovery. Interestingly, treatment with the secretome of UC-MSC modulated under physioxic conditions sustained part of the neuronal network integrity and modulated several mitochondrial proteins, including those proteins involved in ATP production. This suggests that the unique composition of the physioxia-modulated secretome may offer a therapeutical advantage in restoring essential cellular processes that help neurons maintain their function, compared to traditionally expanded UC-MSCs. These findings suggest that both the presence of UC-MSCs and their secretome alone can influence multiple targets and signaling pathways, collectively promoting neuronal survival following an OGD insult.

## Introduction

Birth asphyxia is a clinical condition that results from the lack of an adequate blood supply to the newborn in the perinatal period (1, 2). This event may affect several organs, but the brain is particularly vulnerable due to the high energy requirements and low energy storage (3, 4). Thus, this event often results in brain damage and can progress to hypoxic-ischemic encephalopathy (HIE) which, according to the World Health Organization, is one of the leading causes of child death and disability worldwide (5–8). In fact, more than half of the survivors die within the first years of life or develop neurological disorders such as cerebral palsy, seizures, motor, behavioral and/or cognitive issues (9–12).

Briefly, after the initial lack of energy resulting from asphyxia, there is a blood flow restoration (reperfusion) followed by a latent phase, which can last from 1 to 6-12 h (13). The first phase is predominantly characterized by an anaerobic metabolism, resulting in an increase in lactic acid production and a decrease in ATP production. This energy failure leads to an impairment of active membrane transport and, consequently, an increased concentration of intracellular sodium and calcium, culminating in permanent membrane depolarization with consequent neurotransmitter release (13, 14). Excitotoxicity is a common phenomenon observed in this stage, and it all commences with the accumulation of the neurotransmitter glutamate at the synaptic cleft (due to inefficient clearance by astrocytes) that will promote calcium entry, thus restarting the cycle, thereby amplifying the excitotoxicity process. Moreover, the electro-chemical and osmotic imbalance leads to water influx, cellular swelling, and consequently, cytotoxic edema (14, 15). Also, oxidative stress leads to the accumulation of reactive oxygen species, followed by lipid peroxidation (15). Together, these events initiate inflammatory cytokine secretion and trigger necrotic and apoptotic pathways. Although most neurons partially recover during the initial phase of reperfusion, many of these cells will eventually die hours or days later due to a delayed energy failure during the secondary phase, which can last up to three days. Besides excitotoxicity and cytotoxic edema, the secondary deterioration is characterized by a failure of mitochondrial activity, leading to redox misbalance and cytochrome-c release, activating apoptotic signaling cascades (16). These episodes are followed by the third phase, which comprises chronic inflammation, epigenetic modifications, brain repair and remodeling, and can last for months or even years (14, 17).

Presently, therapeutic hypothermia (TH) is the current standard of care for term newborns diagnosed with moderate to severe HIE. This procedure aims to reduce the newborn’s body temperature to 33.5°-35°C for 72 hours to decrease the cellular metabolic rate to its minimum (18–20). It has been reported that this treatment can attenuate microglia activation, suppressing the release of pro-inflammatory molecules and halting the intracellular cell death pathways (21, 22). However, its efficacy is far from ideal. In recent years, stem cells have emerged as a promising therapeutic approach in preclinical studies since they have been described to stimulate neural stem cell proliferation and differentiation, inhibit apoptotic pathways, and decrease astrogliosis and microglial activation, mainly through paracrine signaling (23–25). On the negative side, producing a single dose of stem cells requires an extensive in vitro expansion, which might compromise their stemness and, consequently, their therapeutic potential (26, 27). Our team demonstrated that priming umbilical cord-mesenchymal stem cells (UC-MSCs) with an environment that resembles their *in vivo* stiffness (3kPa) and oxygen levels (5%O_2_) is capable of modulating their proteome and secretome (28, 29).

The rescue potential of physiological primed UC-MSCs was accessed using an *in vitro* HIE model, i.e., immature primary cultures of cortical neurons subjected to an oxygen-glucose deprivation (OGD) insult, to mimic a perinatal asphyxia (PA) event (30). After the OGD insult, the neurons were treated with primed MSCs or their secretome and maintained in culture for sixteen hours. The therapeutic potential was evaluated by maintaining the neuronal network, and the proteome of treated and non-treated neurons was analyzed to unveil the alterations triggered by these approaches and identify potential therapeutic targets.

## Methods

### Cell culture

#### Preparation of primary cultures of embryonic rat cerebrocortical neurons

Primary cultures of cortical neurons were isolated as previously described (31). Briefly, cortices of E18-E19 rat embryos were washed with HBSS (5.36 mM KCl, 0.44 mM KH_2_PO_4_, 137 mM NaCl, 4.16 mM NaHCO_3_, 0.34 mM Na_2_HPO_4_·2H_2_O, 5 mM glucose, 1 mM sodium pyruvate, 10 mM HEPES and 0.001% phenol red, pH 7.2) and the tissue was treated with a 0.4 mg/mL trypsin solution (Gibco™) in HBSS, for 10 minutes at 37°C. The cortices were then washed twice with HBSS and mechanically dissociated with a glass pipette. Viable neurons were counted manually using a Trypan Blue solution (Gibco™) and plated directly onto multi-well plates (to obtain protein extracts) or on coverslips (for immunocytochemistry assay) previously coated with poly-D-lysine (0.1 mg/mL) with neuronal plating medium (MEM supplemented (Sigma-Aldrich), 0.6% glucose and 1 mM pyruvic acid, pH 7.2) at a cell density of 92.8 10^3^ cells/cm^2^. After 2-3 hours, the culture medium was exchanged to Neurobasal medium (Gibco™), supplemented with NeuroCult™ SM1 Neuronal Supplement (1:50) (STEM CELL), 0.5 mM L-glutamine (Gibco™) and 0.12 mg/mL gentamycin (Gibco™). Cells were maintained on a humidified incubator containing 95% air and 5 % CO_2_, at 37°C for 7 days. At day 3-4, one third of the medium was replaced by fresh Neurobasal medium supplemented with 30 μM 5-FDU (final concentration 10 μM), to halt the growth of glia cells.

#### UC-MSCs expansion and secretome preparation

MSCs were expanded as previously published (32, 33). Briefly, MSCs were isolated from the UC matrix (Wharton’s jelly) and expanded with proliferation medium (Minimum Essential Medium-α (MEM-α) (Gibco™) supplemented with 5% (v/v) fibrinogen depleted human platelet lysate (HPL) (UltraGRO™, Helios) and antibiotics: 100 U/mL of Penicillin, 100 μg/mL Streptomycin and 2.5 μg/mL Amphotericin B or Antibiotic-Antimycotic (all from Gibco™). Upon confluence, cells were trypsinized by adding 0.05% Trypsin-EDTA solution (Gibco™) for five minutes at 37°C, homogenized, centrifuged, and re-seeded until P3. At P4, UC-MSCs were plated at 10,000/cm^2^ and kept in a humidified incubator with 5% CO_2_/atmospheric oxygen levels in tissue culture polystyrene Petri dishes (2-3 GPa) – standard conditions - or in the same incubator but on 3kPa PDMS coated dishes – mechanomodulation. For physioxia set up, cells were seeded in tissue culture polystyrene (TCP) Petri dishes but moved to a humidified oxygen level-controlled environment (5%O_2_/5%CO_2_). After 24h, cells were washed twice with PBS 1×, and resuspended in serum-free Neurobasal medium (supplemented with Antibiotic-Antimycotic (Gibco™). The cell secretome was collected after 24h of incubation as previously described (34). In total, eleven donors were used (six for the secretome and five for the UC-MSCs co-culture treatment).

The secretome was harvested and centrifuged at 290×g, for 5 min at 4°C to remove cell debris. Then, the secretome also known as conditioned medium (CM) was transferred to a low molecular weight cut-off (5kDa Vivaspin^®^ 20, Sartorius) and centrifuged at 3000×g, at 4°C. The final concentration (v/v) for each UC-MSCs donor was assessed. Ready to use aliquots were stored at −20°C to prevent freeze/thaw cycles.

#### Oxygen glucose deprivation stimuli and treatment assay

At DIV7, cortical neurons were incubated with a glucose-free medium (10 mM HEPES, 116 mM NaCl, 5.4 mM KCl, 0.8 mM MgSO_4_, 1 mM NaH_2_PO_4_, 25 mM NaHCO_3_, 1.8 mM CaCl_2_, 25 mM sucrose, pH 7.3), for five hours in an InvivO₂® 400 Physoxia Workstation (Baker) at 0%O_2_/5%CO_2_. The medium of control neurons (sham) was replaced by a similar medium but containing 25 mM glucose instead of sucrose. These cells were maintained for the same period in a humidified incubator with 5% CO_2_/atmospheric oxygen levels.

For the UC-MSCs secretome treatment assay, after the insult, the CM was diluted 10× in the neurons’ conditioned medium, and the cells were allowed to recover for 16 hours. For the MSCs rescue assay, primed cells were seeded in coverslips at 10,000/cm2 and placed on top of the neurons. Glass beads (0.5 mm) were glued to the surface to guarantee a tight space between both cultures without direct touch.

For proteomic analysis or Western Blot, neuronal protein extracts were collected on ice by the addition of a Lysis buffer (120 mM Tris-HCl pH 6.8, 4% (w/v) Sodium dodecyl sulfate (SDS) and 20% (v/v) glycerol). Then, sample buffer 6× (350 mM Tris-HCl pH 6.8, 30% (v/v) glycerol, 10% (w/v) SDS, 0.93% (w/v) dithiothreitol (DTT)) was added to the sample to obtain a final concentration of 1×. Samples were denatured at 95°C for five minutes and sonicated. Protein was quantified using Pierce^TM^ 660nm Protein Assay (Thermo Fischer Scientific) according to the manufacturer’s instructions. For cell imaging, the immunocytochemistry protocol was followed.

#### Glutamate excitotoxicity assay

This procedure was performed as previously described (35). Briefly, at DIV7, cortical neurons were incubated with 125 μM glutamate prepared in Neurobasal medium for 20 min, which was then replaced by neuronal conditioned medium, and a post-incubation was performed for six hours. Samples were collected for Western Blot or immunocytochemistry analysis. For propidium iodide staining (PI), neurons were incubated with a 2 μg/mL PI (Invitrogen) solution in PBS, for 10 minutes at 37°C. Cells were then fixed with 4% (w/v) paraformaldehyde (Acros) for twenty minutes at room temperature.

#### Immunocytochemistry (ICC) assay

Cultured neurons were fixed with 4% (w/v) paraformaldehyde (Acros) for twenty minutes at room temperature. Next, cells were washed twice with PBS and permeabilized by the addition of 0.5% Triton-x100 (Acros), for ten minutes at 4°C. Fixed neurons were blocked with 10% bovine serum albumin (BSA) in PBS for one hour at 37°C, to prevent unspecific binding. Then, coverslips were incubated with the following primary antibodies (PDS95 1:200 #6G6-1C9, Invitrogen; VGlut1 1:100 #482400, Invitrogen; NMDAR1 (GluN1) 1:400 #AGC-001, Alomone; MAP-2 1:200 #sc-74421, Santa Cruz; Cleaved PARP (Asp14) 1:800 #94885 Cell Signaling Technology, Inc.) diluted in 3% BSA in PBS and left overnight at 4°C. On the next day, samples were washed three times with PBS and incubated with the secondary antibodies (goat anti-rabbit Alexa fluor 488 1:200 #A-11008, Invitrogen; goat anti-mouse Alexa fluor 568 1:200 #A-11004, Invitrogen) and Hoechst 33342 (1:1000) (Thermo Scientific) for one hour at room temperature. Coverslips were washed and mounted with fluorescence mounting medium (Dako). Images were acquired using a Zeiss Axiovert 200 M fluorescence microscope and AxioVision release 4.8 software (Zeiss). Acquisition settings were kept under the same conditions.

Images were analyzed on Cell Profiler 4.1.3 (36) and the total number of cells was counted using the nucleus (stained with Hoechst 33342) using the IdentifyPrimaryObjects module. For the glutamate excitotoxicity assay, PI staining integrated intensity in the nuclear area (using MeasureObjectIntensity) was used to calculate the total number of positive cells (using the ClassifyObjects module). The mean of all twelve images acquired per experimental condition was calculated for each sample. For OGD experiments, MAP2 and cleaved PARP2 integrated intensity were calculated on the cellular body region (using IdentifySecondaryObjects and MeasureObjectIntensity modules) to calculate the total number of MAP2 and cleavedPARP positive cells (using ClassifyObjects module).

#### Western blot (WB)

Cellular extracts were prepared as described above. Protein extracts (12.5 µg) were loaded and resolved by using SDS-PAGE with 10% (w/v) acrylamide–bisacrylamide gels (BioRad). Next, proteins were transferred to a low fluorescence polyvinylidene fluoride (PVDF) membrane for 30 min at a constant voltage of 25 V (up to 1 A), using a Trans-Blot Turbo Transfer System (BioRad). Membranes and solutions used were provided with the transfer kit (TBT RTA TRANSFER KIT, BioRad). To confirm protein transfer, membranes were stained with Ponceau S staining Solution (Thermo Scientific). After removing the stain, membranes were blocked for 1 hour with 5% (w/v) skimmed milk powder in TBS-Tween 20 (Bio-Rad) (TBS-T) [0.1% (v/v)]. Then, membranes were incubated with the following primary antibodies (MAP-2 1:1000 #sc-74421, Santa Cruz; Cleaved PARP (Asp14) 1:3000 #94885, Cell Signaling) overnight at 4°C followed by 1 hour at room temperature, with agitation. Following this incubation, membranes were washed three times (5 minutes, with agitation) with TBS-T and incubated with the respective secondary antibody (anti-mouse 1:10,000 (115-055-003)) conjugated with alkaline phosphatase for 1 hour, at room temperature. Finally, membranes were incubated with enhanced chemifluorescence substrates (ECF – GE Healthcare) for 5 minutes and imaged using a VWR Imager (VWR). For the cleaved PARP antibody, the secondary antibody Cy®5 ECL Plex 1:2000 (#PA45011, Cytiva) was used. To determine the total amount of protein in each lane, membranes were stained with SERVA purple (SERVA electrophoresis, Enzo) following the manufacturer’s guidelines. Images were analyzed with Image Lab software, version 5.1 (BioRad).

#### Measurement of oxygen levels and oxygen consumption rate (OCR)

The OCR and the levels of oxygen dissolved in the culture media were continuously monitored on a MW96, using Resipher (Lucid Scientific), on each well during the OGD insult or for the following thirty-six hours. Also, after the glutamate excitotoxicity assay, these parameters were monitored for eighteen hours. Data was acquired in duplicates or triplicates for each experimental condition.

#### Proteomic analysis

A protein standard MBP-GFP was added to each protein extract (37). Proteins were alkylated by the addition of a 40% acrylamide solution, resolved through a short-GeLC approach, and stained with Coomassie Brilliant Blue G-250 (38, 39). Pools of each condition were prepared for protein identification. Concerning protein quantification, each sample was loaded individually. Gel lanes were cut and destained, and proteins were digested overnight using trypsin. Finally, the peptides were extracted through the sequential addition of solutions with an increased acetonitrile concentration (30%, 50% and 98%) and 1% formic acid, dried by vacuum centrifugation at 60°C, and stored at −20°C.

Samples were resuspended on 2% acetonitrile and 0.1% formic acid solution and analyzed on a NanoLC™ 425 System (Eksigent) coupled to a Triple TOF™ 6600 mass spectrometer (Sciex) with an electrospray ionization source (DuoSpray™ Ion Source from Sciex). Peptides were separated on a Triart C18 Capillary Column 1/32” (12 nm, 3 μm, 150 mm × 0.3 mm, YMC, Dinslaken, Germany) and using a Triart C18 Capillary Guard Column (0.5 mm × 5 mm, 3 μm, 12 nm, YMC) at 50 °C. The flow rate was set to 5 µL/min and mobile phases A and B were 5% DMSO plus 0.1% formic acid in water and 5% DMSO plus 0.1% formic acid in acetonitrile, respectively. The LC program was performed as follows: 5 – 30% of B (0 - 50 min), 30 – 98% of B (50 – 52 min), 98% of B (52-54 min), 98 - 5% of B (54 – 56 min), and 5% of B (56 – 65 min). The ionization source was operated in the positive mode set to an ion spray voltage of 5500 V, 25 psi for nebulizer gas 1 (GS1), 10 psi for nebulizer gas 2 (GS2), 25 psi for the curtain gas (CUR), and source temperature (TEM) at 100°C. For Data Dependent Acquisition (DDA) experiments, the mass spectrometer was set to scan full spectra (m/z 350–1500) for 250 ms, followed by up to 100 MS/MS scans (m/z 100–1500) with a 30 ms accumulation time, resulting in a cycle time of 3.3s. Candidate ions with a charge state between + 1 and + 5 and counts above a minimum threshold of 100 counts per second were isolated for fragmentation, and one MS/MS spectrum was collected before adding those ions to the exclusion list for 15 s (mass spectrometer operated by Analyst® TF 1.8.1, Sciex®). For Data Independent Acquisition (DIA) (or Sequential Window Acquisition of all Theoretical Mass Spectra (SWATH)) experiments, the mass spectrometer was operated in a looped product ion mode and specifically tuned to a set of 168 overlapping windows, covering the precursor mass range of 350–2250 m/z. A 50 ms survey scan (m/z 350–2250) was acquired at the beginning of each cycle, and SWATH-MS/MS spectra were collected from 100 to 2250 m/z for 19 ms, resulting in a cycle time of 3.3s.

Each fraction of pooled samples was acquired individually in DDA mode. ProteinPilot™ software was used to generate the ion library of the precursor masses and fragment ions (v5.0, Sciex), combining all files from the pools. The data was searched against the reviewed Rat (Swissprot) database (downloaded on 22^nd^ March 2023), using the following settings: cysteine alkylation by acrylamide, digestion by trypsin, and gel-based ID factors. The quality of the identifications was assessed by an independent False Discovery Rate (FDR) analysis, which was performed using the target-decoy approach provided by the software.

For protein relative quantification, samples were acquired in DIA mode/SWATH. Data was processed using the SWATH™ plug-in for PeakView™ (v2.0.01, Sciex®), considering up to 15 peptides per protein, and up to 5 transitions per peptide. Only peptides with an FDR below 1% in at least 1/3 of the samples per group per experimental condition were quantified. Data was normalized to the total intensity. The MS data have been deposited to the ProteomeXchange Consortium via the PRIDE partner repository with the dataset identifier PXD057623 (40, 41).

#### Data analysis

Statistical analysis was performed on MetaboAnalyst 6.0 (42). For proteomic screenings, differentially expressed proteins (DEPs) were selected through multivariate (Partial least squares-discriminant analysis (PLS-DA)) or univariate (t-test, Mann-Whitney or Kruskal-Wallis tests) analysis if they had a VIP>1 and/or p<0.05 and fold change (FC) above 1.5 or below 0.67 (|Log_1.5_FC|>1). These proteins were then subjected to further analyses. Gene Ontology enrichment analysis was performed on Funrich (43). Reactome database was used to map pathways (44), Venn diagrams were generated on DeepVenn (45), and violin plots were created on GraphPad Prism 9. Clustering analysis was performed on R studio, using Mfuzz package (46)For WB or ICC, the statistical analysis was performed on GraphPad Prism 9, using the non-parametric Kolmogorov-Smirnov or Mann-Whitney tests, respectively.

## Results

### Primary cortical neurons lose their neuronal network after the OGD insult

To model an immature brain similar to that of a newborn, primary cortical neurons were isolated and cultured for seven days to allow maturation (30, 47). Considering that HIE pathophysiology relies on excitotoxicity pathways dependent on the activation of glutamate receptors, the expression of glutamatergic markers as GluN1 and VGlut1 were confirmed by immunocytochemistry, as well as the postsynaptic marker PSD95 (Supplementary Figure 1A). Moreover, to assess whether these are functional receptors, we investigated the effect of glutamate added to the culture (125 µM for 20 minutes) on neuronal viability. After six hours, there was a loss of MAP2 expression (a dendritic marker) (Figure 1A,) and an increased number of PI positive cells (Figure 1B, Supplementary Figure 1B), confirming the loss of the neuronal network and plasma membrane integrity, respectively.

**Figure 1.**
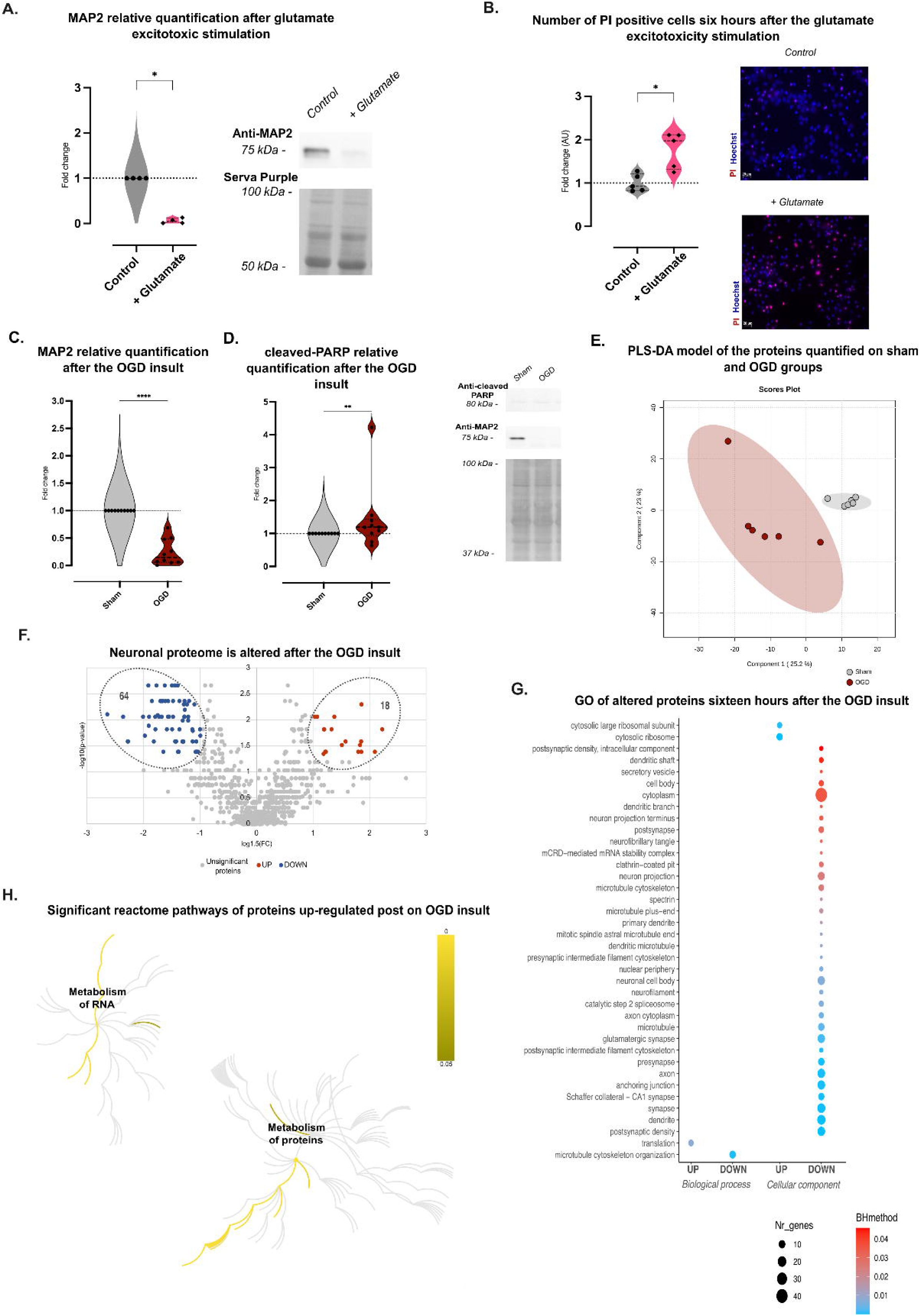
Neuronal network integrity and protein metabolism are compromised after the OGD insult. (A) Neurons were subjected to an excitotoxicity glutamate stimulation. After six hours, there was a decrease in total levels of MAP2 levels (*p<0.05, Kolmogorov-Smirnov) and (B) an increase in the total number of positive PI cells (*p<0.05, Mann-Whitney). Moreover, neurons were subjected to a five-hour OGD insult and left in culture for an additional sixteen hours. (C) There was a significant decrease in the total levels of MAP2 and (D) a slight increase in the total levels of cleaved-PARP (****p<0.0001; **p<0.001, Kolmogorov-Smirnov). Cellular extracts were also analyzed by proteomics. (E) The total number of proteins quantified were used to generate a PLS-DA model, where 308 proteins were found to have a VIP>1. (F) Several proteins were found to be differentially expressed in neurons incubated under OGD when compared to sham neurons (Mann-Whitney, p<0.05; upregulated proteins are represented in red, and downregulated proteins are highlighted in blue). (G) Go analysis of the proteins showing significant alterations in their abundance (VIP>1, p<0.05, or both; |Log1.5FC|>1)). (H) Reactome pathways of proteins whose levels significantly increase after the insult are highlighted (Supplementary Table 1). No significant pathways were found to be associated with downregulated proteins. Abbreviations: Mechano (mechano-modulation); OGD (oxygen-glucose deprivation).

After confirming the glutamatergic phenotype, neurons were subjected to an OGD insult to evaluate their ability to respond to it. Oxygen levels in the culture media reached near zero values during the insult, which were reestablished to atmospheric levels immediately after the insult (Supplementary Figure 1C). Sixteen hours later, no differences were observed in HIF1α localization (Supplementary Figure 1D). At this time point, it was possible to observe a decrease in the total levels of MAP2 and a slight increase in the total levels of cleaved PARP, which is suggestive that the neuronal network integrity is compromised and confirms that this cellular model is responsive to the OGD insult (Figure 1C, D, respectively).

Apart from evaluating the neuronal network integrity, a proteomic screening was performed to further explore the impact of the insult in this *in vitro* HIE model. The cellular proteome of sham and OGD neurons was collected and analyzed sixteen hours after the five-hour insult. Both supervised and unsupervised analyses show a separation between the two conditions (Figure 1E, Supplementary Figure 1E). Additionally, using a univariate approach, eighteen proteins were found to be upregulated after the OGD insult, while sixty-four were found to be decreased (Figure 1F).

Moreover, the GO enrichment analysis of DEPs (VIP>1, p<0.05, or both and |Log_1.5_FC|>1) showed that the decreased proteins are associated with the microtubule cytoskeleton (Figure 1G), confirming the loss of the neuronal network described above. Furthermore, data indicate that proteins related with translation processes are upregulated with this insult, as well as those involved in RNA metabolism (Figure 1H; Supplementary Table 1). Together, these results demonstrated that, after the OGD insult, this HIE model shows defects in the neuronal network and changes in the neuronal proteome that may impact neuronal survival.

### Physiologically modulated MSCs lead to a modulation of neuronal translation processes after the OGD insult

Previous reports indicated that priming UC-MSCs on soft substrates or low oxygen levels to resemble a more physiological environment altered their proteome (28). Therefore, it was hypothesized that physiologically modulated MSCs could store “physiological memory” and trigger different pathways when in contact with the neurons. To test this, after the OGD insult, standard, mechano-modulated, or physioxia-modulated UC-MSCs were added to the culture immediately after the OGD insult, and the two populations of cells were allowed to communicate. Neuronal extracts were collected after 16 hours of recovery. Despite the decrease of the total levels of MAP-2 in OGD-treated and non-treated neurons, compared to sham, it appears that the levels of cleaved-PARP are slightly lower in the presence of physioxia-modulated UC-MSCs when compared to the other treatments (Figure 2A, B, respectively). So, to explore the differences triggered by modulated MSCs, a proteomic screening was performed to investigate the impact of the presence of conventional, mechanomodulated, or physioxia-modulated MSCs cocultured with post-OGD neurons. A clustering analysis was performed, and two clusters were selected per experimental condition: the ones that identified proteins whose levels would be restored to control levels after the addition of MSCs to the culture (Supplementary Figure 2A-C). Then, these proteins were selected for further analysis. These unsupervised approaches (Supplementary Figure 3A-C) showed a closer relationship between neurons maintained under sham conditions and those incubated under experimental conditions that provided neuroprotection. In addition, the supervised method (PLS-DA) could distinguish in the latter group the neurons incubated in the presence of mechano-modulated and physioxia-modulated MSCs, suggesting that their differential background can result in a different message transmitted to the post-OGD neurons (Figure 2C-E, respectively). To further understand which pathways were altered by the modulated MSCs, the proteins clustered in the selected clusters were analyzed using the Reactome Database (Figure 2F; Supplementary Table 2). Despite being different processes, a higher number of pathways is associated to proteins whose levels increase after the insult but are restored after the presence of UC-MSCs. Conventional and modulated UC-MSCs appear to modify proteins associated with cell responses to stimuli (including hypoxia), while only mechano-modulated and physioxia-modulated UC-MSCs can interfere with the translation processes, which might also justify the differences encountered on cleaved-PARP levels. Particularly, mechanomodulated UC-MSCs seemed to impact more pathways, as cell metabolism and vesicle-mediated transport, that were not shared by the other treatment conditions.

**Figure 2.**
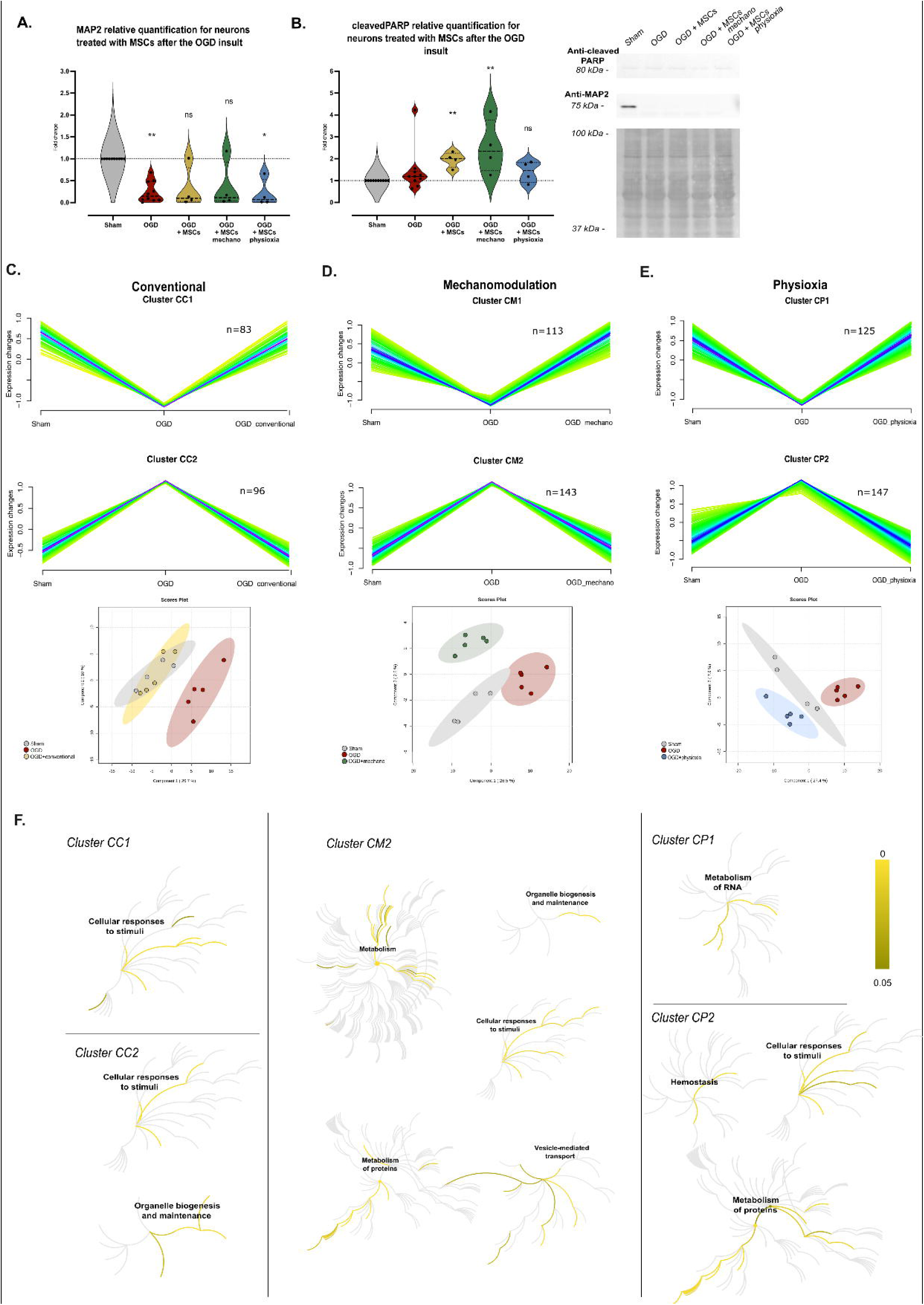
Co-culture of physiologically primed MSCs and post-OGD neurons differentially modulate the neuronal proteome during treatment. Neurons were subjected to an OGD insult. Conventional or physiologically modulated UC-MSCs were added to the culture and left in culture for sixteen hours when the cellular extracts were collected. The levels of MAP2 and cleaved-PARP were quantified by WB (A, B, respectively) (Kruskall-Wallis test; *p<0.05; **p<0.01). Also, the proteomics screening allowed the identification of 1852 proteins, from which 1077 were confidently quantified. An MFuzz clustering analysis was performed to identify proteins whose profiles were similar to the sham in the presence of conventional (C), mechano-modulated (D), or physioxia-modulated (E) UC-MSCs (blue represents higher membership values; yellow represents lower membership values). These proteins were then used to generate a generate a PLS-DA model for each experimental condition (bottom) (sham: grey; OGD: red; standard UC-MSCs: yellow; mechano-modulated UC-MSCs: dark green; physioxia-modulated UC-MSCs: dark blue). (F): Reactome pathways of the clustered proteins are represented. Abbreviations: Mechano (mechano-modulation); OGD (oxygen-glucose deprivation); CC (Cells conventional); CM (Cells mechanomodulated); CP (Cells physioxia-modulated).

To identify potential therapeutic response markers in the neuronal proteome, a non-parametric univariate analysis (Mann-Whitney) was performed to select proteins that showed statistically significant alterations in their expression levels when neurons maintained under sham conditions vs OGD by a mechanism sensitive to the presence of UC-MSCs (Supplementary Tables 3-5). Proteins that matched the univariate and the selection done via the clustering analysis are listed in Table 1, and only four were shared by the three types of treatment: Heat shock protein HSP 90-alpha and 70kDa protein 4, Ras-related GTP-binding protein B and Dipeptidyl peptidase 33 (Figure 3A, Supplementary Figure 3D-F). From this list, six were DEPs previously identified when comparing sham and OGD, and the levels were restored in neurons co-cultured with physiologically primed UC-MSCs (Figure 3B). Moreover, comparing the processes found to be altered in the neuronal proteome after the OGD insult (Figure 1H) but restored by the treatment with MSCs (Figure 2F), nine processes were found to be uniquely modified by physiologically modulated MSCs but not conventional UC-MSCs (Figure 3C). These pathways were mainly related to translation mechanisms and L13a-mediated translational silencing of Ceruloplasmin expression. This evidence suggests that the differences found between the modulation of standard and physiologically modulated UC-MSCs are suggestive that the crosstalk between UC-MSCs and neurons depends on the previous background of UC-MSCs culture condition.

**Figure 3.**
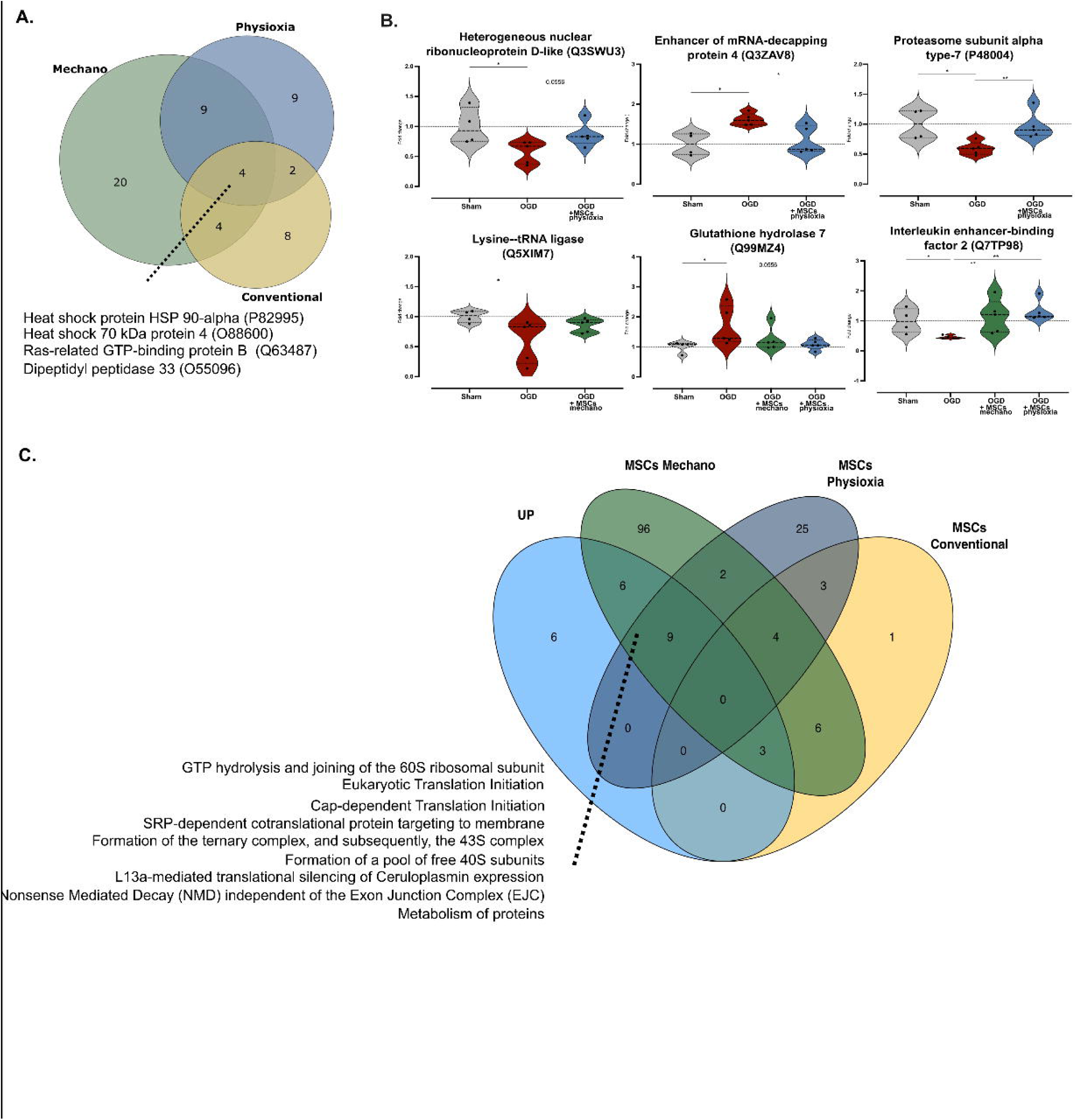
Co-culture with physiological modulated UC-MSCs modify neuronal translation processes. (A) After the OGD insult, neurons were left in culture with conventional or physiologically modulated UC-MSCs. By comparing the proteins listed in Table 1, only four proteins were shared between the three experimental conditions. (B) DEPs between sham and OGD (|Log_1.5_FC|>1) but that lost their significance in the presence of physiologically-modulated UC-MSCs are represented (Mann Whitney; *p<0.05, **p<0.01). (C) Comparison of Reactome pathways associated with upregulated proteins comparing sham and OGD (Figure 1H) and those whose levels were restored in the presence of conventional or physiologically modulated MSCs. Abbreviations: Mechano-modulation (mechano).

**Table 1.**
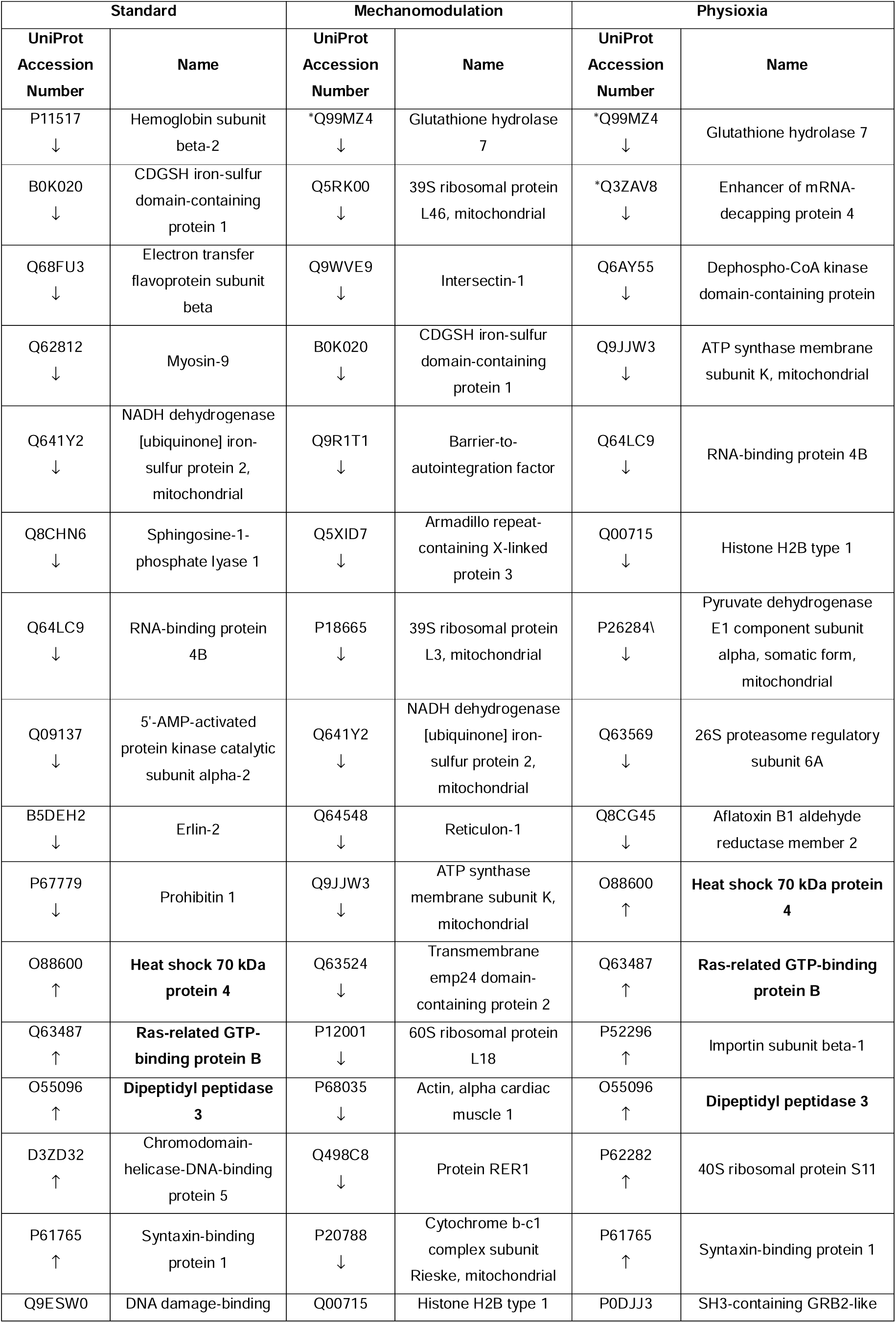
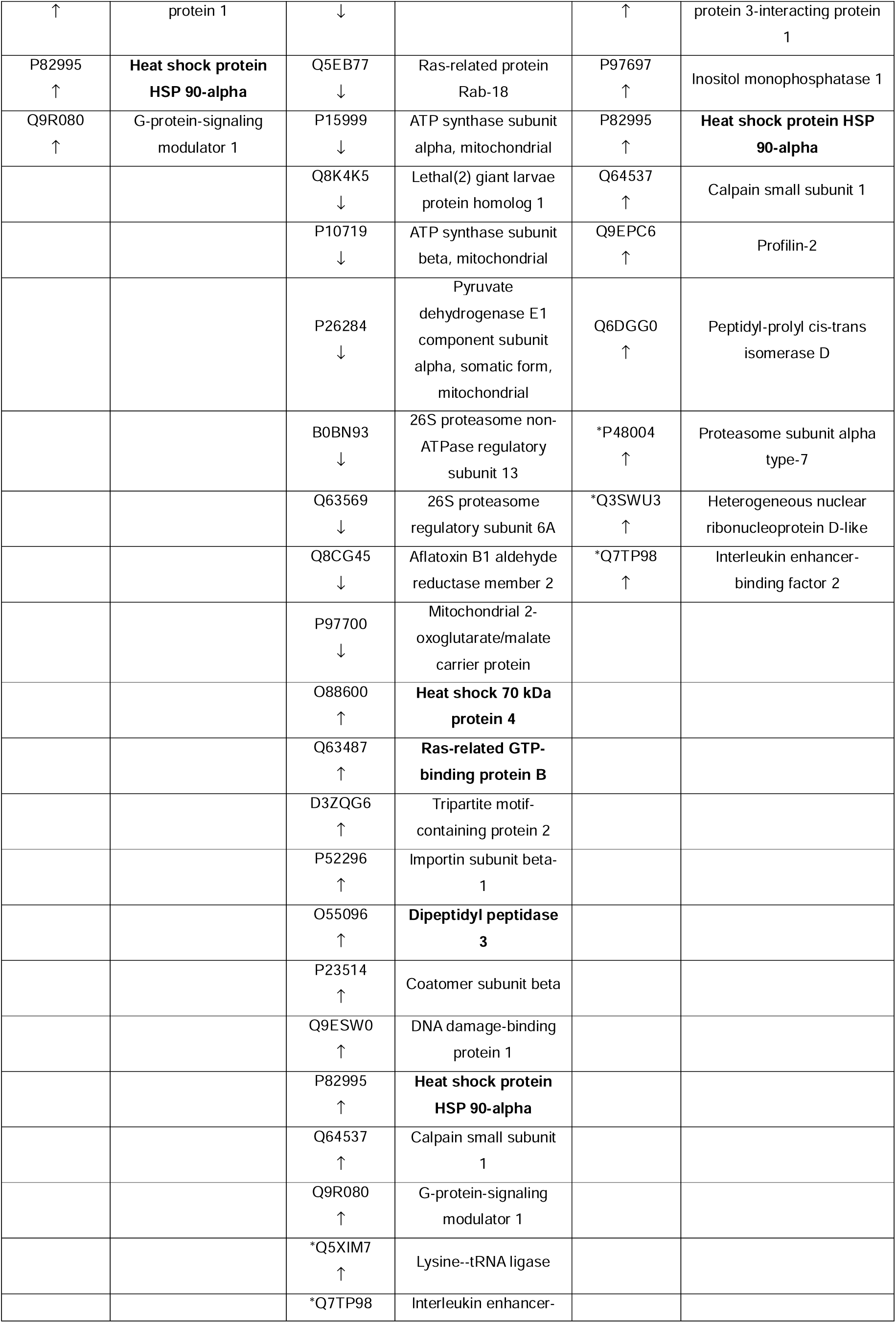
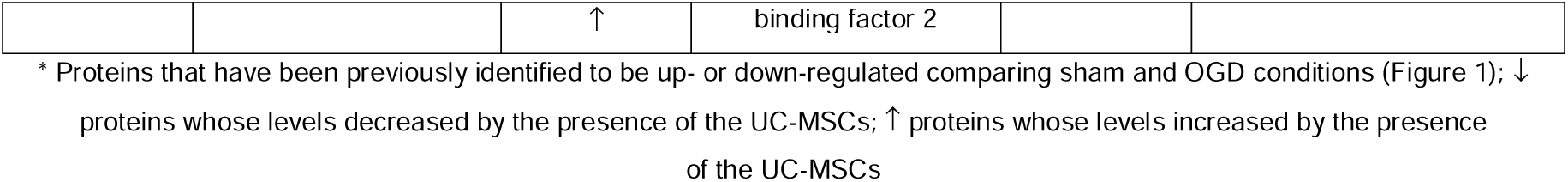
List of the proteins differentially modulated by the presence of conventional mechano-modulated or physioxia-modulated UC-MSCs.

### Physioxia-modulated UC-MSCs’ secretome prevents the complete loss of the neuronal network

Our previous study (29) reported the unique composition of UC-MSCs’ secretome after priming. Therefore, the impact of adding physiologically modulated UC-MSC’ secretome to neuronal culture after an OGD was also investigated. To do that, conventional expanded or physiologically-modulated UC-MSCs’ secretome (either at 3kPa substrates (mechano) or 5%O_2_ (physioxia)) was added to both sham and OGD neurons after the insult and left to post-incubate for sixteen hours. At this time point, no differences were found in the total levels of MAP2 or cleaved-PARP in neurons treated with UC-MSCs secretome (Supplementary Figure 4A, B). Still, the total number of MAP2-positive cells did not significantly decrease in the presence of physioxia-modulated secretome and the total number of cells (Figure 4A-C). However, the number of positive cleaved-PARP neurons did not differ comparing treated or non-treated neurons (Supplementary Figure 4C). Together, these data suggest that the treatment with physioxia-modulated secretome leads to the maintenance of the neuronal cytoskeleton and, consequently, the neuronal network.

**Figure 4.**
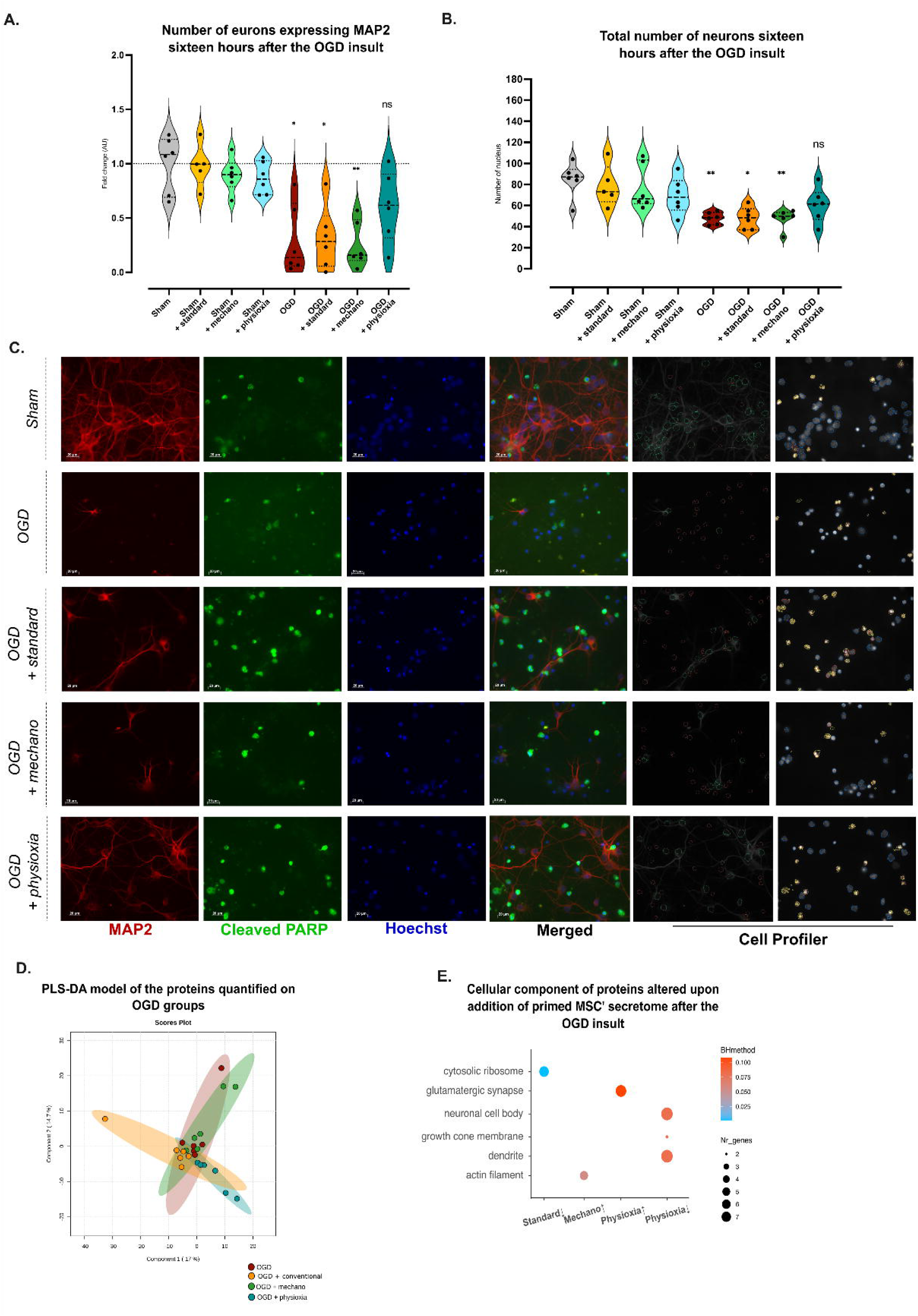
Secretome of physioxia-modulated UC-MSCs reduces the loss of the neuronal cytoskeleton integrity. Neurons were subjected to a five-hour OGD insult and then incubated with CM obtained from UC-MSCs under conventional culture conditions or in low stiffness environments (3kPa) – mechano-modulation – or physioxia-modulated (5%O_2_). The total number of positive MAP2 neurons and the total number of cells were assessed by ICC. (A and B, respectively) (*p<0.05, Mann-Whitney). (C) Representative images. On the right, CellProfiler classification settings are highlighted: positive MAP2 neurons (green), negative MAP2 neurons (orange), cleaved PARP (yellow), positive cleaved PARP cells (red), and negative cleaved PARP cells (blue). Proteins quantified on OGD treated and non-treated neurons were used to generate a PLS-DA model (D). Also, a GO analysis was performed using DEP comparing treated and non-treated OGD neurons (E). Abbreviations: Mechano (mechano-modulation); OGD (oxygen-glucose deprivation).

To further explore the alterations on the proteome triggered by UC-MSCs’ secretome, the proteome of OGD neurons treated with UC-MSCs secretome was compared to non-treated neurons. Comparing the total proteome of cerebrocortical neurons subjected to OGD followed by incubation with control conditioned medium with the proteome of neurons incubated with the physioxia-modulated secretome after OGD (represented in blue) revealed a more distant relationship on the hierarchical clustering and a tendency for PLS-DA scores plot separation (Figure 4D, Supplementary Figure 5A). This is in accordance with imaging data since the total number of MAP2-positive neurons and cells did not significantly change, compared to sham, when OGD neurons were left to post-incubate with this secretome. Moreover, physioxia-modulated secretome appears to differentially modulate a higher number of proteins in treated neurons, as well as significant ontologies, that include proteins involved in glutamatergic synapses and the maintenance of cell polarity (Figure 4E, Supplementary Figure 5B and C). The three treatments commonly upregulated only five proteins (Supplementary Figure 5D). Collectively, these results indicate that the secretome provident from physioxia-modulated UC-MSCs can prevent the loss of cytoskeleton integrity in neurons subjected to OGD, a process known to be pivotal in regulating proper neuronal function, most likely through translational regulation of the neuronal proteome.

Considering the impact of the presence of conventional or modulated secretome in the culture of sham neurons, no differences were found in the total number of MAP2 or cleaved PARP cells, as well as the total number of cells (Figure 4A and B; Supplementary Figure 4C and D). Moreover, the proteomic analysis shows no separation on the PLS-DA scores plot (Supplementary Figure 5E), consistent with the smaller number of proteins altered in the presence of the secretome (Supplementary Figure 5F and G).

A clustering analysis was performed to identify which protein classes might be involved in favoring the maintenance of the neuronal network, similar to what was described in the previous section. Proteins showing changes in neurons subject to OGD and whose levels were restored after adding UC-MCS’ secretomes were selected for each experimental condition (Figure 5A-C). Selected proteins of the two clusters were used on non-supervised and supervised multivariate analysis (Figure 5A-C, Supplementary Figure 7A-C). Although the unsupervised analysis (PCA) did not separate the groups, for all supervised analyses (PLS-DA models), there was an overlap between the results obtained in neurons maintained in sham conditions *vs.* treated neurons and a closer relationship between these two conditions (hierarchical clustering), suggesting that these proteins may be involved in reestablishing cellular mechanisms that might be responsible for promoting neuronal survival.

**Figure 5.**
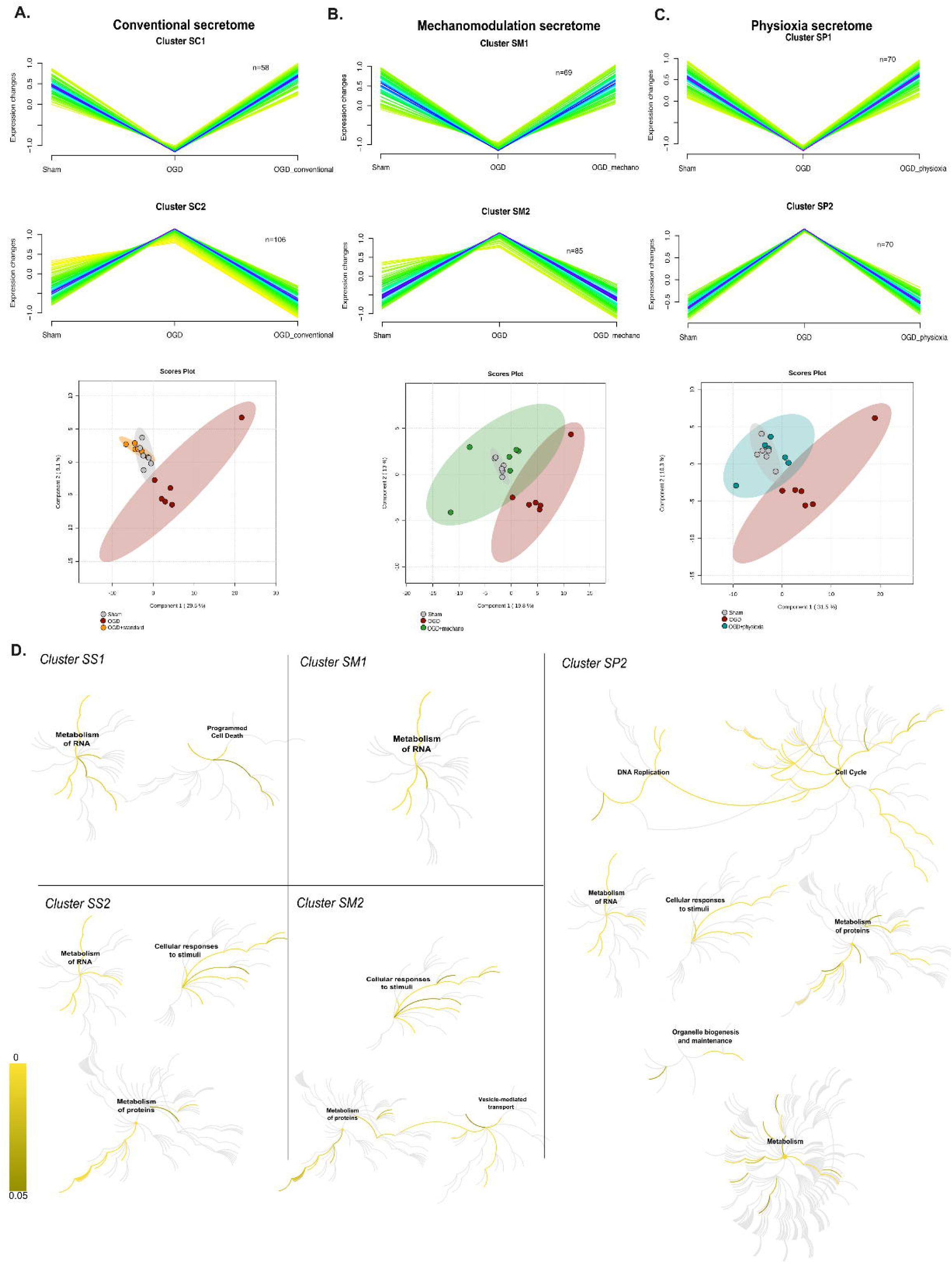
*UC-MSCs secretome* modulates *protein metabolism in post-OGD neurons.* After the insult, conventional or physiologically-modulated UC-MSCs CM (either at 3kPa substrates (mechano) or 5%O_2_ (physioxia)) was added to OGD neurons and left in culture for an additional sixteen hours, when the cellular extract was collected for proteome analysis. Of the 1152 proteins identified, 650 were quantified in all experimental conditions. The neuronal proteome was subjected to MFuzz clustering analysis to identify proteins whose levels were altered after the insult but restored after the addition of conventional (A), mechano-modulated (B), or physioxia-modulated (C) UC-MSCs secretome. PLS-DA scores plots were generated for each experimental condition using the proteins belonging to the selected clusters (bottom) (sham: grey; OGD: red; standard UC-MSCs: orange; mechano-modulated UC-MSCs: green; physioxia-modulated UC-MSCs: blue). (E): Reactome pathways of the proteins belonging to the selected clusters (Supplementary Table 8 for details). Abbreviations: Mechano (mechano-modulation); OGD (oxygen-glucose deprivation). Abbreviations: Mechano (Mechanomodulation); SC (Secretome conventional); SM (Secretome mechanomodulated); SP (Secretome physioxia-modulated).

The role of clustered proteins was then investigated using the Reactome database (Supplementary Table 8). Proteins whose levels decreased after the stimuli but were restored by the addition of the conventional or mechanomodulated secretome are mainly associated with the metabolism of RNA, including rRNA processing and nonsense-mediated decay processes (Figure 5D). However, a higher number of pathways displaying significant changes was associated with proteins whose levels decreased after adding the different MSCs’ secretomes. While all types of secretome appear to reduce the levels of proteins associated with cellular response to stress (including hypoxia) and the metabolism of proteins, including translation processes, physioxia-secretome interfere with other cellular processes, including metabolism (such as amino acids metabolism and nucleotide synthesis), cell cycle (including mitosis phases) and organelle biogenesis and maintenance, namely mitochondria (Figure 5D, Supplementary Table 8).

Furthermore, a cellular component (GO) analysis of clustered proteins revealed that proteins grouped on clusters SC2, SM2, and SP2 belong to cytosolic ribosomes and the proteasome complex ontologies (Supplementary Figure 7D). Interestingly, the addition of physioxia-modulated secretome modifies ontologies that were not shared with the other treatments, such as mitochondrial-proton transporting ATP synthase complex (OCR is not altered (Supplementary Figure 7E) and box H/ACA snoRNP complex were downregulated (for cluster SP2), and proteins associated with neuronal projections (for cluster SP1). These data suggests that the unique modulation of these processes might be supporting the maintenance of the cytoskeleton integrity, as described above (Figure 4).

To explore which neuronal pathways are being modulated by the different UC-MSCs secretome, the Reactome pathways of proteins whose levels decreased in the presence of UC-MSCs secretomes (cluster SC2, SM2 and SP2) were compared to the pathways associated with up-regulated proteins sixteen hours after the OGD insult (Figure 1H). The expression of proteins associated to translation processes, including ribosomal subunits and L13a-mediated translational silencing of Ceruloplasmin expression were altered by all types of secretome, while several other processes, as the RNA metabolism, including rRNA processing, were shared by standard and physioxia-modulated UC-MSCs derived secretome (Figure 6A). Interestingly, pathways associated with mitochondria were only altered by the physioxia-modulated stem cells’ secretome, suggesting that the energy support provided by these secretomes might be the key to preventing neuronal cytoskeleton integrity. Together, these data suggest that the unique composition of the secretomes (29) leads to the modulation of different proteins, most likely affecting different intracellular pathways.

**Figure 6.**
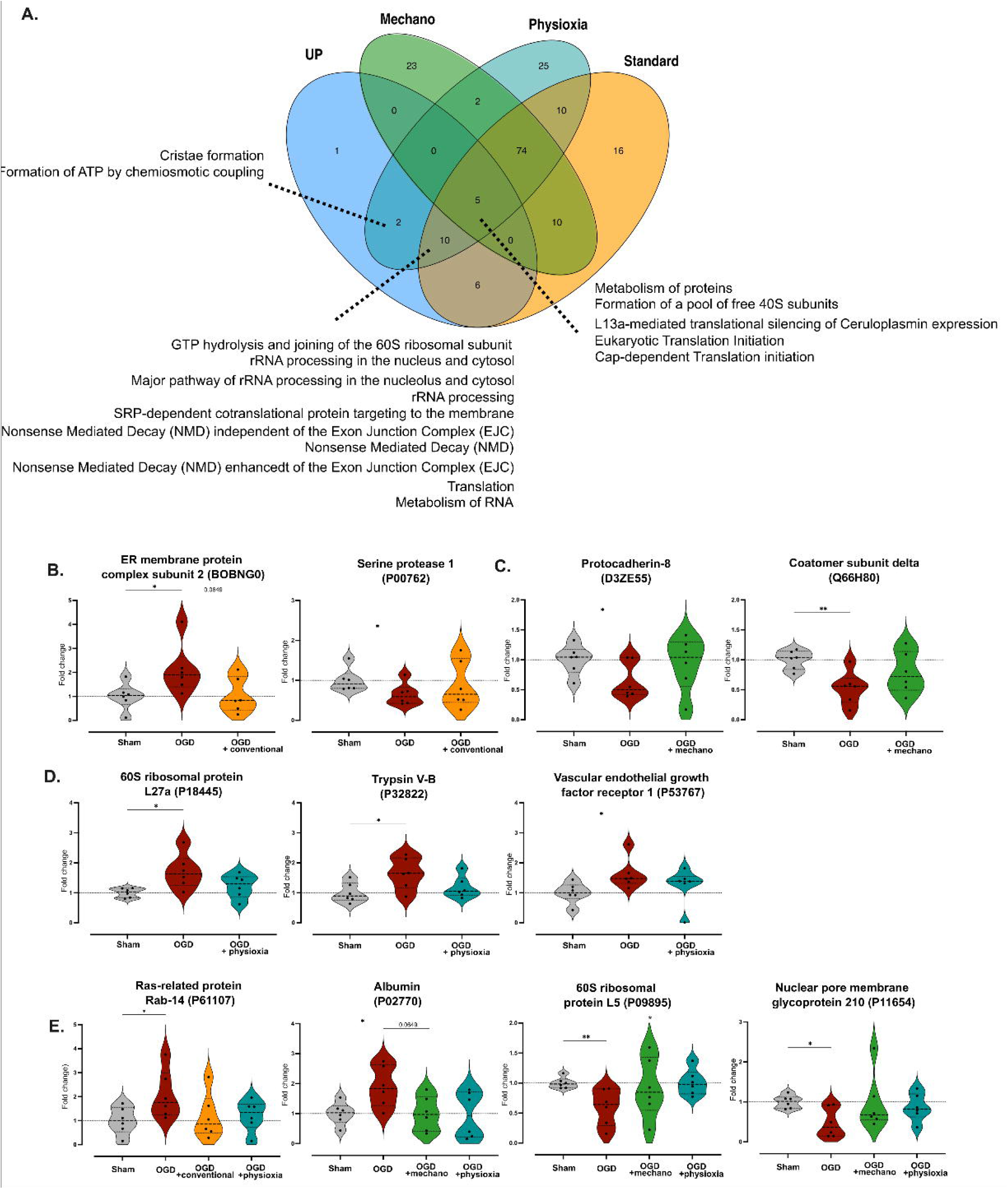
Neuroprotection by physioxia-modulated UC-MSCs secretome is correlated with alteration of neuronal translation. (A) Reactome pathways of upregulated proteins upon the OGD insult (Figure 2E) were compared to the rescue pathways identified by the clustering analysis (Cluster SC2, SM2, SP2). DEPs previously identified by comparing sham and OGD conditions (Figure 1) but restored to sham levels by the presence of conventional (B), mechanomodulated (C), or physioxia-modulated secretome (D) or more than one secretome (E) are represented (Mann-Whitney; *p<0.05; **p<0.01).

Finally, to identify proteins differentially expressed by the presence of UC-MSCs secretome, a non-parametric univariate approach (Mann-Whitney) was performed. Proteins that exhibited statistically significant expression changes when comparing neurons under sham conditions to those exposed to OGD were selected. The analysis focused on proteins whose alterations were reversed in the presence of the secretome in each experimental setup. Results were then compared to the two selected clusters (Supplementary Figure 7F-H, Supplementary Table 5-7). Proteins selected by both approaches are listed in Table 2. Comparing the proteins altered by the presence of the three secretome treatments, physioxia appears to modulate the highest number of proteins (twenty-four), sharing eight proteins with the results obtained when the mechano-modulated secretome was added to cultured neurons (Supplementary Figure 7I). Three, six, and eight proteins were previously identified as DEPs comparing sham to OGD but their levels were restored in the presence of conventional, mechano-modulated, or physioxia-modulated UC-MSCs secretome, respectively (Figure 6B-E). In conclusion, not only does the physioxia-modulated secretome appear to modulate a larger number of pathways on OGD-treated neurons, but it also shows a larger number of treatment response indicators, suggesting that these candidates are responsible for sustaining the neuronal network.

**Table 2.**
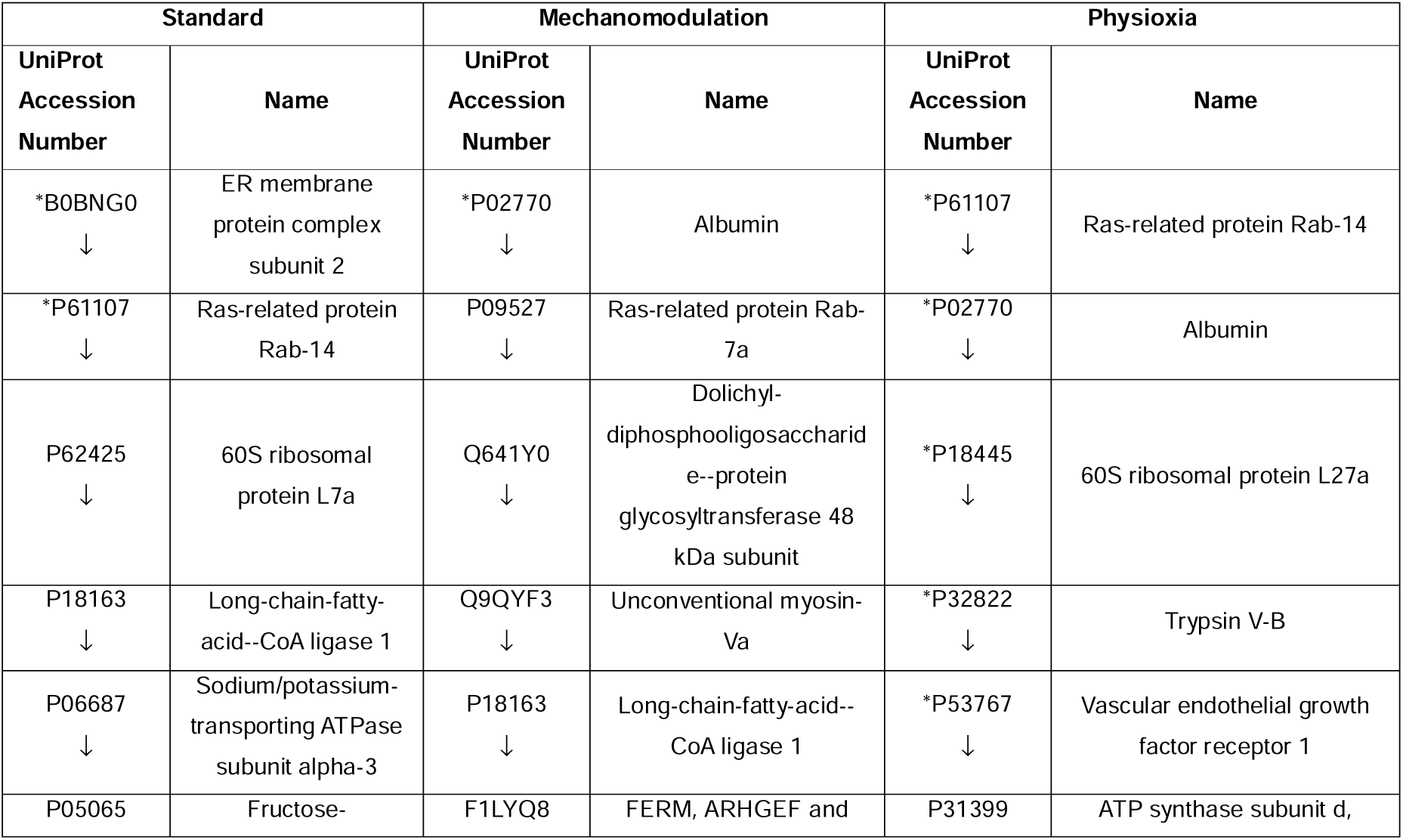
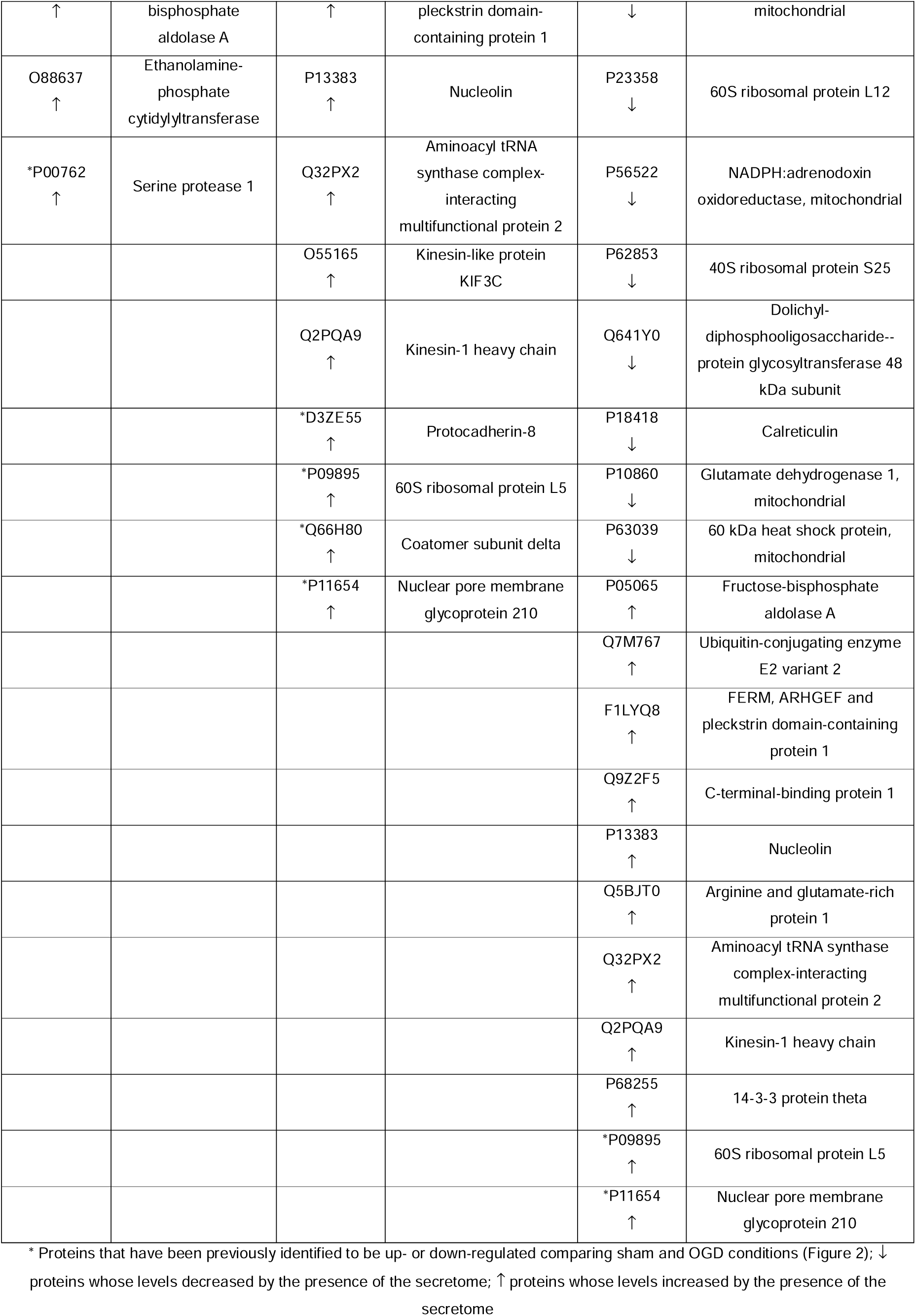
Differentially expressed proteins in the neuronal proteome after an OGD insult by the presence of the conventional, mechano-modulated, or physioxia-modulated UC-MSCs secretomes.

### Addition of physioxia-modulated secretome allowed the identification of potential treatment response indicators

Since the physioxia-modulated secretome was the only treatment that demonstrated a capacity to sustain the neuronal network integrity of the neurons, candidate proteins altered by the presence of this secretome were tested using unsupervised statistical methods. For that, the treatment response indicator candidates previously identified - 60S ribosomal protein L27a (P18445), Ras-related protein Rab-14 (P61107), Albumin (P02770), 60S ribosomal protein L5 (P09895), Nuclear pore membrane glycoprotein 210 (P11654), Trypsin V-B (P32822) and Vascular endothelial growth factor receptor 1 (P53767) (despite the last three were not quantified on the neurons treated with UC-MSCs) - were analyzed using heatmaps with hierarchical clustering (Figure 7A and B). For both screenings, it was possible to separate both sham and post-OGD neurons, with a closer relationship between sham and physioxia modulated secretome treated neurons. Accordingly, the PCA does not show an overlap between sham and post-OGD neurons, and the treatment group with physioxia-modulated secretome is the only one that shows a tendency to overlap with sham neurons (Figure 7C, Supplementary Figure 8A and B). However, it should be noted that PCA models for the UC-MSCs treated neurons show a higher dispersion of the data, possibly because of the lower sample size. Nevertheless, the separation of the experimental groups, and its applicability for the two independent screenings, indicate that these proteins are good candidates for a panel of treatment response indicators. Two of these proteins are ribosomal subunits, proteins known to be involved in translation processes. This suggests that the therapeutical targets of physioxia secretome might interfere with the metabolism of proteins that possibly play a key role in sustaining neuronal survival.

**Figure 7.**
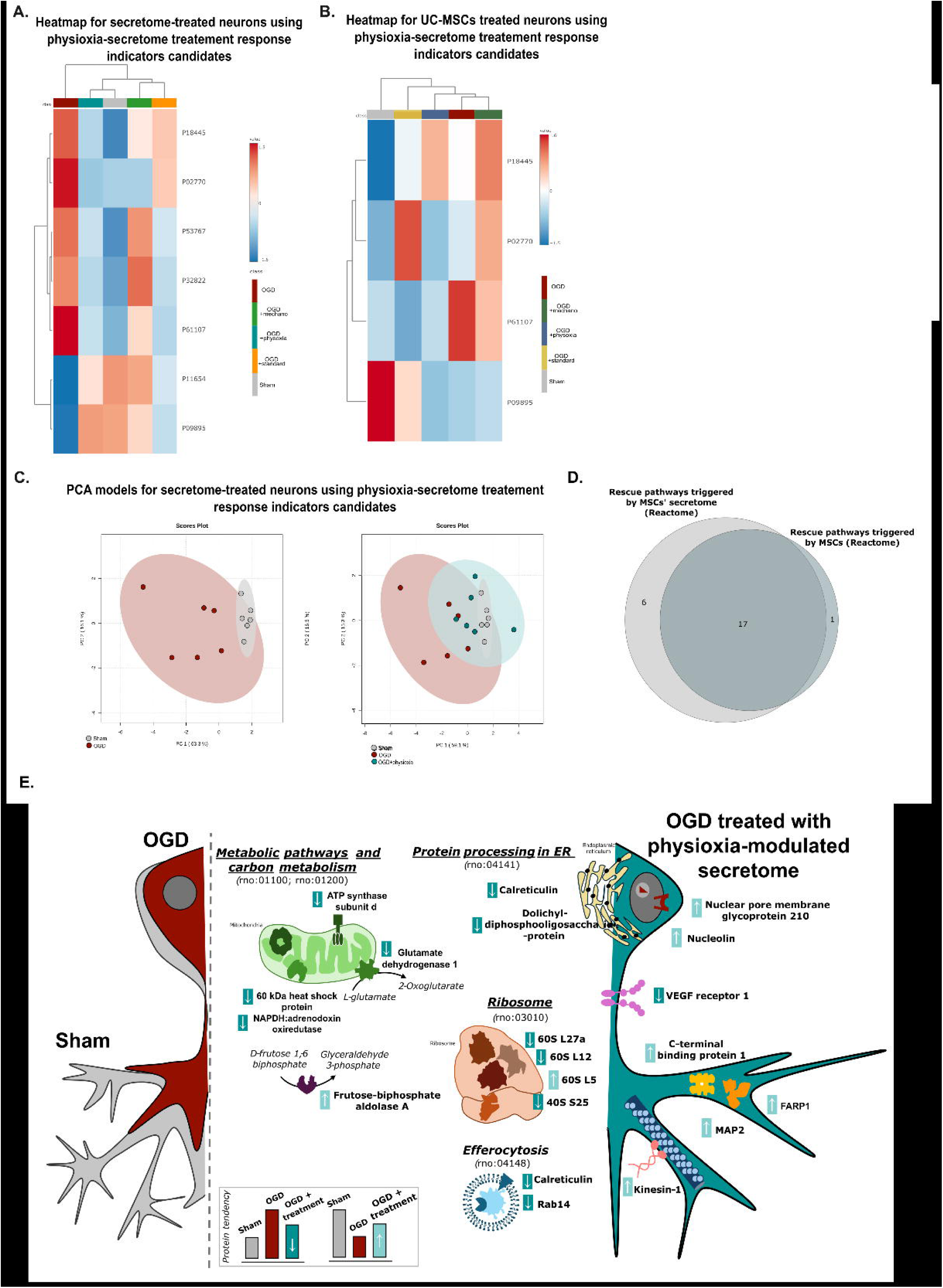
Panel of candidates for treatment response indicators. Proteins previously found to be differentially expressed in neurons treated with physioxia-modulated secretome (Figure 6D and E) were analyzed using heatmaps with hierarchical clustering for the secretome (A) and UC-MSCs (B) treatment. PCA models of sham, OGD and physioxia-modulated secretome treated post-OGD neurons (C). Common Reactome pathways altered by the presence of standard or physiologically modulated UC-MSCs or their secretome. (D) Shematic overview of the treatment response candidates and associated Kegg Pathways.

Moreover, the proteins and pathways altered by the two types of treatments (secretome or co-culture) were compared. Although no single molecules are shared by both approaches, the results indicate that, regardless of the therapeutic approach, common rescue mechanisms rescue are triggered, namely translation initiation and L13a-mediated translational silencing of Ceruloplasmin expression processes (Figure 7D; Table 3). Therefore, it can be concluded that maintaining proteostasis is crucial to sustaining the neuronal network integrity. However, further extensive experiments are required to validate the specific mechanism.

**Table 3.**
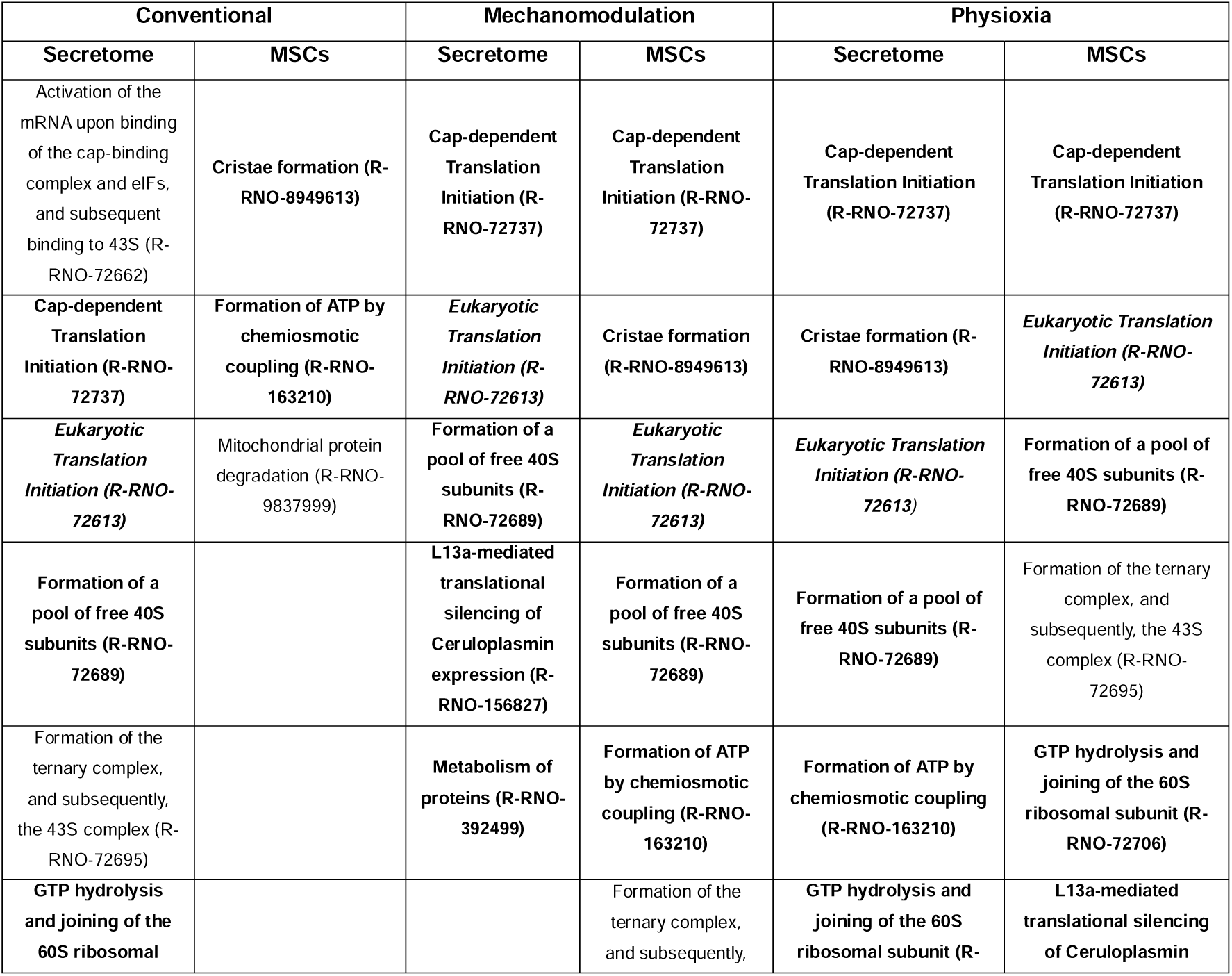
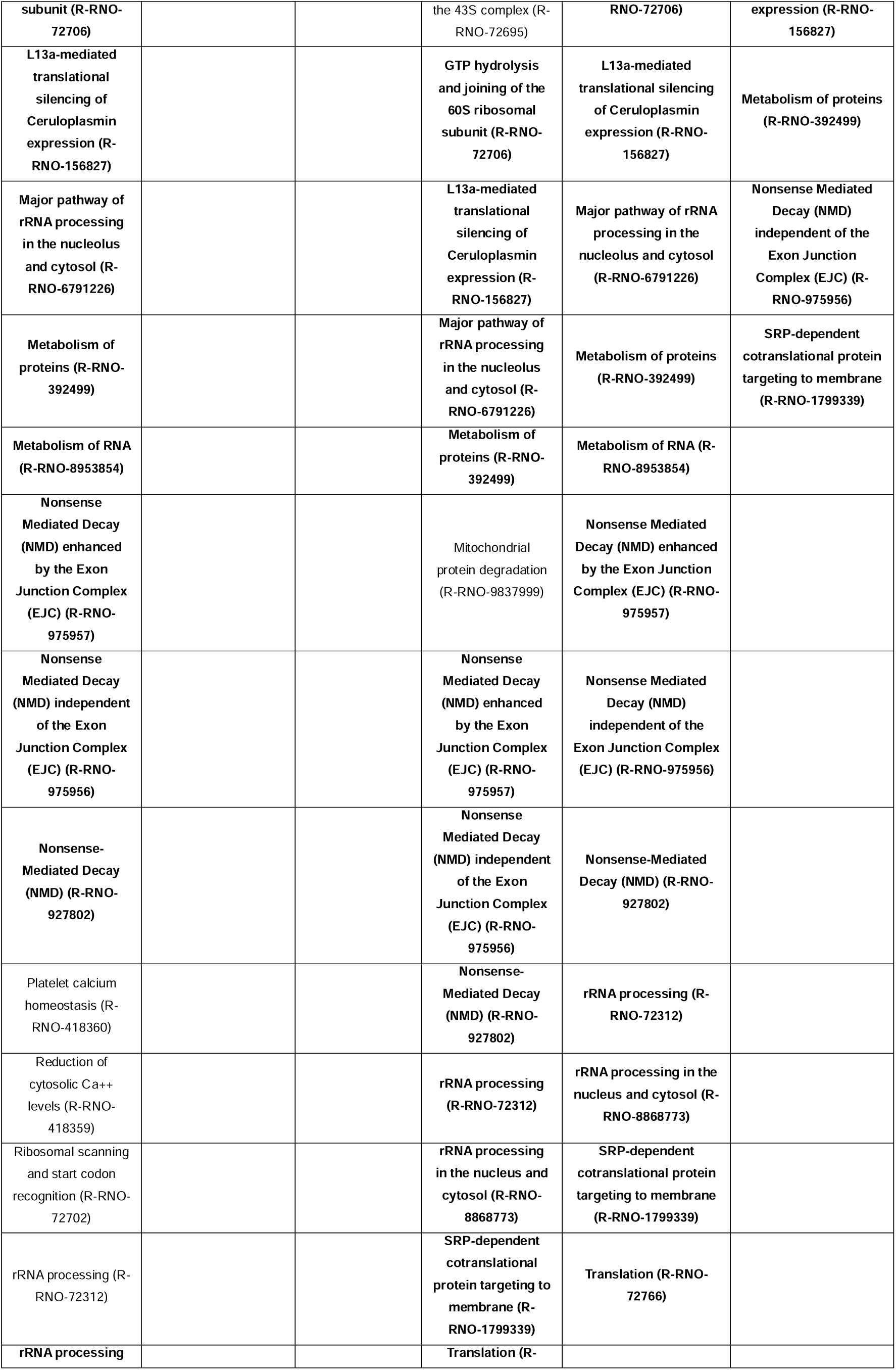
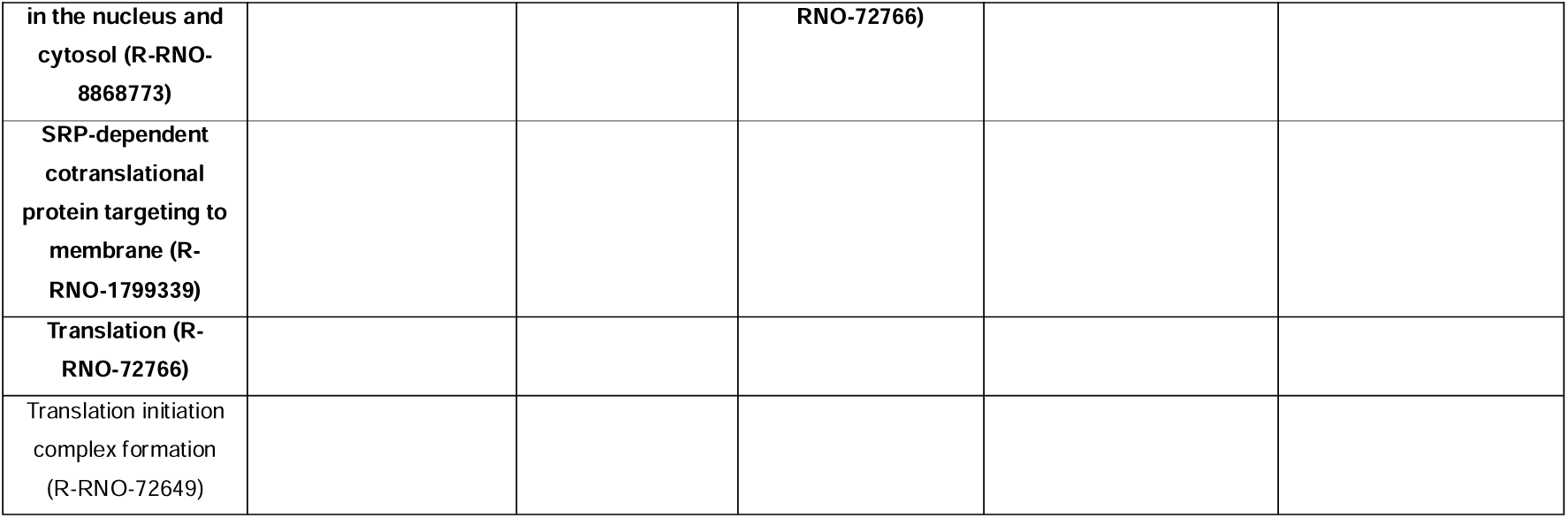
Reactome pathways associated with proteins whose levels were restored in the presence of conventional or physiologically modulated UC-MSCs or their secretome.

## Discussion

HIE is considered a rare disease but one of the leading causes of child death and disability (5, 11). Currently, the lack of effectiveness of hypothermia has prompted the development of innovative therapeutic strategies, such as MSCs. Our team has previously reported that priming MSCs at their physiological stiffness (3kPa) or oxygen levels (5% O_2_) induces changes in the protein signatures, both at the cellular and secretome level (28). Therefore, an in vitro HIE model was used to evaluate the therapeutic potential of these cell-based treatments.

A newborn brain is immature compared to an adult brain. Thus, in order to mimic a less extensive neuronal network, the primary cortical neurons were maintained in culture for 7 days prior to OGD insult instead of the 14 days usually used in the case of the stroke model (30, 47). Since HIE pathophysiology relies on excitotoxicity pathways, a glutamate assay was performed to confirm the functional expression of glutamate receptors (Figure 1A and B). Additionally, the responsiveness of this cellular model to an OGD insult at this time point was assessed. The loss of the neuronal cytoskeleton after this stimulus was confirmed by imaging and proteomics, indicating cell damage as neurons rely on their network for proper communication and subsequent survival (Figure 1C and G). Interestingly, proteins related to translation were found to be upregulated, as well as the levels of proteins associated with rRNA processing (Figure 1G and H). In the adult brain, a decrease in protein synthesis immediately after the insult has been observed, without irreversible damage to the ribosomes, which should prevent the accumulation of misfolded proteins (48–50). However, it is known that protein synthesis is reestablished hours later (51, 52). These pieces of evidence suggest that sixteen hours after the insult, higher levels of translation-related proteins may be involved in restoring proteostasis in surviving neurons in the experimental model used.

The addition of secretome with different compositions to neurons subjected to OGD led to significant modifications in their proteome profile (Supplementary Figure 5A). More specifically, the presence of physioxia-modulated secretome sustained part of the neuronal network integrity (Figure 4A). Also, the secretome of UC-MSC modulated under physioxia was the most effective in preventing the alterations in the abundance of proteins and unique pathways, as highlighted on the PLS-DA scores plot (Figure 4D and 5D). In particular, the presence of this secretome helped restore the basal levels of mitochondrial proteins, specifically ATP synthase subunits (Figure 6A, Supplementary Table 10). The regulation of oxidative stress levels, possibly through mitochondria transfer, may be crucial for restoring energy balance post-OGD and promoting the maintenance of the neuronal network (53, 54). Moreover, antioxidant proteins, such as AMBP (P02760) or Ceruloplasmin (P00450) were found to be more abundant on this type of secretome, which might be essential for balancing ROS levels (55, 56). In addition, physioxia-modulated secretome interfered with the levels of several other relevant proteins that could play a role in cell survival (Table 2, Figure 7E). For example, Vascular endothelial growth factor receptor 1 (VEGFR1) was shown to be increased in hippocampal neurons in a stroke model and described to activate a protective mechanism (57, 58). In fact, this protein has already been described as a possible biomarker for HIE diagnosis (59). Other neuroprotective proteins identified include Ras Family protein members, 14-3-3 or kinesin-1 (60–62). These findings underscore that the therapeutical properties of this fluid are not limited to the differential expression of a single protein but are due to a combination of factors that will synergistically work to promote neuronal survival, modulating several pathways simultaneously, as protein and carbon metabolism and neuronal cytoskeleton integrity. Future research should validate the activation of the canonical signaling pathways regulated by these proteins.

Conventional and physiologically modulated UC-MSCs were also co-cultured with neurons after the OGD insult and tested for their neuroprotective effects to evaluate the differences triggered in the neuronal culture. The different types of primed MSCs lead to the differential modulation of the neuronal proteome, suggesting that MSCs’ can somehow retain a memory of the culture conditions (Table 1), possibly leading to a differential secretion of factors. The concept of mechanical memory has previously been described, but to our knowledge, memory in response to oxygen levels has not been documented (63). Future studies should investigate how manipulating oxygen levels might induce long-term permanent changes in MSCs.

Although no common proteins were altered by secretome or co-culture treatment, the protein metabolism, namely translation, emerged as a common target (7D, Table 3). Following ischemia, protein synthesis in the brain is mainly controlled by translation initiation mechanisms. Initially, the GTP-linked eukaryotic initiation factor (eIF2_α_) recruits 40S small ribosomal subunits, the ternary complex, and other initiation factors. Upon recognition of the start codon, eIF2-GTP is hydrolyzed to eIF2-GDP, leading to the recruitment of 60S ribosomal subunits (48, 64). However, post-ischemia, the intracellular calcium overload causes endoplasmic reticulum stress, activating PERK (PKR-like endoplasmic reticulum eIF2_α_ kinase), which phosphorylates eIF2_α_ (49, 65, 66). These events result in the accumulation of untranslated transcripts and RNA-binding proteins, forming stress granules (64). Notably, several ribosomal subunits were found to be more abundant in physiologically modulated secretomes, possibly supporting alterations in the neuronal proteostasis after OGD (29). Restoring proteins involved in translation initiation mechanisms in neurons incubated with all types of secretomes and with all types of modulated MSCs demonstrate how these approaches can provide neuroprotection possibly by balancing proteostasis following the OGD insult (Table 3). However, it should also be noted that this treatment is susceptible to inter-individual variability, which might lead to differences in the treatment outcome (28, 67). Indeed, the variability observed in the data presented here is likely due to the use of different cell donors for each replicate.

In summary, physiologically modulated UC-MSCs, or their secretome, trigger different responses in post-OGD neurons compared to conventionally expanded cells. However, all treatments culminate in the modulation of translation initiation mechanisms, possibly interfering with protein metabolism. Physioxia-modulated secretomes can also modulate mitochondrial proteins, which might be essential for restoring the energy balance after the insult. The combination of reestablishing proteostasis and controlling oxidative stress might be key in sustaining the neuronal network, which is required for neuronal viability and to secure physiological functioning. Future studies should assess the activated neuronal pathways in response to this treatment to validate the potential therapeutic targets that promote cytoskeleton integrity and neuronal survival.

---

**Supplementary Figure 1.**
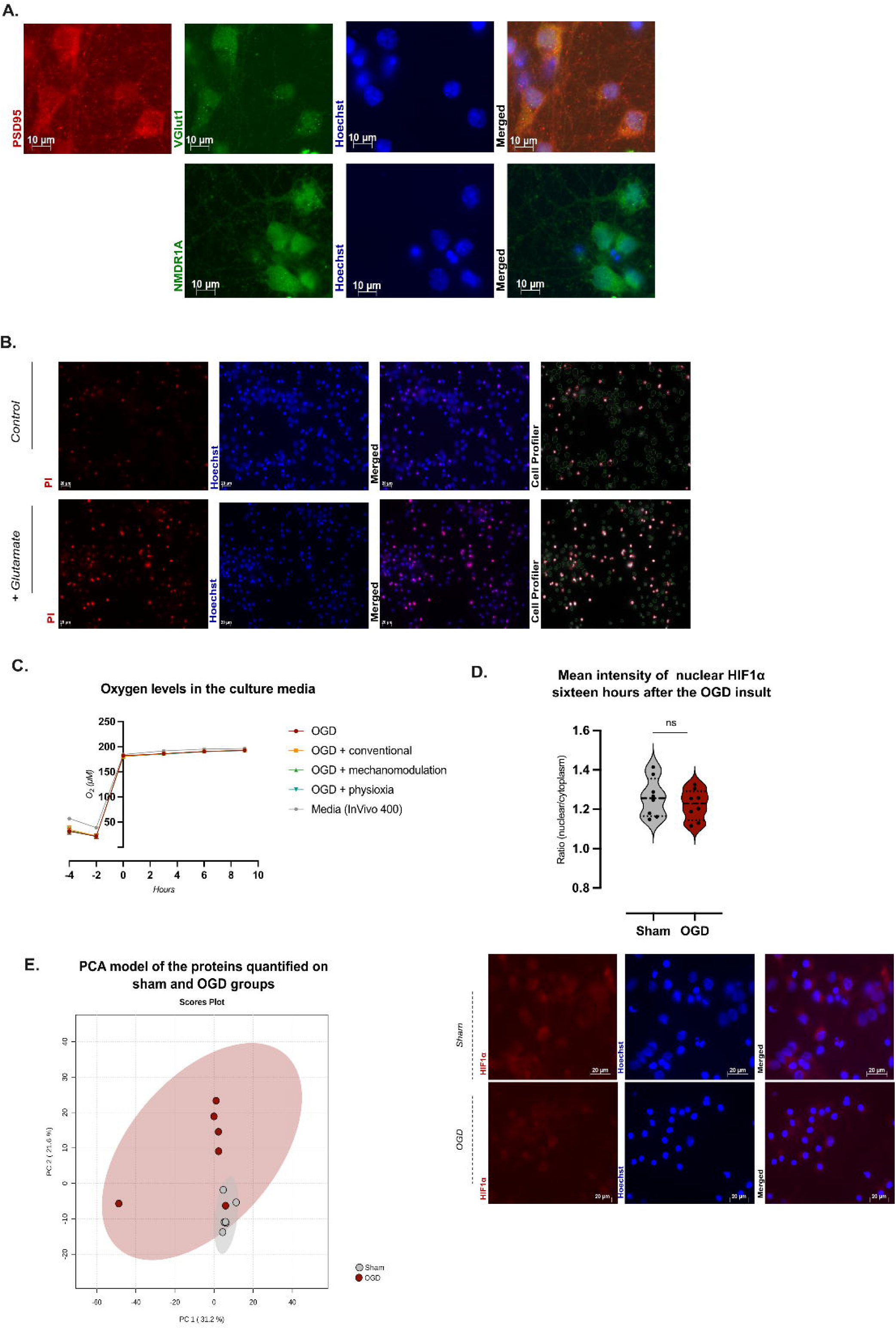
Primary cortical neurons express glutamatergic phenotype at DIV7. (A) Neurons were isolated from embryo cortices and left to mature for seven days, where the expression of PSD95, VGlut1 and NMDAR1 was confirmed. (B) Neurons were subjected to an excitotoxicity glutamate stimulation, and after six hours, the number of PI-positive cells was calculated (representative images). In addition, neurons were subjected to a five hours OGD insult. (C) The oxygen levels present in the culture media were measured using Resipher, where t=0h is the beginning of the reperfusion period. (D) The mean intensity of Hif1α was quantified on sham and OGD groups sixteen hours after the OGD insult (Mann-Whitney test; ns>0.05). (F) PCA model of all quantified proteins (using LC-MS/MS) on sham and OGD groups. Abbreviations: OGD (oxygen-glucose deprivation).

**Supplementary Figure 2.**
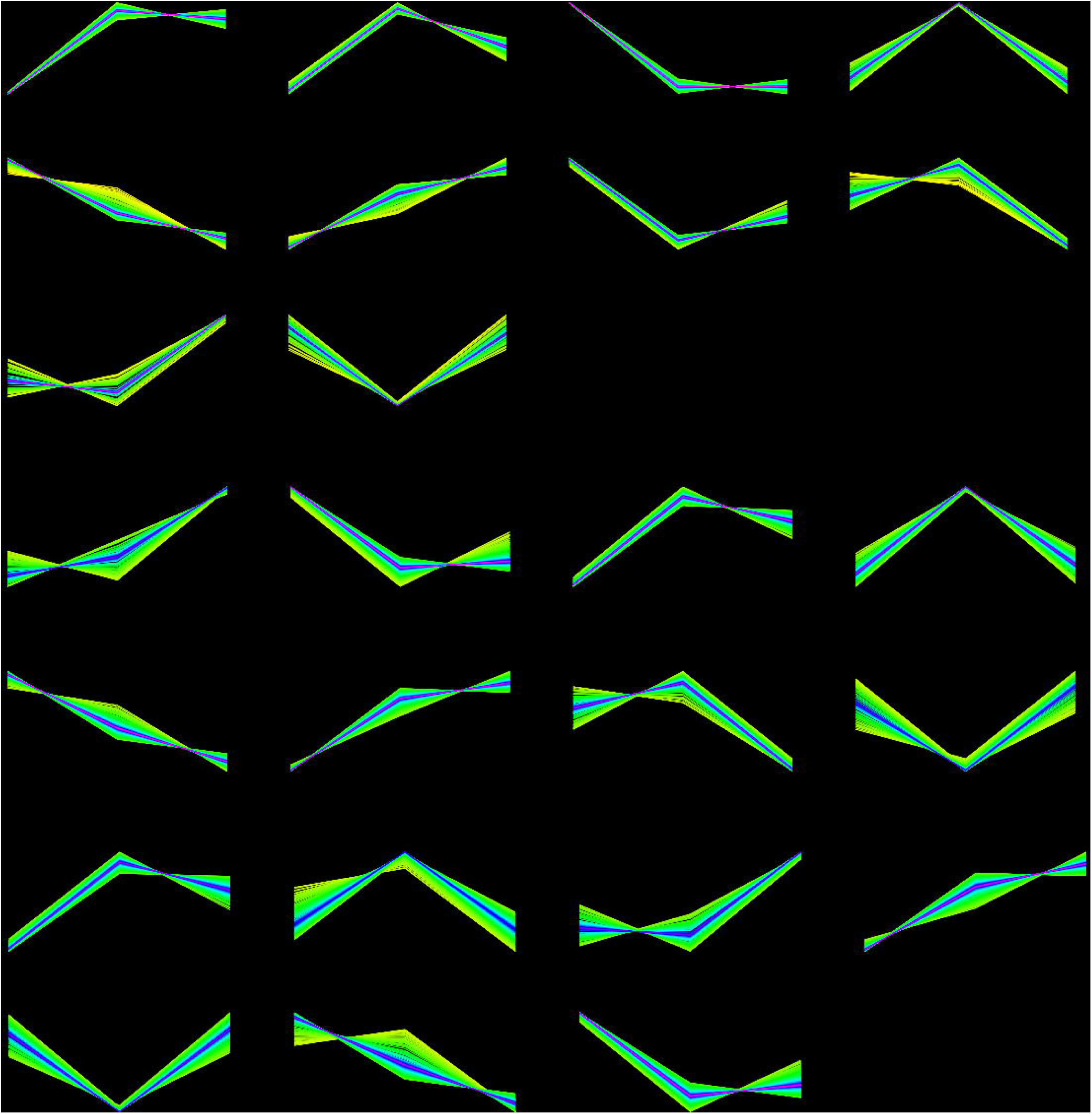
Clustering analysis of proteomics data from neurons co-cultured with UC-MSCs after the OGD insult. Proteins were clustered to identify pattern trends by comparing sham, OGD, and OGD-treated neurons with conventional (A), mechanomodulated (B), and physioxia-modulated (C) UC-MSCs. Abbreviations: CC (Cells conventional); CM (Cells mechanomodulated); CP (Cells physioxia-modulated)

**Supplementary Figure 3.**
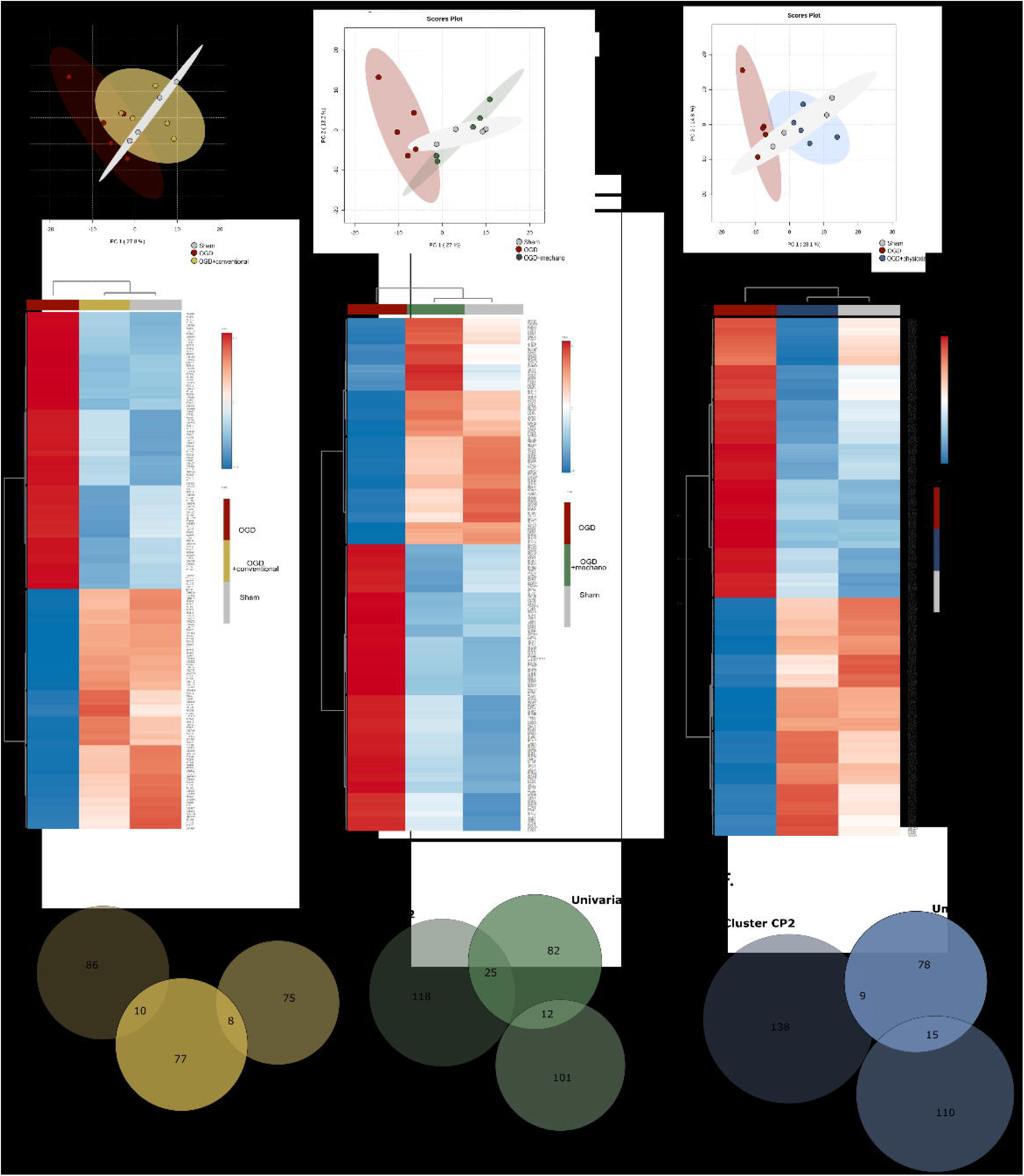
Co-culture of physiologically modulated UC-MCSs differentially regulates the neuronal proteome. Standard and physiologically modulated UC-MSCs were added to the neuronal culture after the OGD insult and left in culture for an additional sixteen hours. Proteins grouped on the clustering analysis for conventional (A), mechano-modulation (B) and physioxia-modulated (C) UC-MSCs are represented in the PCA (top) and heatmaps (bottom). For each experimental condition, clustering and univariate approaches were compared (D-F). Abbreviations: CC (Cells conventional); CM (Cells mechanomodulated); CP (Cells physioxia-modulated)

**Supplementary Figure 4.**
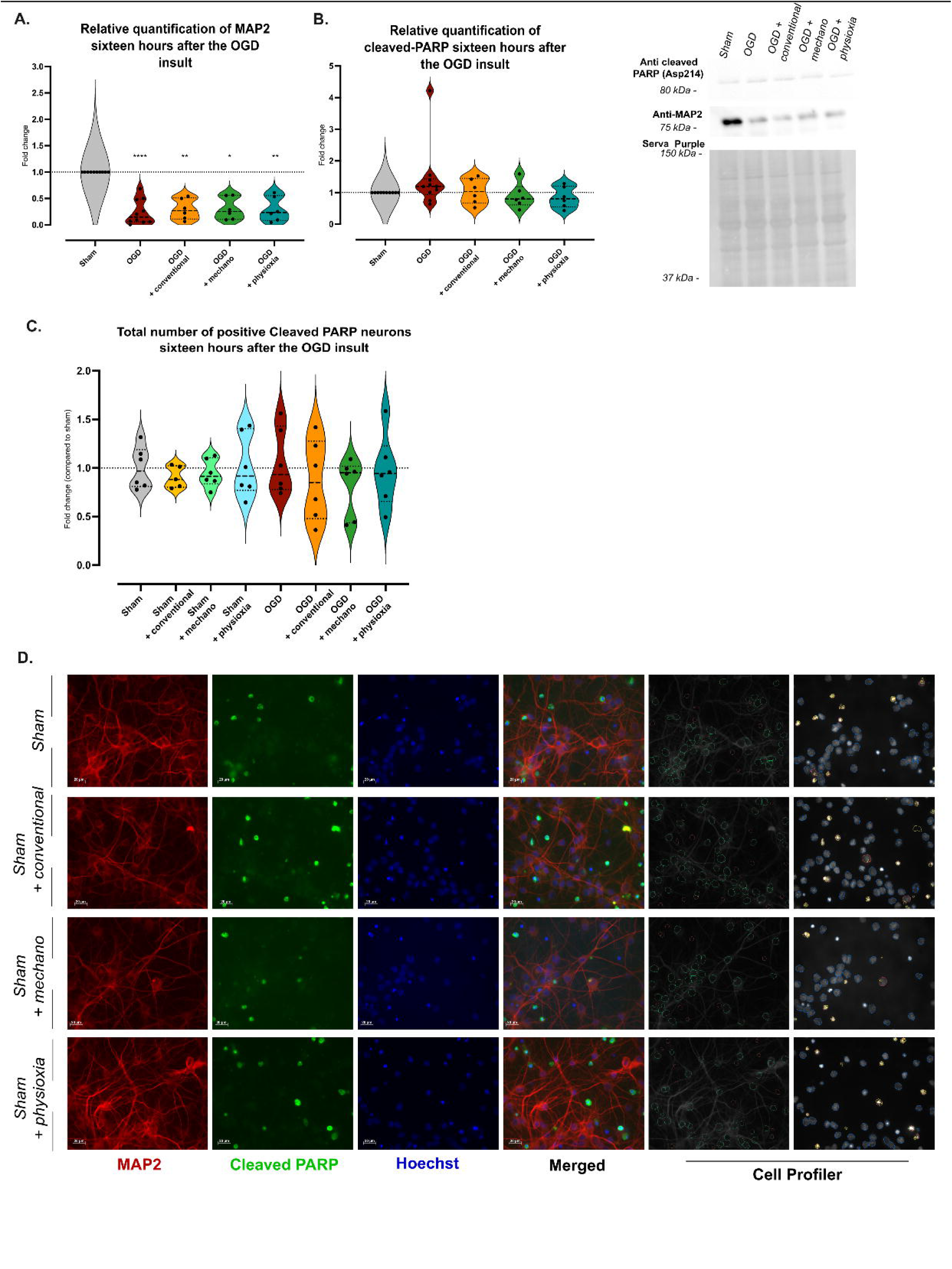
Evaluation of the neuronal cytoskeleton integrity in secretome treated neurons after an OGD insult. Neurons were subjected to a five hours OGD insult and then incubated with CM obtained from UC-MSCs cultured under conventional culture conditions or primed to low stiffness environments (3kPa) – mechano-modulation – or physioxia-modulated (5%O_2_). The total levels of MAP2 and cleaved PARP were quantified by Western blot (A and B, representative western-blot on the right). The total number of positive cleaved-PARP cells was determined by ICC (C). Representative images of the sham conditions incubated with physiological primed-MSCs’ secretome are displayed on (D). On the right, CellProfiler classification settings are highlighted: positive MAP2 neurons (green); negative MAP2 neurons (orange); cleaved PARP (yellow); positive cleaved PARP cells (red): negative cleaved PARP cells (blue). PCA model of all quantified proteins on sham and OGD groups (F). Abbreviations: Mechano (mechano-modulation); OGD (oxygen-glucose deprivation).

**Supplementary Figure 5.**
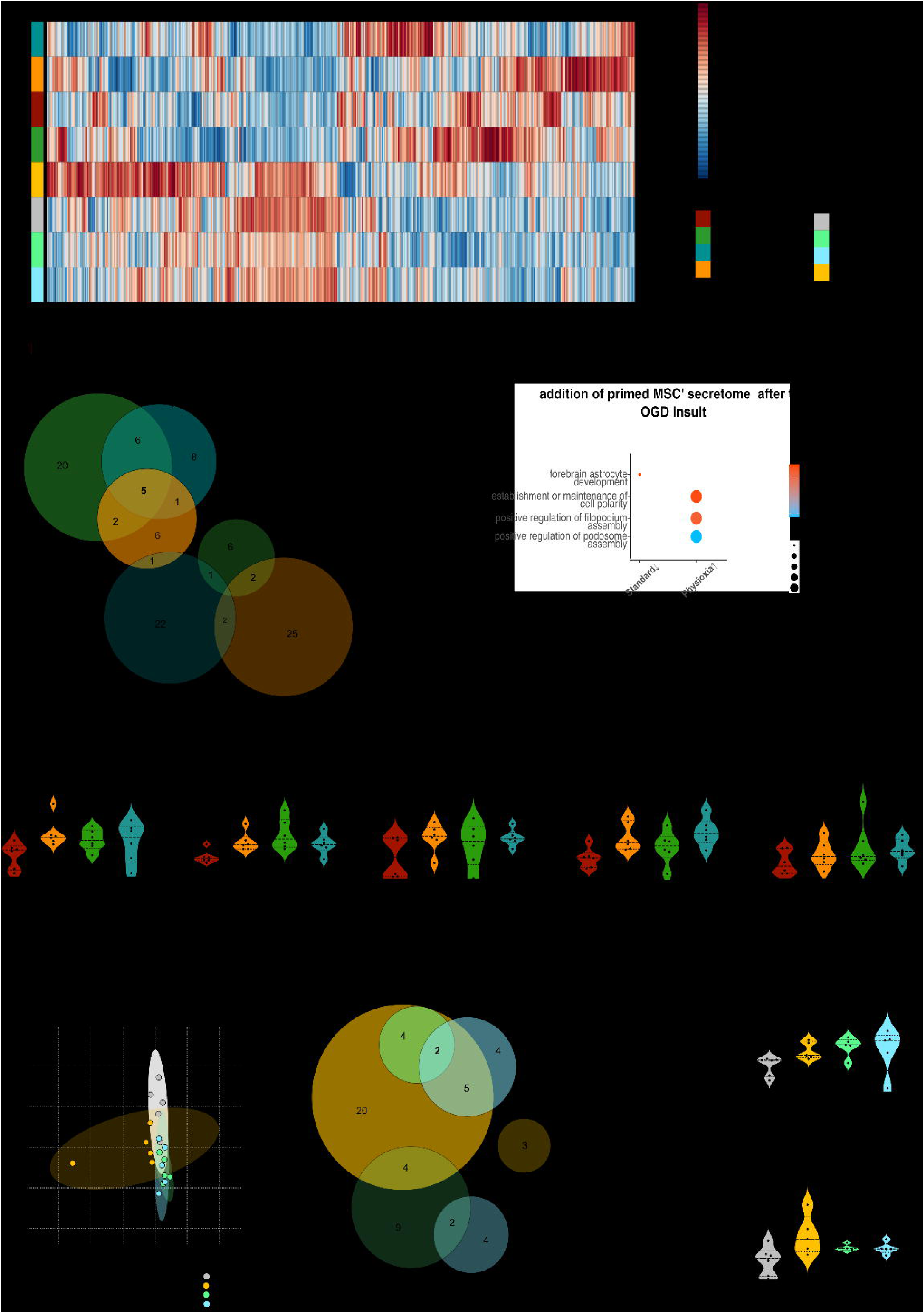
Physiological modulation of UC-MSCs’ secretome triggers different mechanisms on cortical neurons. After the OGD insult, either standard culture conditions or physiologically modulated UC-MSCs CM (either at 3kPa substrates (mechano) or 5%O_2_ (physioxia) was added to both sham and OGD neurons. Proteins quantified on all experimental conditions are represented on the heatmap (A). For OGD-treated and non-treated neurons, DEPs for each treatment were compared (VIP>1, p<0.05, or both; |Log_1.5_FC|>1) (Supplementary Table 6) (B), and a GO analysis (Biological processes) of these proteins are represented (C). Five proteins were found to be commonly upregulated by all three secretomes: AP-2 complex subunit mu, Rho-related GTP-binding protein RhoB, PAT complex subunit CCDC47, Tubulin alpha-1A chain, and Nuclear pore membrane glycoprotein 210 (D) (Mann Whitney; *p<0.05; **<0.01). The same analysis considered only sham and sham-treated neurons (Supplementary Table 7) (E, F). Two proteins were found to be commonly upregulated by the three secretomes: 60S ribosomal protein L27 and Eukaryotic translation initiation factor 3 subunit C (G) (Mann Whitney; *p<0.05; **<0.01). Abbreviations: Mechano (mechano-modulation); OGD (oxygen-glucose deprivation).

**Supplementary Figure 6.**
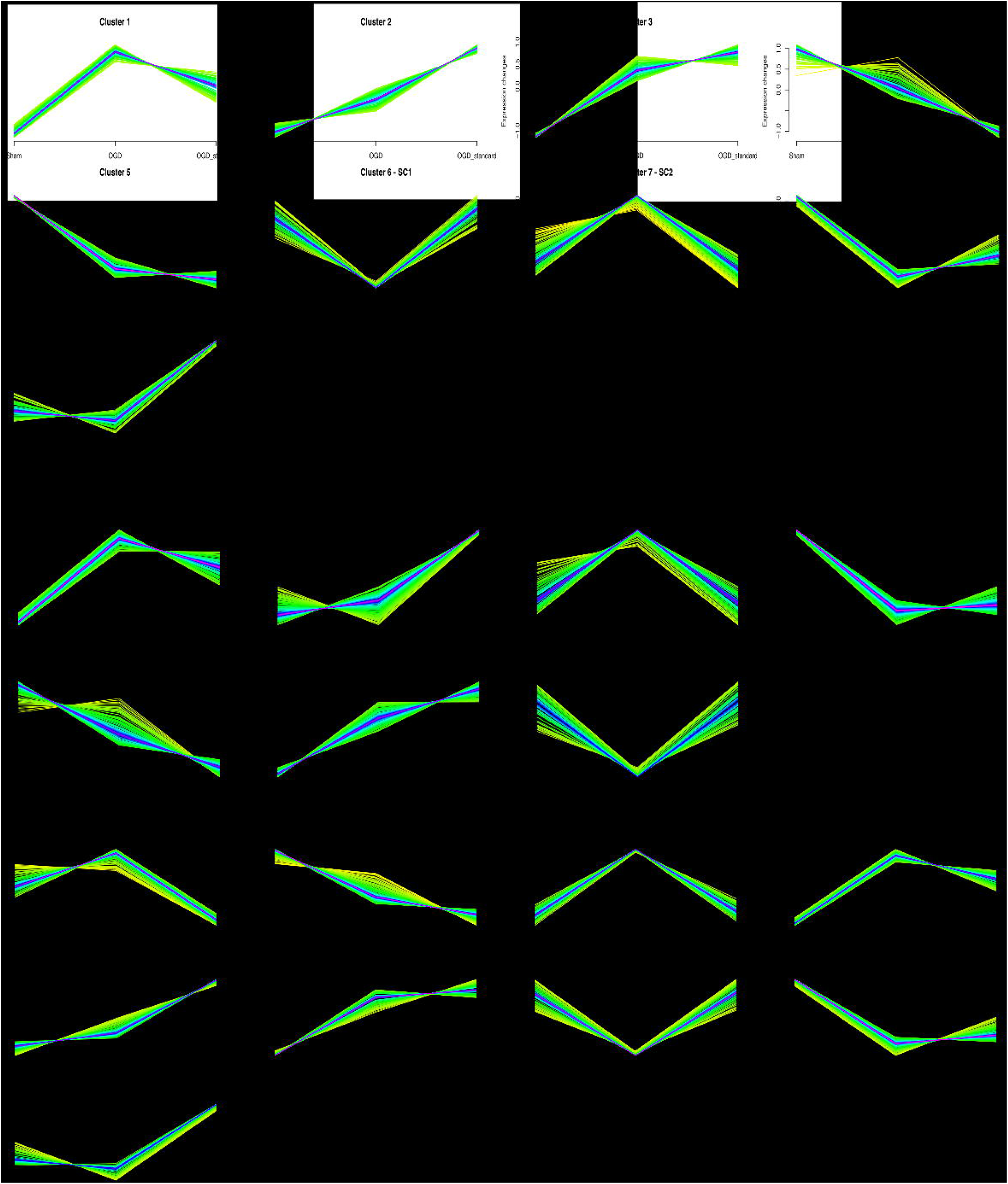
Clustering analysis of the neurons treated with UC-MSCs secretome after the OGD insult. The neuronal proteome was analyzed, and proteins were clustered to identify pattern trends by comparing sham, OGD, and OGD-treated neurons with conventional (A), mechano-modulated (B), and physioxia-modulated (C) UC-MSCs’ secretome. Abbreviations: SC (Secretome conventional); SM (Secretome mechanomodulated); SP (Secretome physioxia-modulated)

**Supplementary Figure 7.**
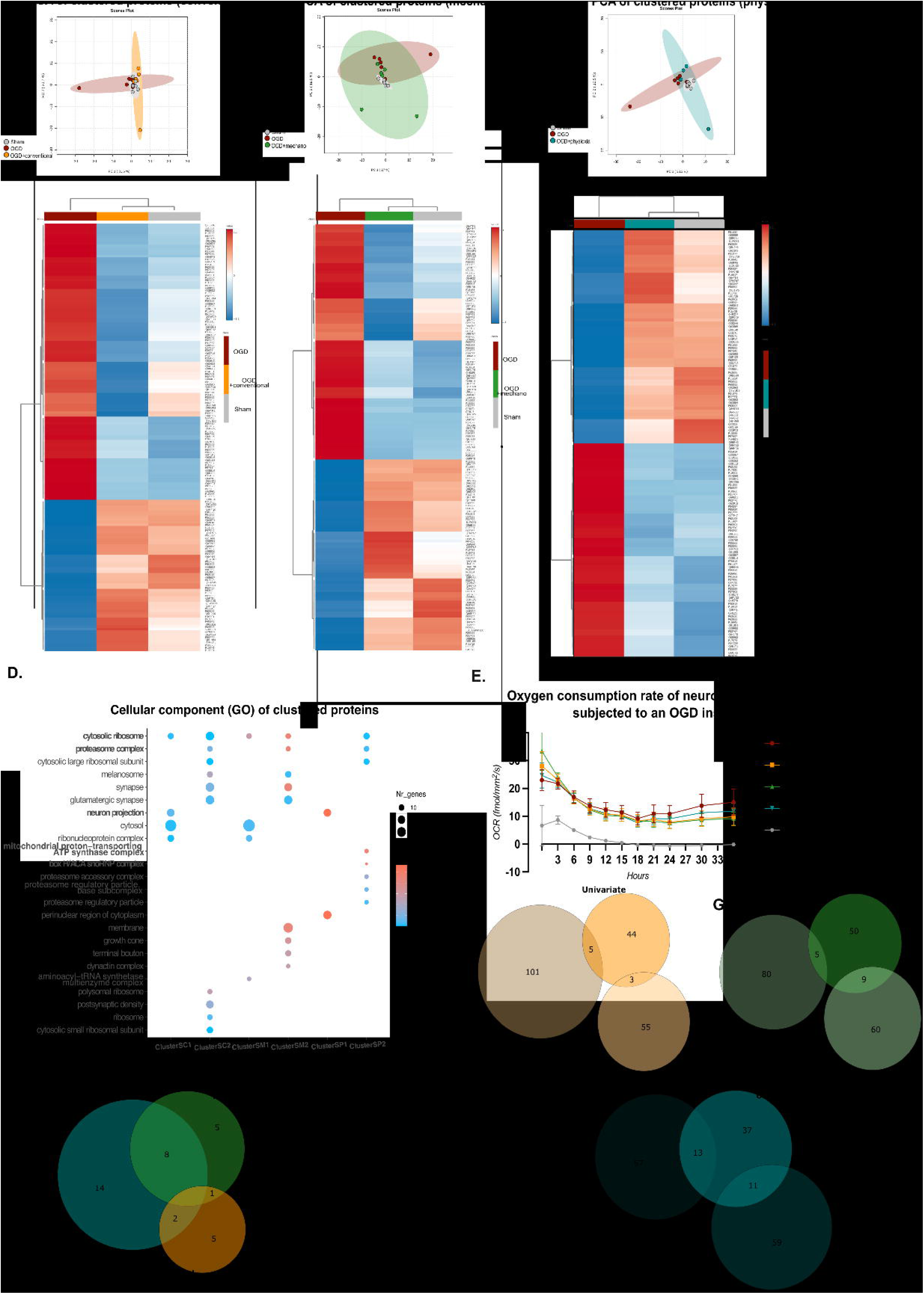
Modulation of MSCs’ secretome composition triggers differential responses on neurons, after the OGD insult. PCA and heatmaps represent proteins previously clustered (Figure 5A) for neurons treated with conventional (A), mechano-modulated (B) and physioxia-modulated (C) secretome. Cellular components (GO) of proteins present on each cluster are highlighted (D) The OCR during the recovery period, after the OGD insult was measured for 36h (E). t=0h is the beginning of the recovery period. Proteins grouped by the clustering and selected through the univariate approach (Mann-Whitney, p<0.05), but that lost their significance after the addition of each secretome) were compared (Supplementary Table 8-10) (F-H). Common altered proteins selected for each experimental condition were compared (I) (Table 1). Abbreviations: SC (Secretome conventional); SM (Secretome mechanomodulated); SP (Secretome physioxia-modulated)

**Supplementary Figure 8.** PCA models using treatment response indicators candidates. Proteins previously selected as promising treatment response indicators were analyzed by PCA. Models show neurons incubated with their UC-MSCs secretome (A) or co-cultured with UC-MSCs (B)

**Supplementary Table 1.**
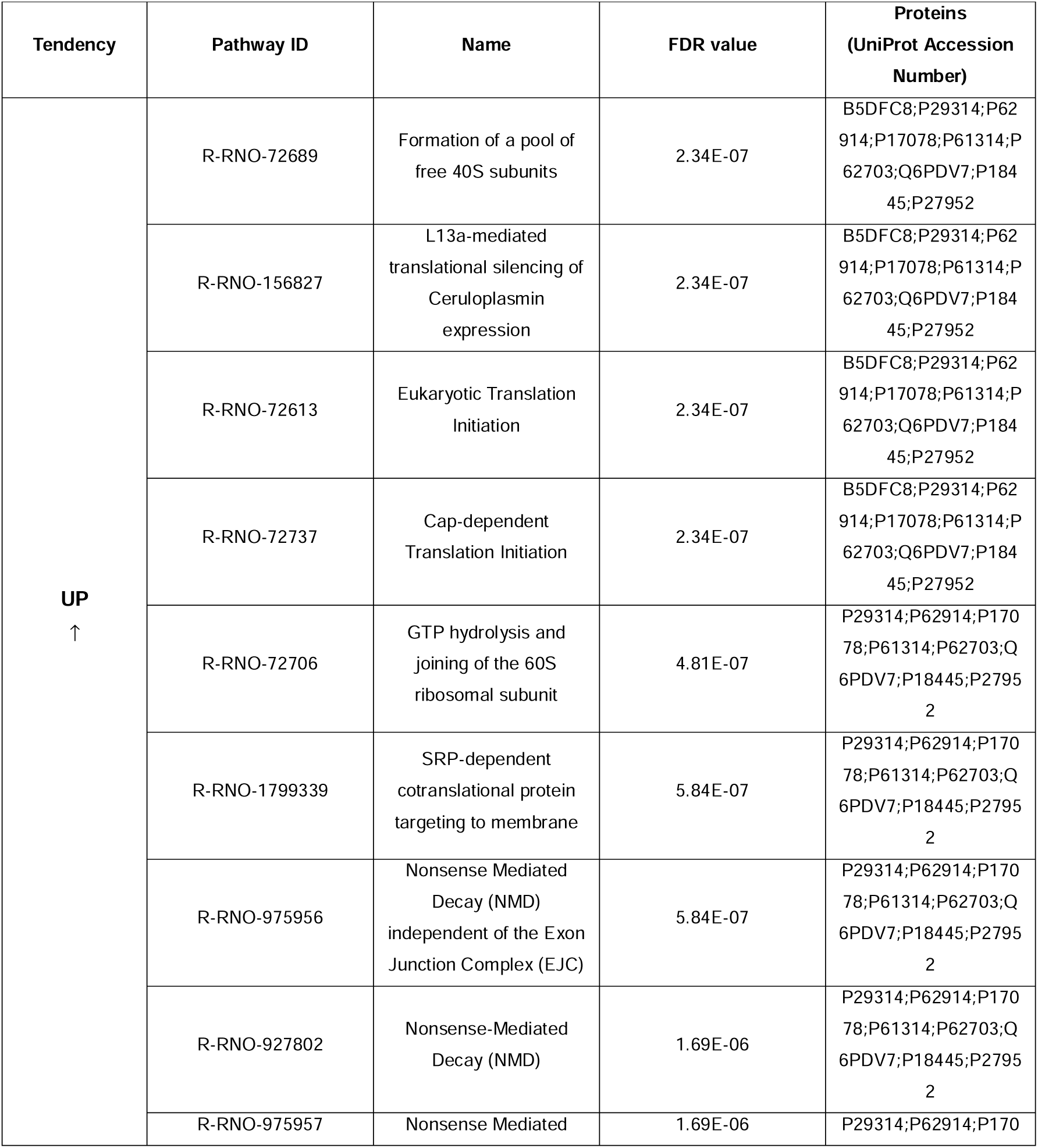
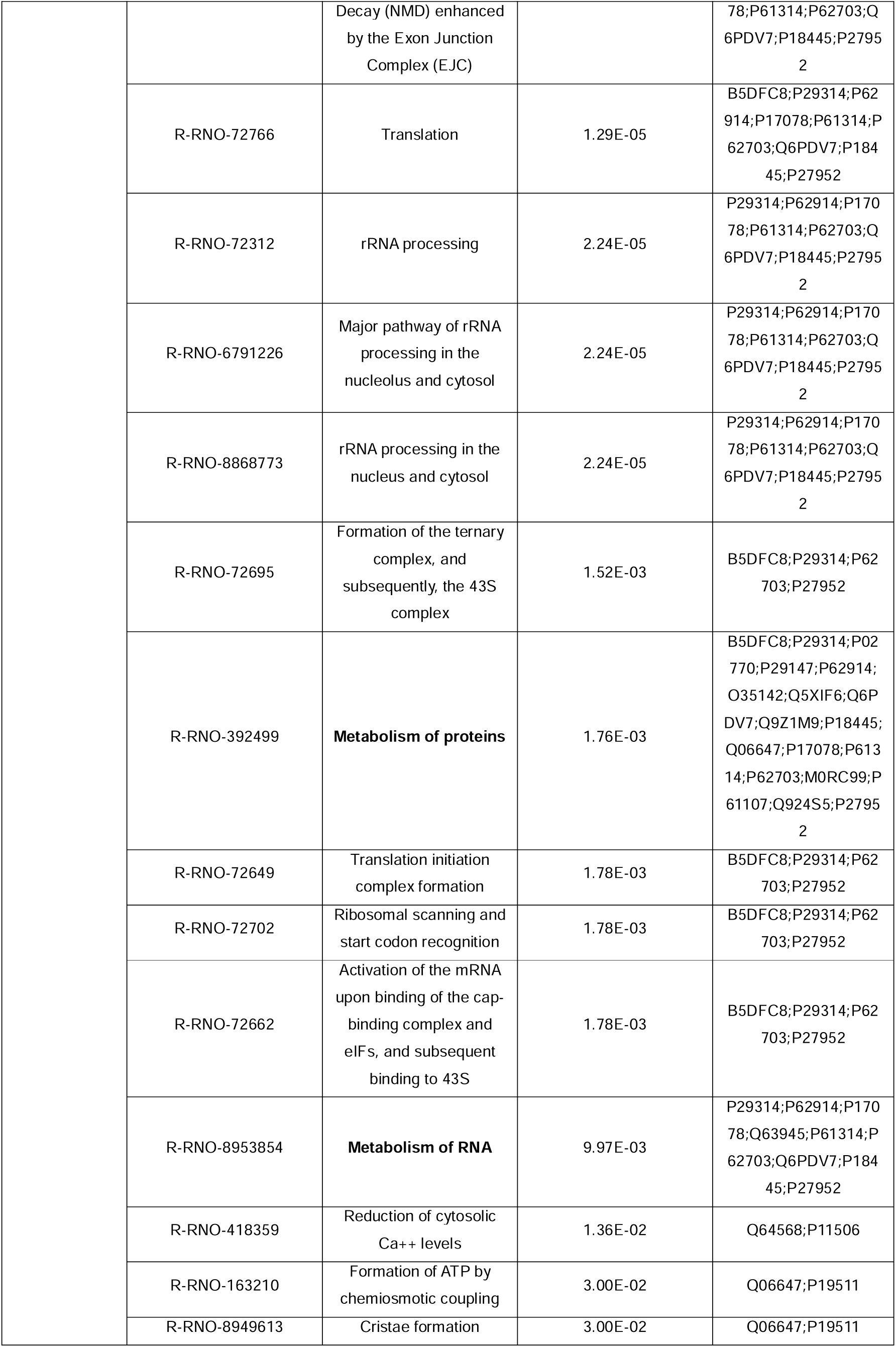
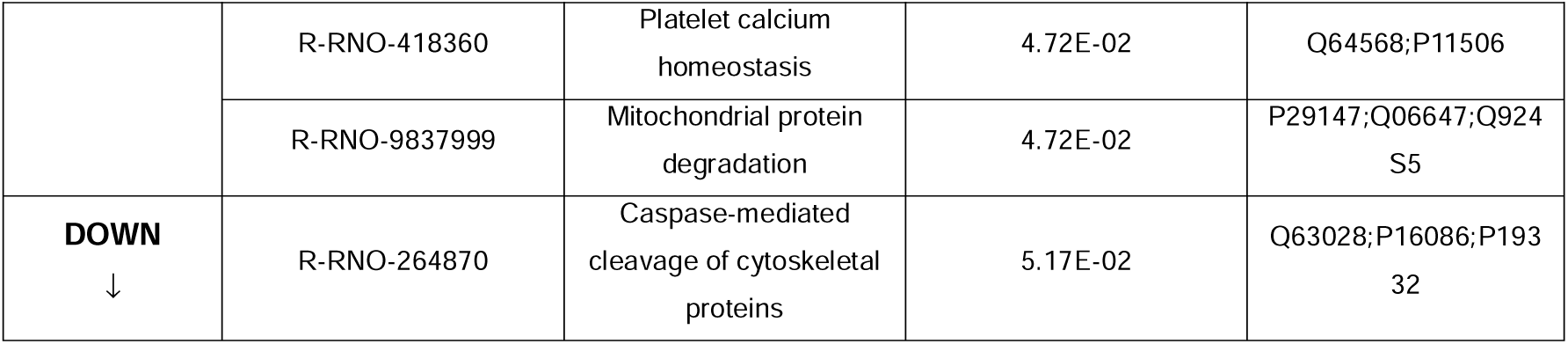
Reactome pathways of proteins significantly altered between sham and OGD conditions (Figure 1H).

**Supplementary Table 2.**
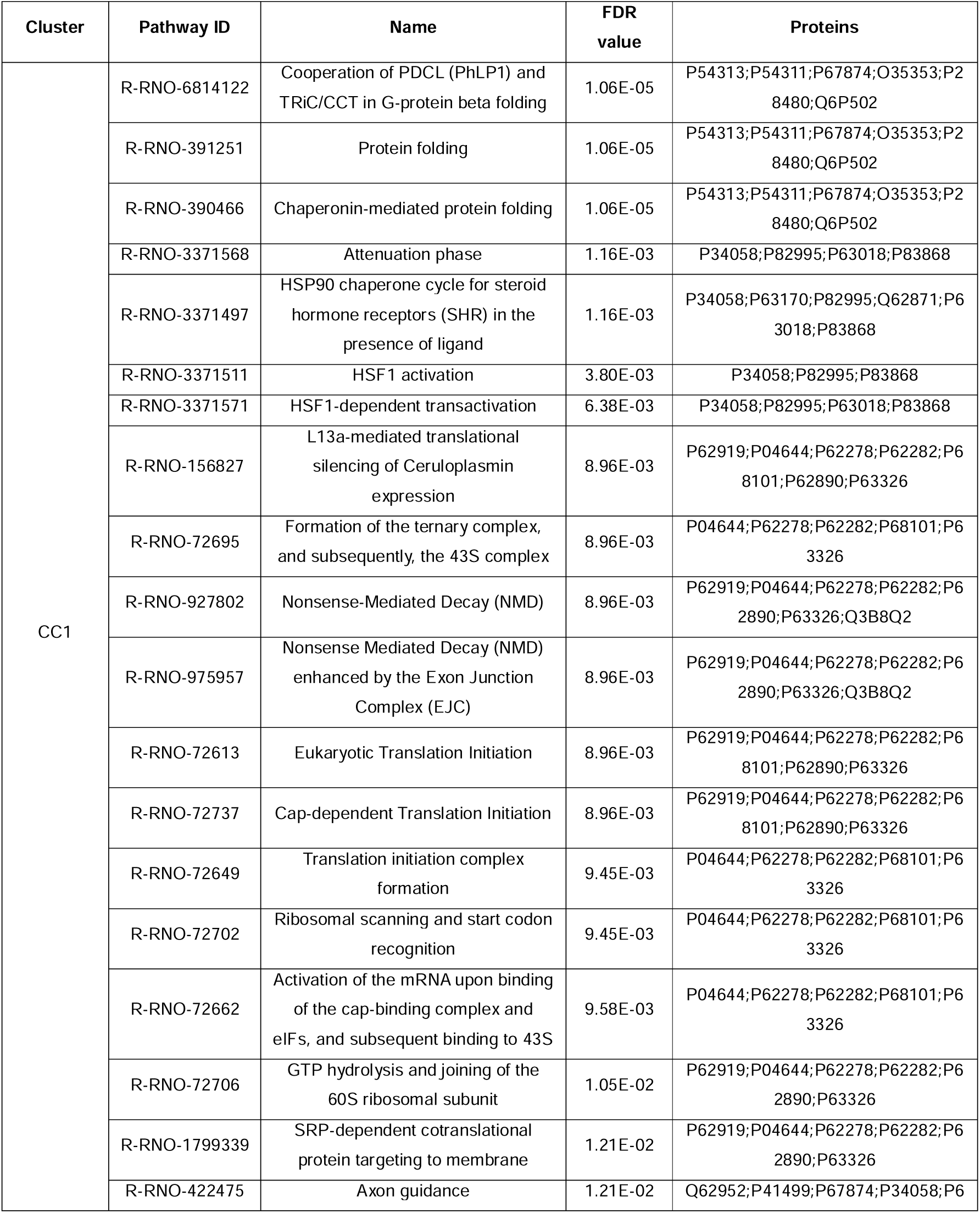
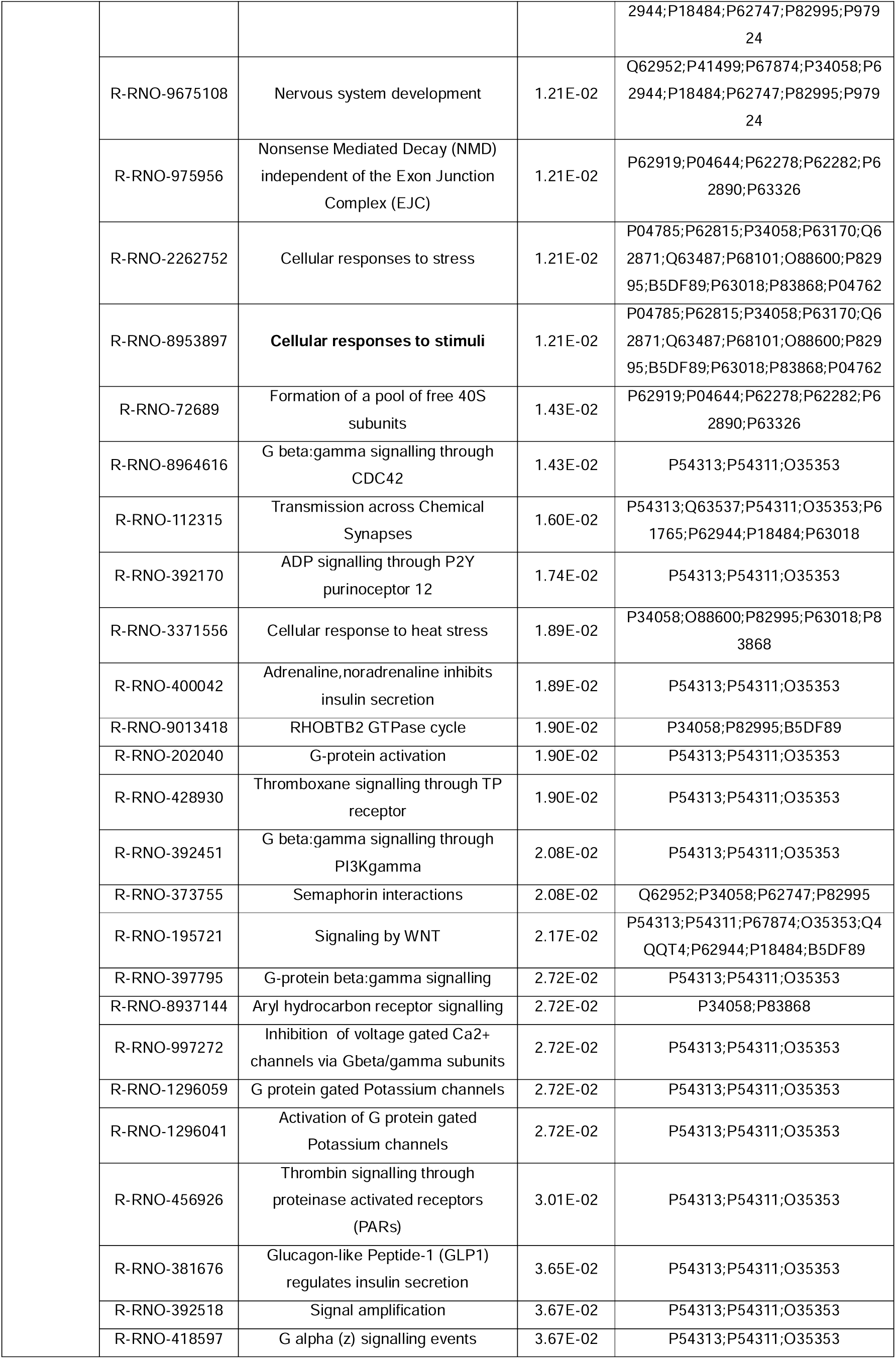
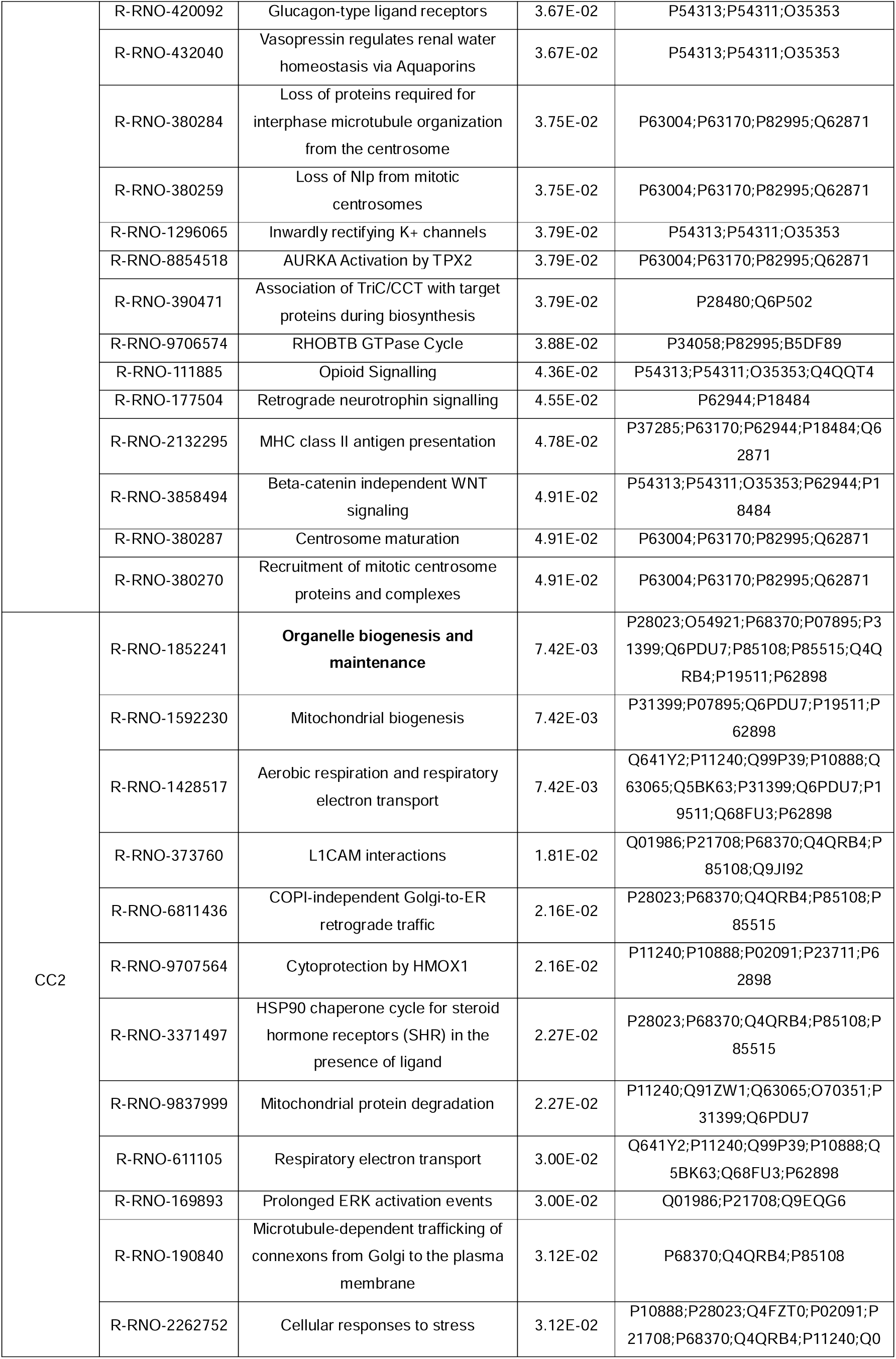
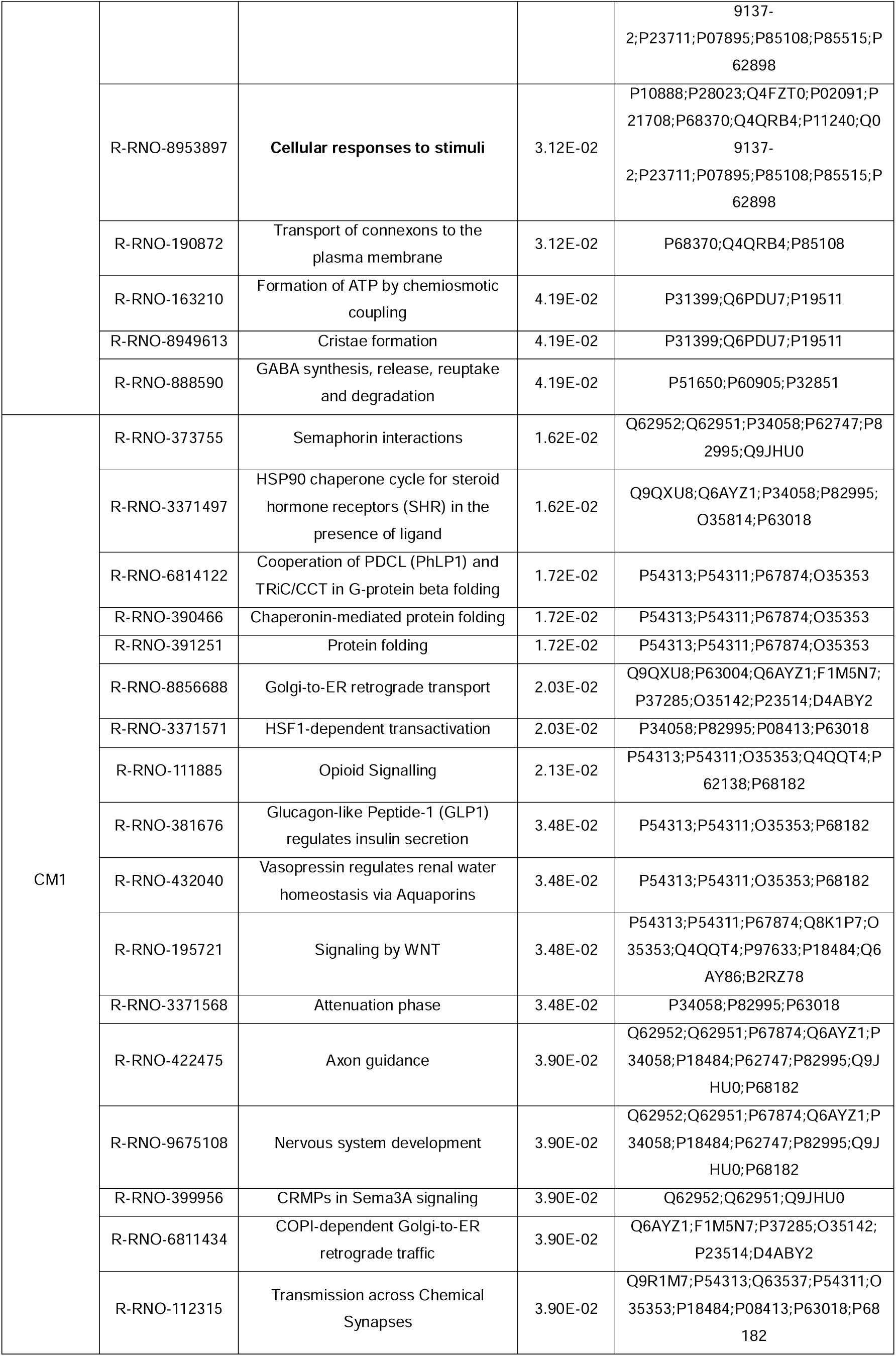
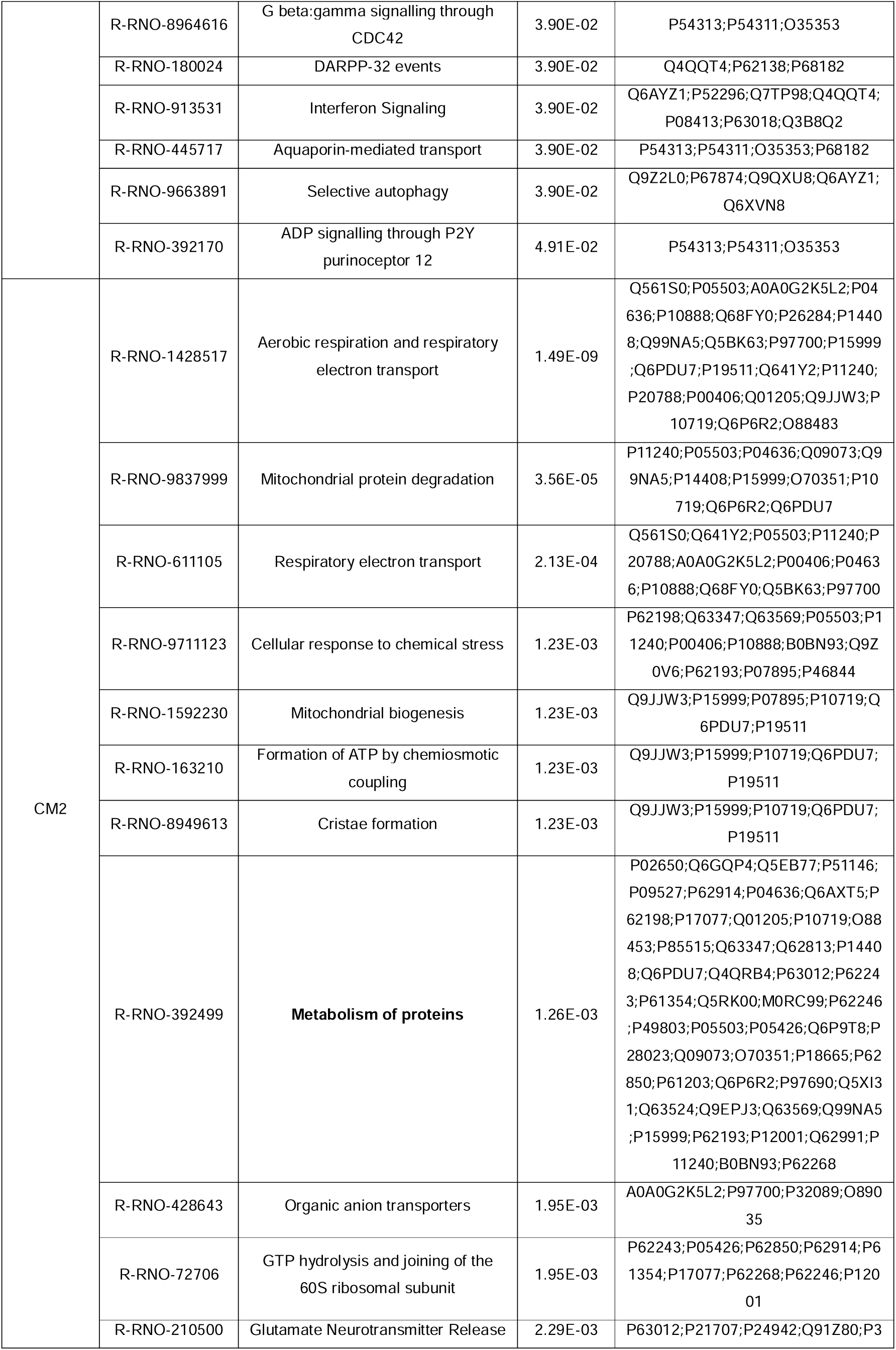
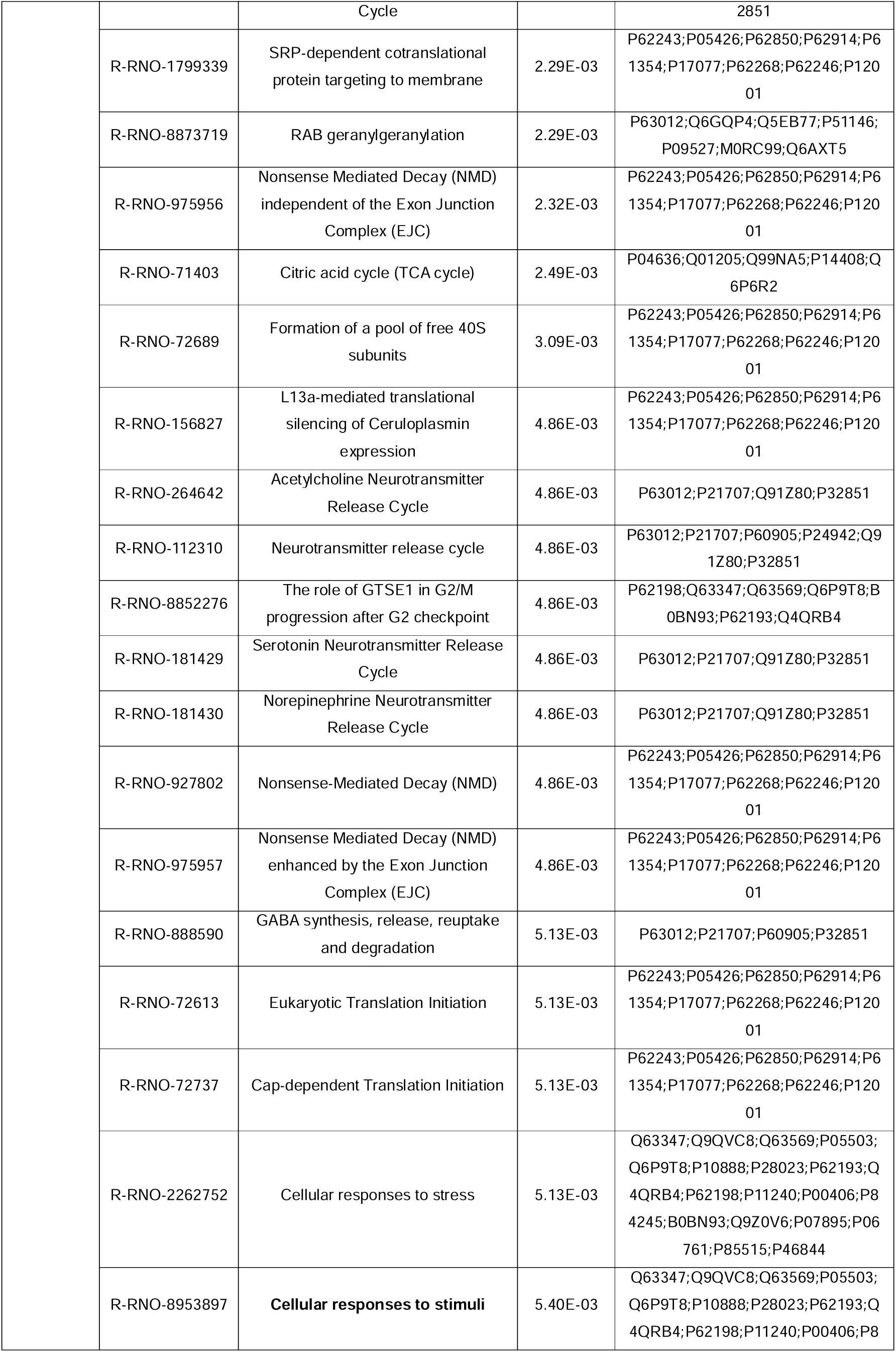
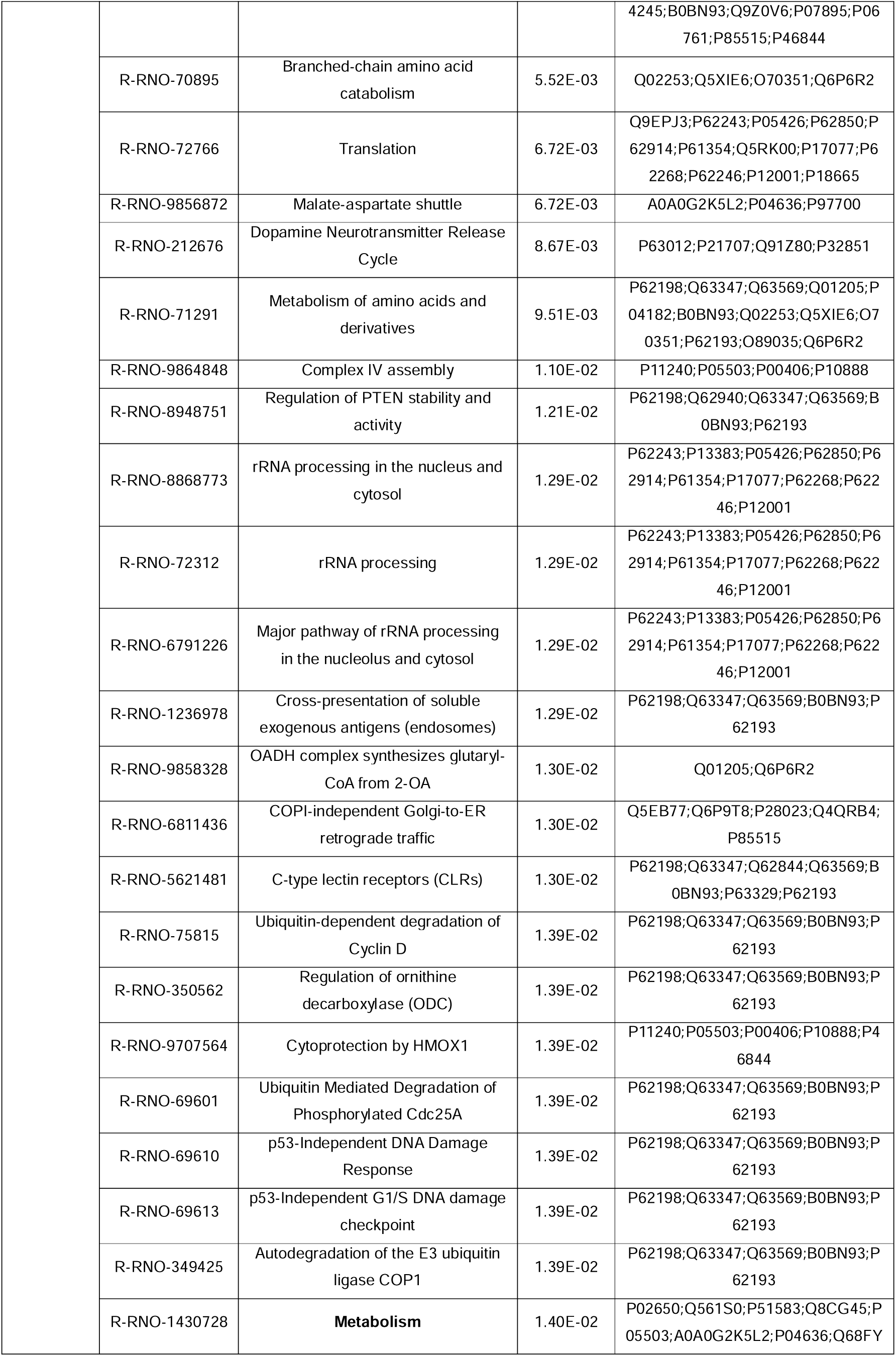
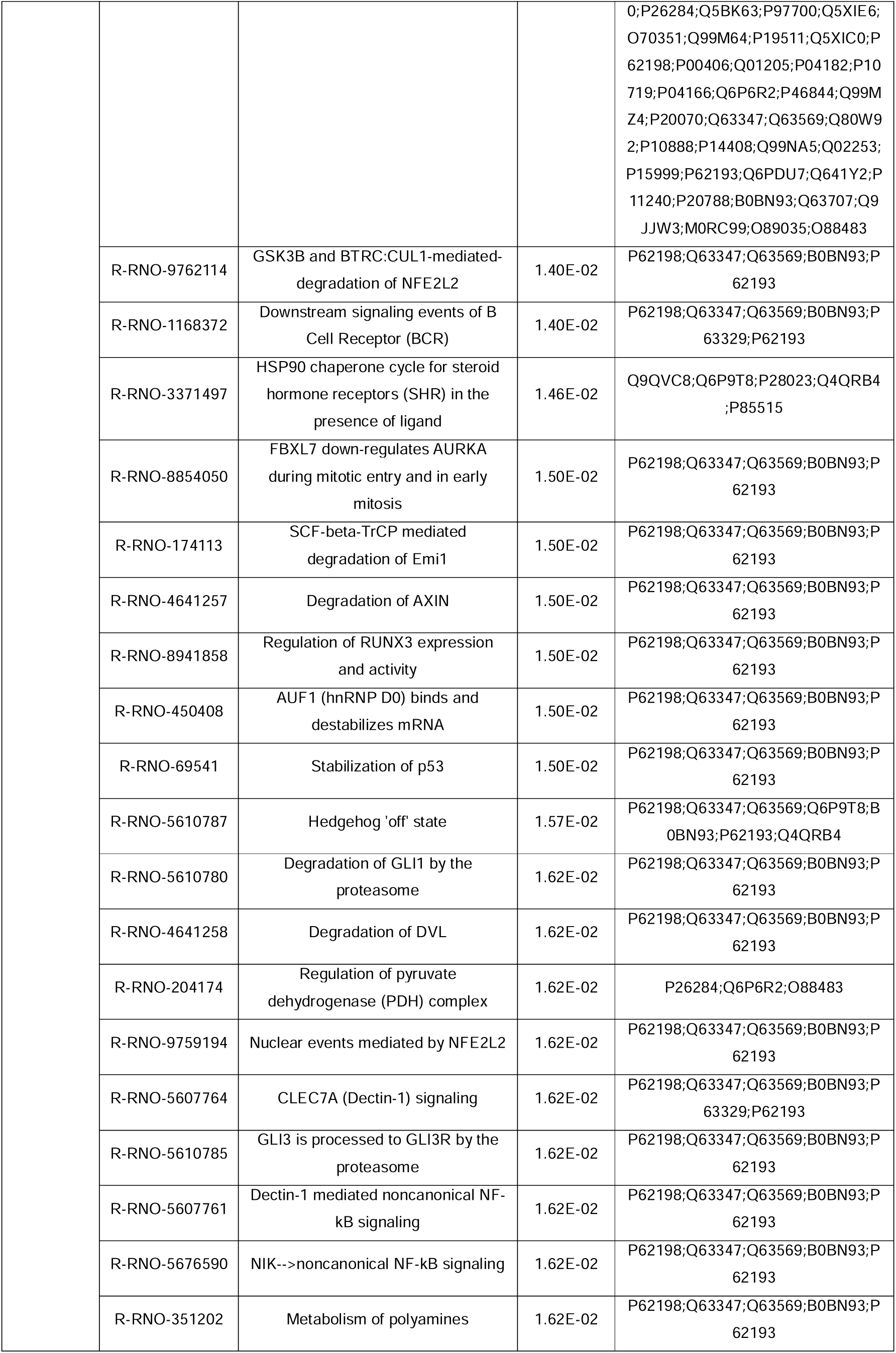
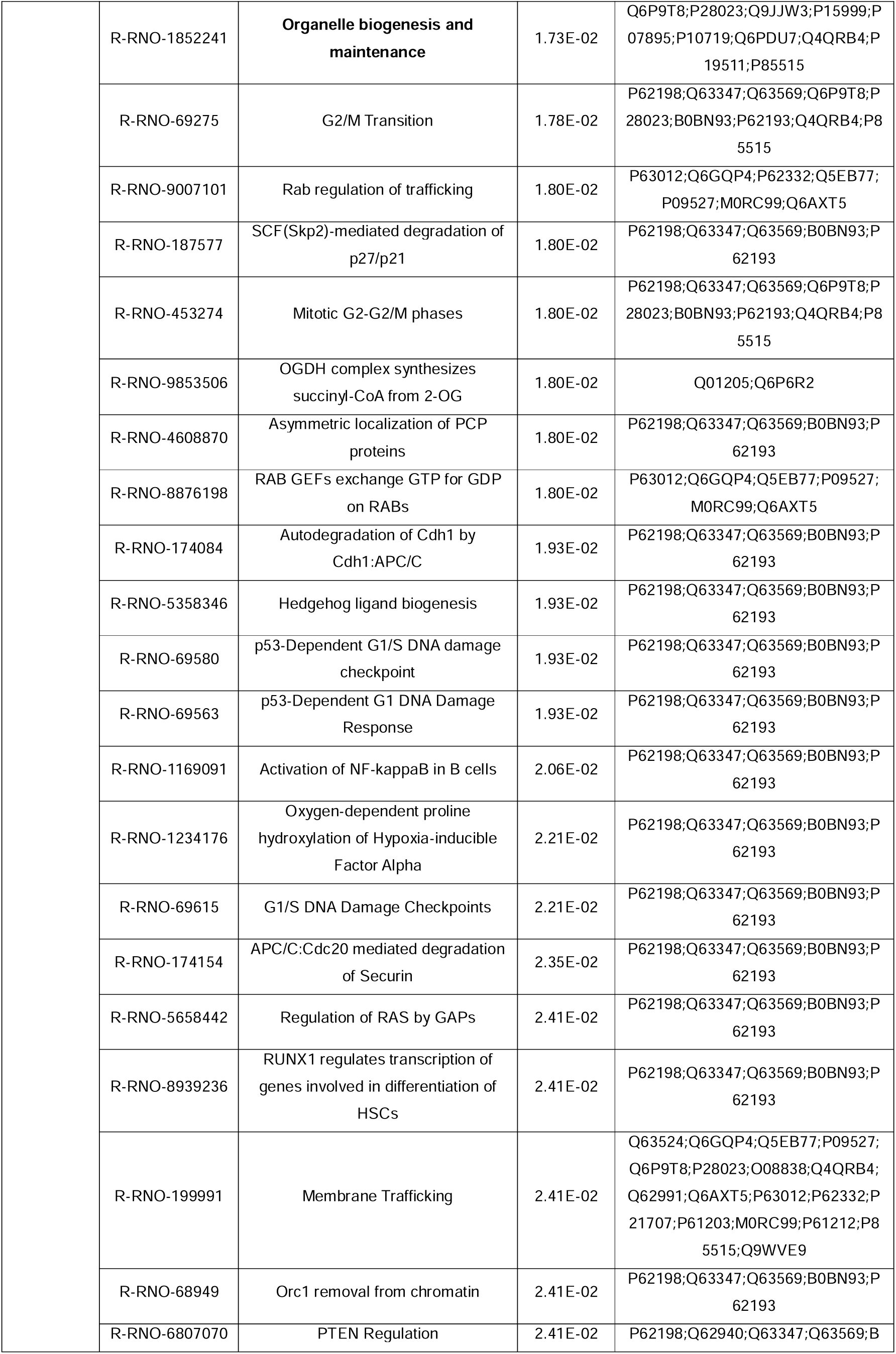
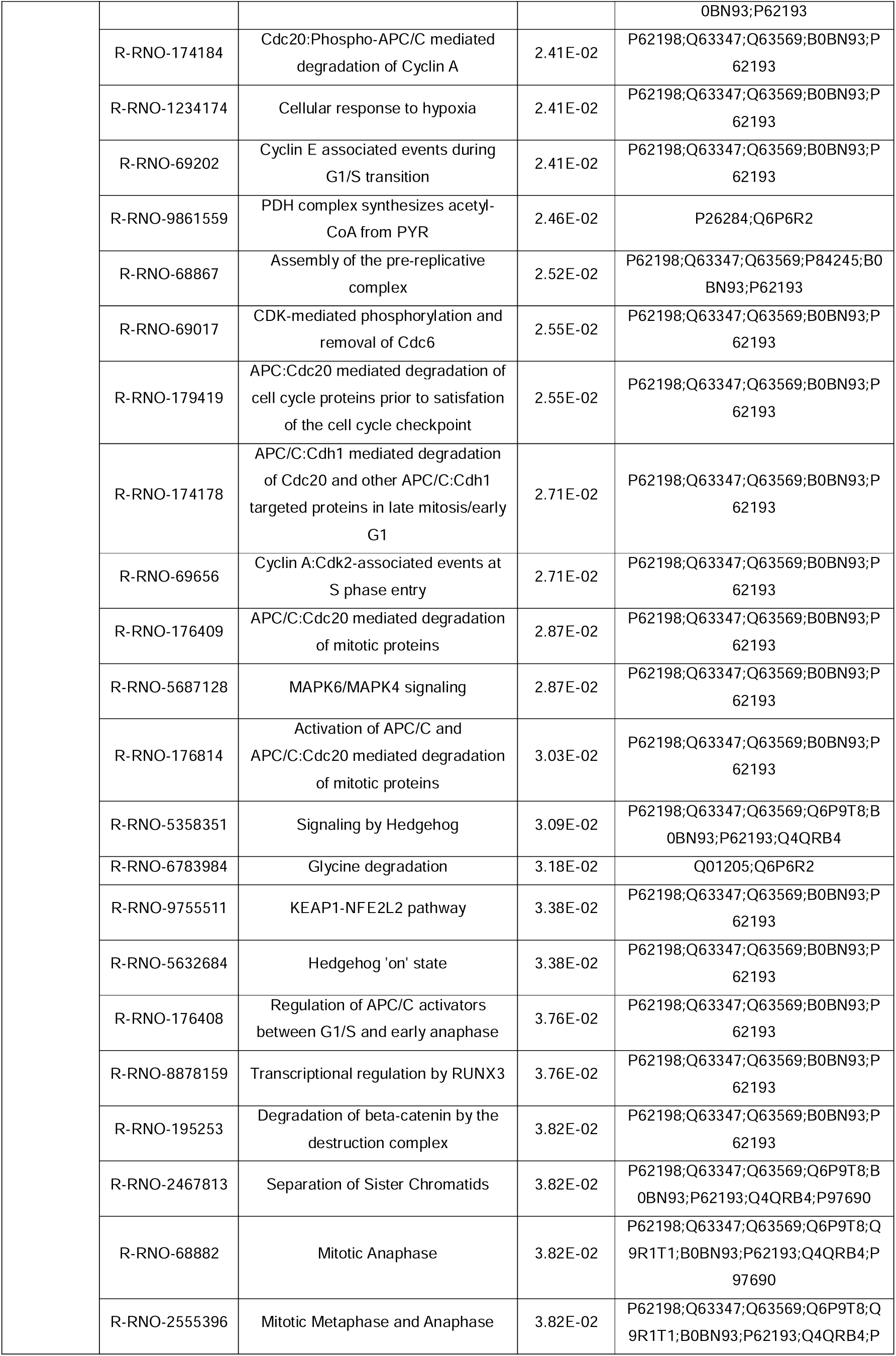
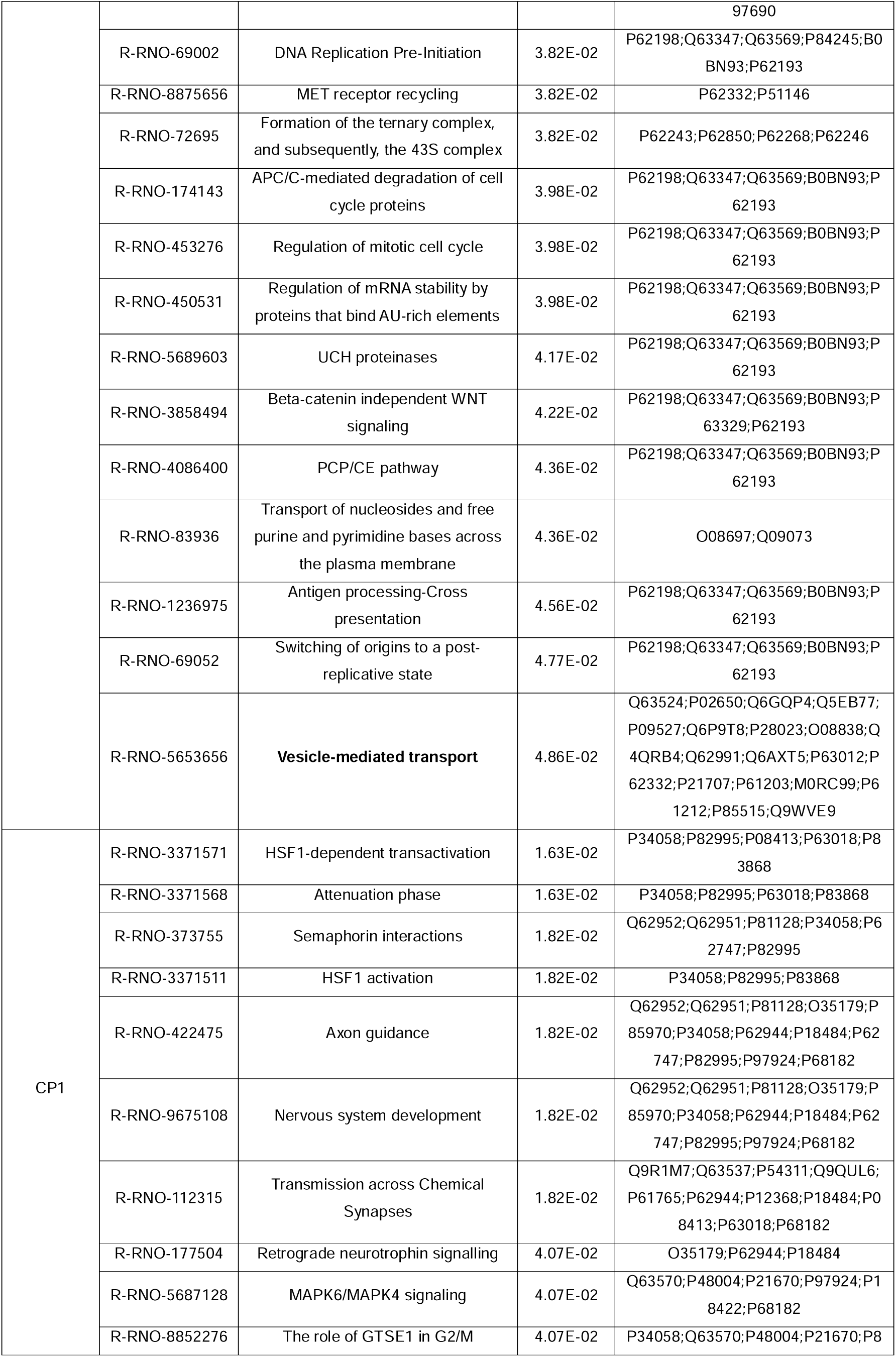
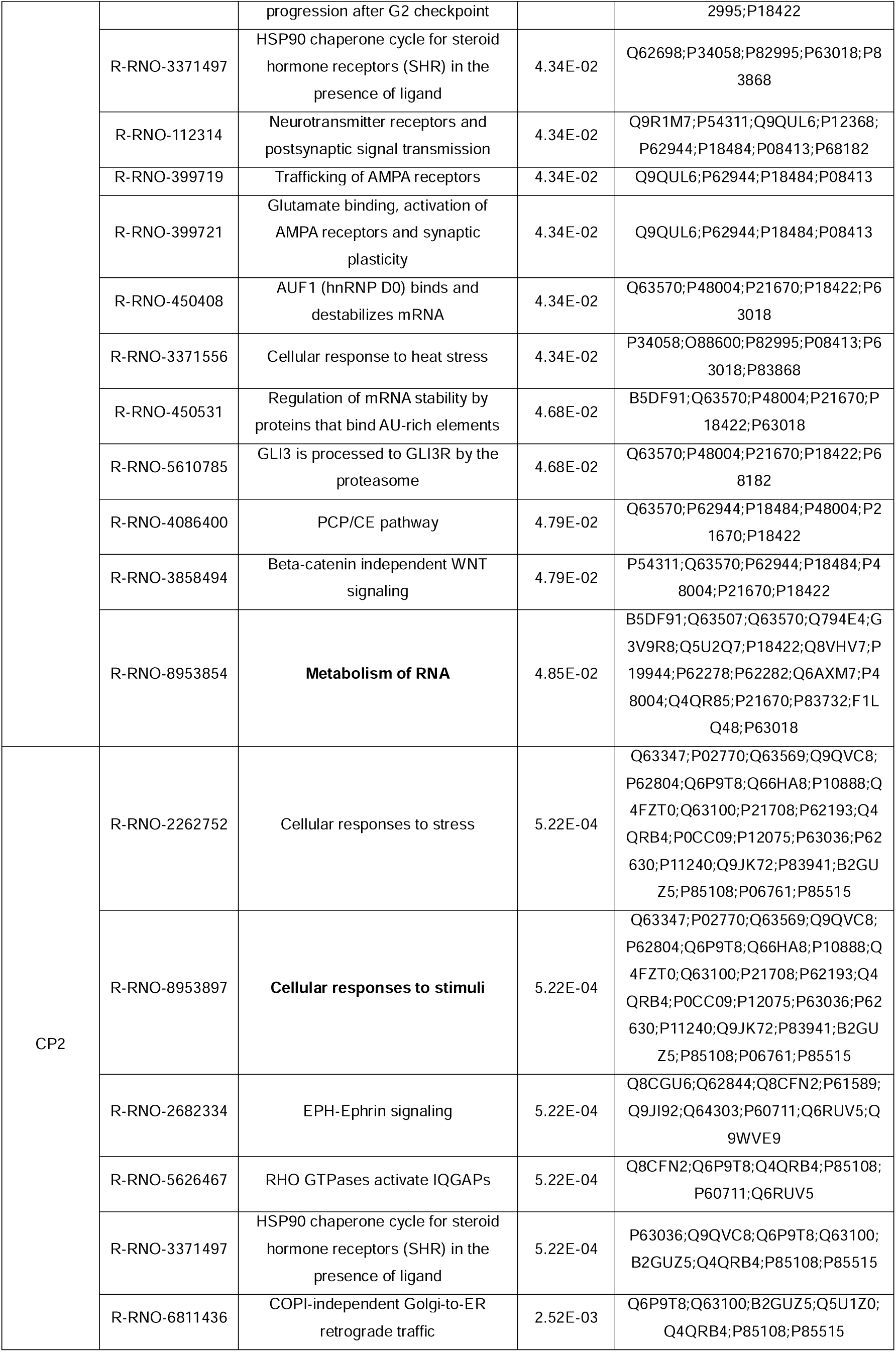
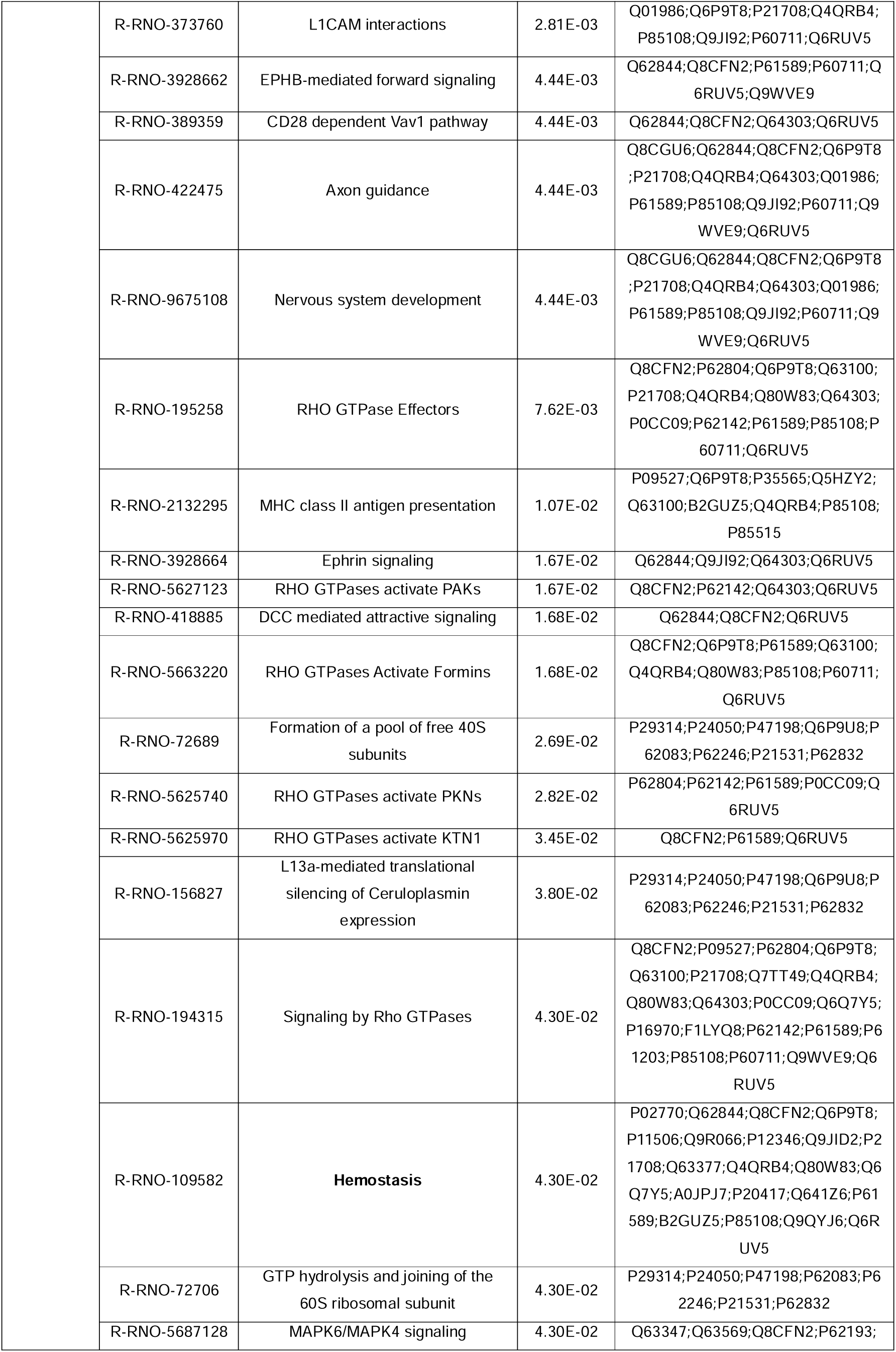
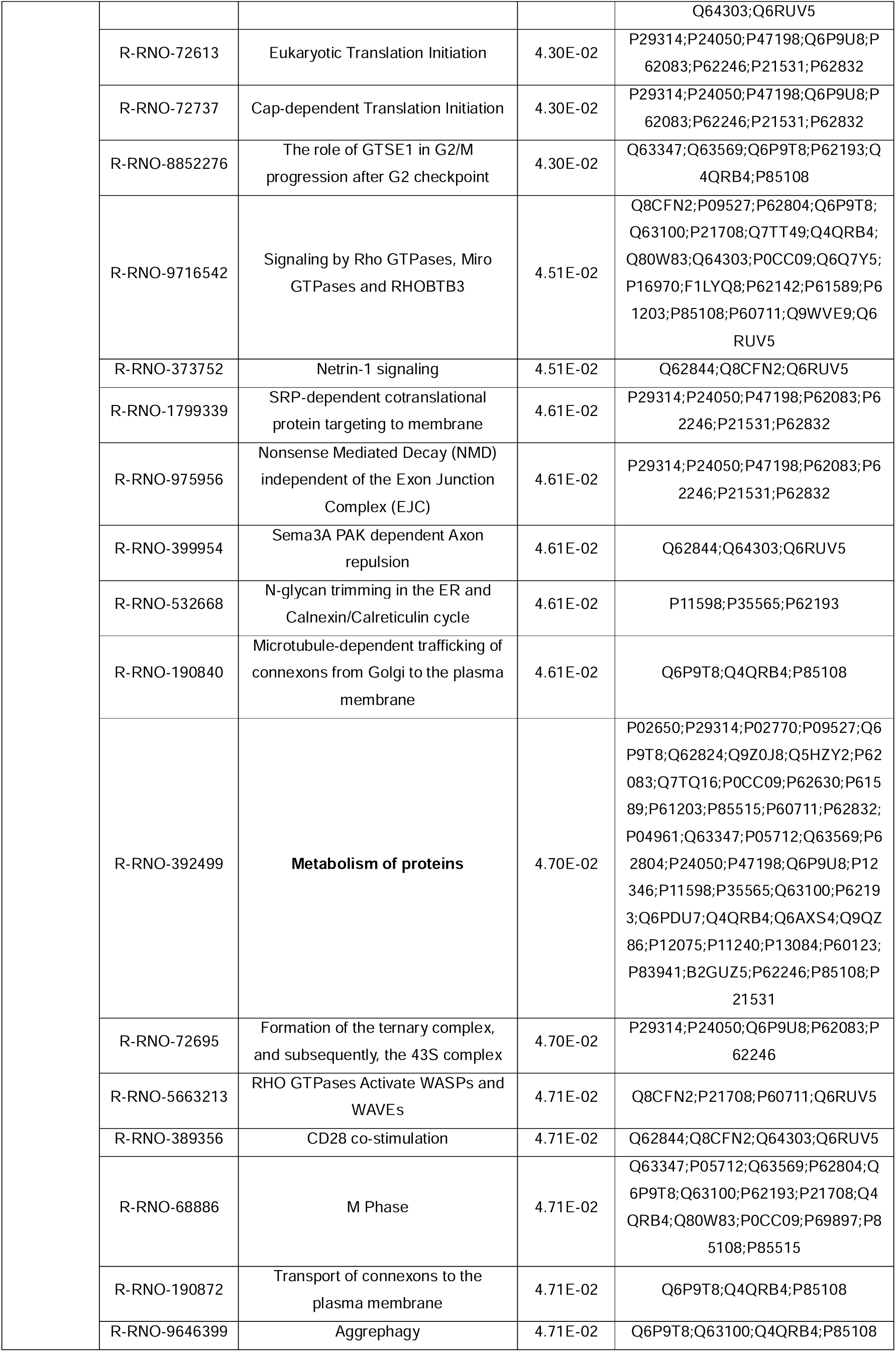
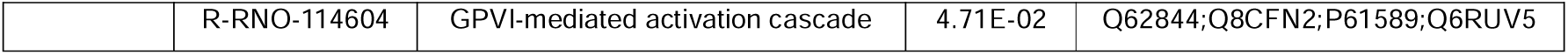
Reactome pathways of the selected clusters for neurons co-cultured with MSCs (Figure 2F).

**Supplementary Table 3.** Proteins identified using a univariate approach comparing sham and OGD co-cultured with conventional UC-MSCs.

| Accession | Name | Mann-Whitney<br>(Sham vs OGD) (p-value) | Mann-Whitney<br>(Sham vs<br>OGD+conventional<br>MSCs) (p-value) |
| --- | --- | --- | --- |
| Q00438 | Polypyrimidine tract-binding protein 1 | 0.032 | 0.286 |
| Q62638 | Golgi apparatus protein 1 | 0.019 | 0.413 |
| P11517 | *Hemoglobin subunit beta-2 | 0.032 | 0.413 |
| P14604 | Enoyl-CoA hydratase, mitochondrial | 0.016 | 0.110 |
| Q5I0K8 | 28S ribosomal protein S7, mitochondrial | 0.016 | 0.190 |
| Q66HF8 | Aldehyde dehydrogenase X, mitochondrial | 0.016 | 0.286 |
| P56574 | Isocitrate dehydrogenase [NADP],<br>mitochondrial | 0.016 | 0.063 |
| Q6AY23 | Pyrroline-5-carboxylate reductase 2 | 0.019 | 0.219 |
| B0K020 | *CDGSH iron-sulfur domain-containing<br>protein 1 | 0.016 | 0.730 |
| F1LPQ2 | Activating signal cointegrator 1 complex<br>subunit 3 | 0.016 | 0.413 |
| Q5XID7 | Armadillo repeat-containing X-linked protein<br>3 | 0.032 | 0.286 |
| Q6AXX6 | Peroxiredoxin-like 2A | 0.036 | 0.539 |
| P52020 | Squalene monooxygenase | 0.026 | 0.065 |
| P53676 | AP-3 complex subunit mu-1 | 0.016 | 0.190 |
| Q91XU8 | Phosphatidate cytidyltransferase 2 | 0.016 | 0.063 |
| Q68FU3 | *Electron transfer flavoprotein subunit beta | 0.016 | 0.413 |
| Q62812 | *Myosin-9 | 0.032 | 0.730 |
| P13803 | Electron transfer flavoprotein subunit alpha,<br>mitochondrial | 0.016 | 0.085 |
| Q641Y2 | *NADH dehydrogenase [ubiquinone] iron-sulfur protein 2, mitochondrial | 0.032 | 1.000 |
| P13233 | 2',3'-cyclic-nucleotide 3'-phosphodiesterase | 0.019 | 0.110 |
| P11884 | Aldehyde dehydrogenase, mitochondrial | 0.019 | 0.063 |
| Q498C8 | Protein RER1 | 0.016 | 0.111 |
| Q8CHN6 | Sphingosine-1-phosphate lyase 1 | 0.032 | 0.905 |
| P56522 | NADPH:adrenodoxin oxidoreductase,<br>mitochondrial | 0.016 | 0.063 |
| P10824 | Guanine nucleotide-binding protein G(i)<br>subunit alpha-1 | 0.032 | 0.325 |
| P97852 | Peroxisomal multifunctional enzyme type 2 | 0.016 | 0.063 |
| Q64LC9 | *RNA-binding protein 4B | 0.027 | 1.000 |
| Q63617 | Hypoxia up-regulated protein 1 | 0.032 | 0.065 |
| P48721 | Stress-70 protein, mitochondrial | 0.016 | 0.413 |
| Q5SGE0 | Leucine-rich PPR motif-containing protein,<br>mitochondrial | 0.032 | 0.286 |
| Q9WUL0 | DNA topoisomerase 1 | 0.016 | 0.085 |
| P20788 | Cytochrome b-c1 complex subunit Rieske,<br>mitochondrial | 0.016 | 0.111 |
| P62997 | Transformer-2 protein homolog beta | 0.016 | 0.063 |
| Q5D006 | Ubiquitin carboxyl-terminal hydrolase 11 | 0.016 | 0.063 |
| Q09137 | *5'-AMP-activated protein kinase catalytic<br>subunit alpha-2 | 0.032 | 0.905 |
| P37199 | Nuclear pore complex protein Nup155 | 0.032 | 0.111 |
| Q641Y0 | Dolichyl-diphosphooligosaccharide--protein<br>glycosyltransferase 48 kDa subunit | 0.016 | 0.138 |
| P17764 | Acetyl-CoA acetyltransferase, mitochondrial | 0.049 | 0.190 |
| P97532 | 3-mercaptopyruvate sulfurtransferase | 0.049 | 0.413 |
| Q3KRE0 | ATPase family AAA domain-containing<br>protein 3 | 0.049 | 0.140 |
| Q5PPL3 | Sterol-4-alpha-carboxylate 3-dehydrogenase,<br>decarboxylating | 0.016 | 0.174 |
| P15999 | ATP synthase subunit alpha, mitochondrial | 0.032 | 0.413 |
| P68136 | Actin, alpha skeletal muscle | 0.019 | 0.190 |
| Q80ZG1 | Synembryn-A | 0.016 | 0.176 |
| Q01205 | Dihydrolipoylysine-residue<br>succinyltransferase component of 2-oxoglutarate dehydrogenase complex,<br>mitochondrial | 0.027 | 0.325 |
| Q8K4K5 | Lethal(2) giant larvae protein homolog 1 | 0.016 | 0.063 |
| B5DEH2 | *Erlin-2 | 0.016 | 0.806 |
| Q63633 | Solute carrier family 12 member 5 | 0.048 | 0.325 |
| P85834 | Elongation factor Tu, mitochondrial | 0.032 | 0.556 |
| Q7TPJ0 | Translocon-associated protein subunit alpha | 0.049 | 0.176 |
| P10719 | ATP synthase subunit beta, mitochondrial | 0.034 | 0.286 |
| O35458 | Vesicular inhibitory amino acid transporter | 0.016 | 0.286 |
| Q5XIH7 | Prohibitin-2 | 0.032 | 0.063 |
| P63039 | 60 kDa heat shock protein, mitochondrial | 0.019 | 0.110 |
| F1LYQ8 | FERM, ARHGEF and pleckstrin domain-containing protein 1 | 0.049 | 0.730 |
| Q06486 | Casein kinase I isoform delta | 0.016 | 0.111 |
| P67779 | *Prohibitin 1 | 0.019 | 1.000 |
| Q56B11 | Proline-, glutamic acid- and leucine-rich<br>protein 1 | 0.026 | 0.176 |
| Q99ML5 | Prenylcysteine oxidase 1 | 0.032 | 0.111 |
| P38718 | Mitochondrial pyruvate carrier 2 | 0.016 | 0.190 |
| Q8CG45 | Aflatoxin B1 aldehyde reductase member 2 | 0.032 | 0.556 |
| Q812E9 | Neuronal membrane glycoprotein M6-a | 0.016 | 0.730 |
| P97700 | Mitochondrial 2-oxoglutarate/malate carrier<br>protein | 0.016 | 0.730 |
| P30835 | ATP-dependent 6-phosphofructokinase, liver<br>type | 0.016 | 0.190 |
| O88600 | +Heat shock 70 kDa protein 4 | 0.019 | 1.000 |
| Q63487 | +Ras-related GTP-binding protein B | 0.034 | 0.461 |
| Q07266 | Drebrin | 0.016 | 0.286 |
| O55096 | +Dipeptidyl peptidase 3 | 0.032 | 0.413 |
| D3ZD32 | +Chromodomain-helicase-DNA-binding protein 5 | 0.019 | 0.805 |
| P62716 | Serine/threonine-protein phosphatase 2A catalytic subunit beta isoform | 0.016 | 0.110 |
| Q08163 | Adenylyl cyclase-associated protein 1 | 0.016 | 0.190 |
| Q7TP47 | Heterogeneous nuclear ribonucleoprotein Q | 0.016 | 0.063 |
| Q6AYK8 | Eukaryotic translation initiation factor 3 subunit D | 0.049 | 0.286 |
| Q9JMJ4 | Pre-mRNA-processing factor 19 | 0.019 | 0.111 |
| P07943 | Aldo-keto reductase family 1 member B1 | 0.037 | 0.063 |
| P23514 | Coatomer subunit beta | 0.019 | 0.111 |
| Q80U96 | Exportin-1 | 0.016 | 0.063 |
| P61765 | +Syntaxin-binding protein 1 | 0.032 | 0.190 |
| Q9ESW0 | +DNA damage-binding protein 1 | 0.037 | 0.385 |
| Q9JK11 | Reticulon-4 | 0.016 | 0.111 |
| P49088 | Asparagine synthetase [glutamine-hydrolyzing] | 0.032 | 0.111 |
| P0DJJ3 | SH3-containing GRB2-like protein 3-interacting protein 1 | 0.032 | 0.190 |
| P82995 | +Heat shock protein HSP 90-alpha | 0.016 | 0.623 |
| P55053 | Fatty acid-binding protein 5 | 0.016 | 0.140 |
| Q64537 | Calpain small subunit 1 | 0.037 | 0.063 |
| Q71UF4 | Histone-binding protein RBBP7 | 0.019 | 0.063 |
| P19527 | Neurofilament light polypeptide | 0.032 | 0.063 |
| O70593 | Small glutamine-rich tetratricopeptide repeat-containing protein alpha | 0.016 | 0.190 |
| Q9R080 | +G-protein-signaling modulator 1 | 0.016 | 1.000 |
| B0BN85 | Protein SGT1 homolog | 0.016 | 0.063 |
| P20651 | Serine/threonine-protein phosphatase 2B catalytic subunit beta isoform | 0.027 | 0.085 |
| P38062 | Methionine aminopeptidase 2 | 0.016 | 0.063 |
| Q5XIM7 | Lysine--tRNA ligase | 0.032 | 0.286 |
| Q3SWU3 | Heterogeneous nuclear ribonucleoprotein D-like | 0.034 | 0.268 |
| Q7TP98 | Interleukin enhancer-binding factor 2 | 0.016 | 0.190 |
| + for proteins present on cluster CC1; * for proteins present on cluster CC2 |  |  |  |

**Supplementary Table 4.**
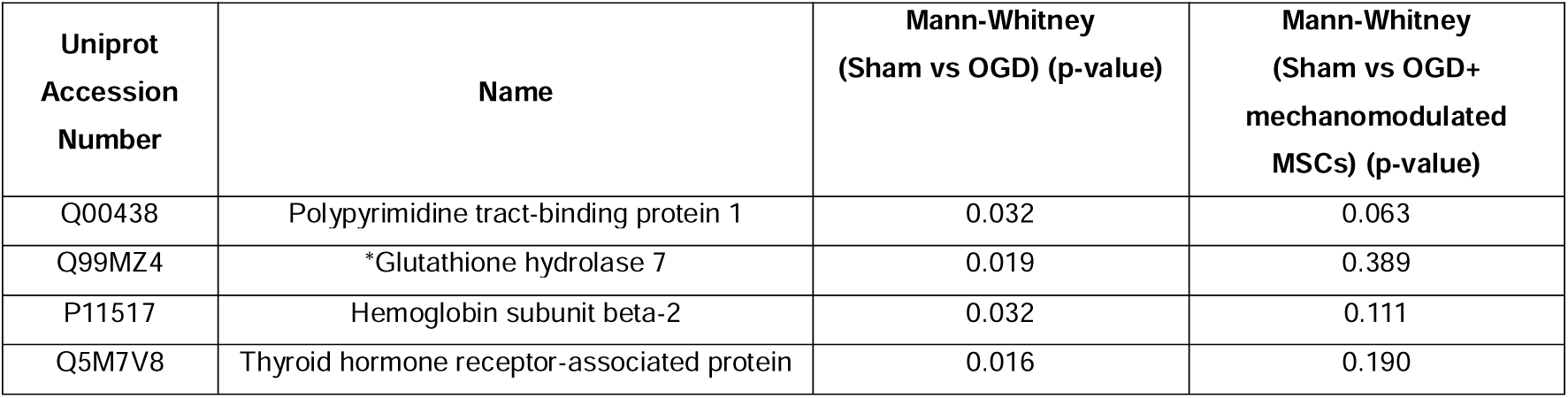
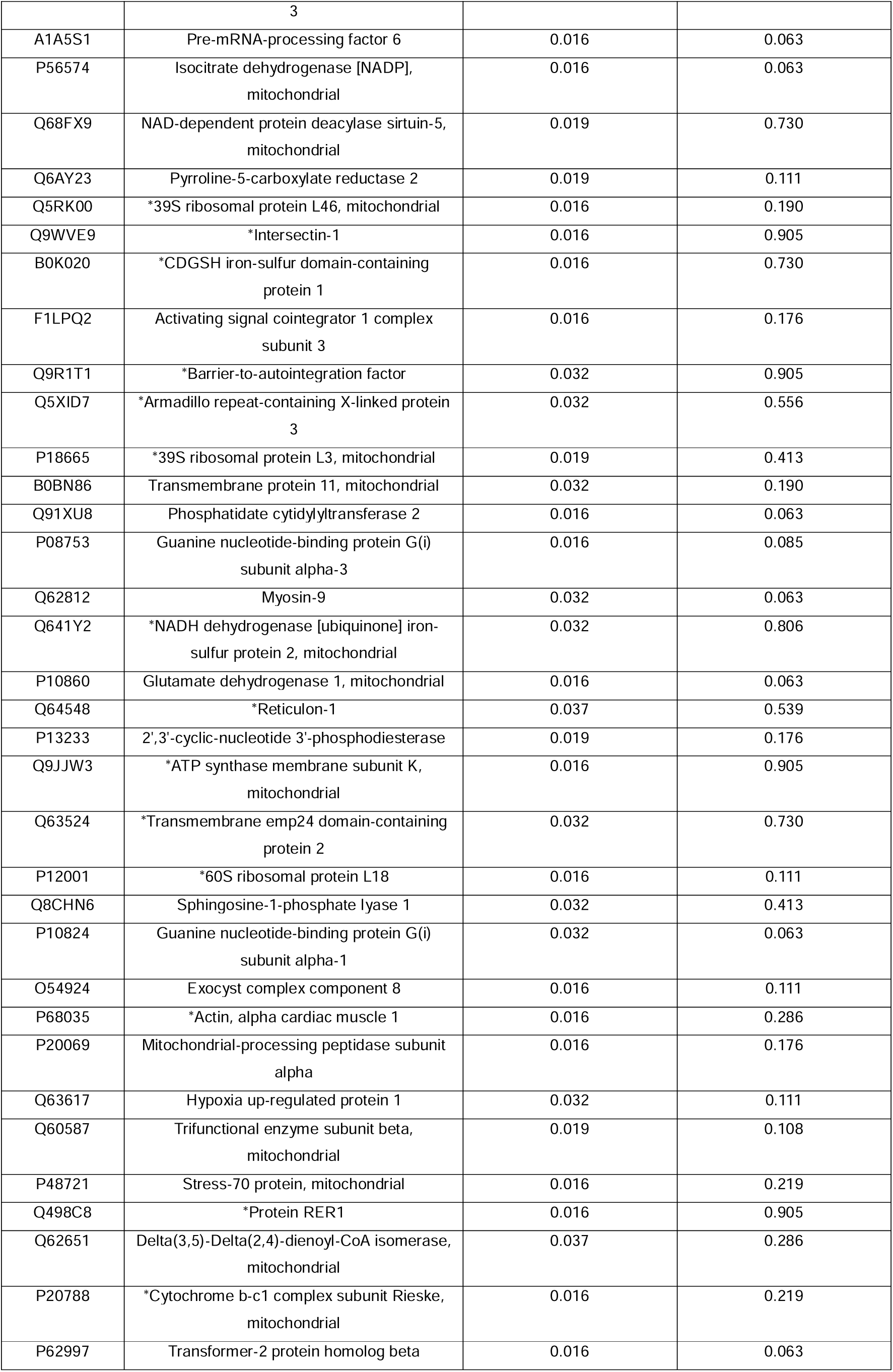
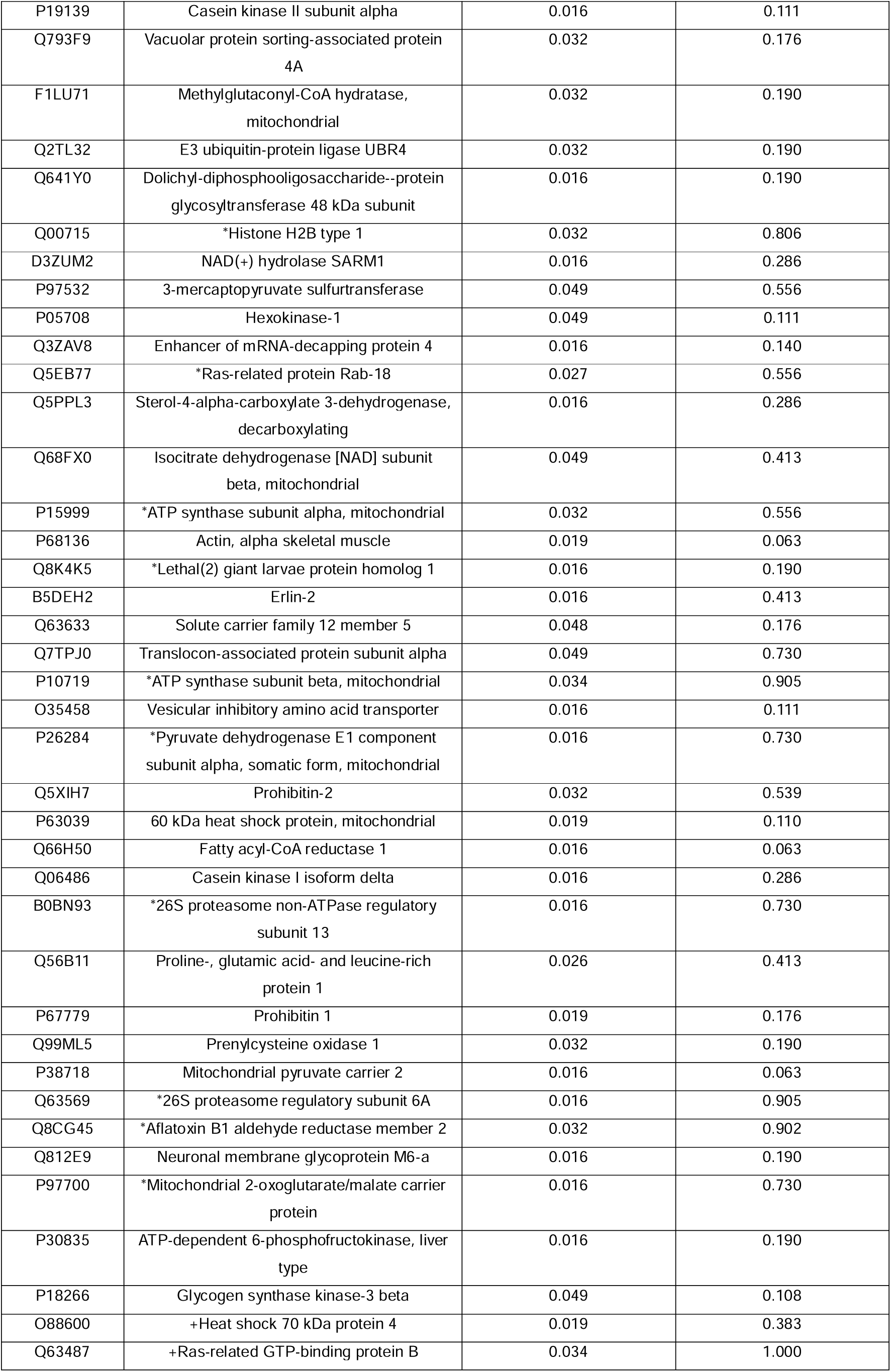
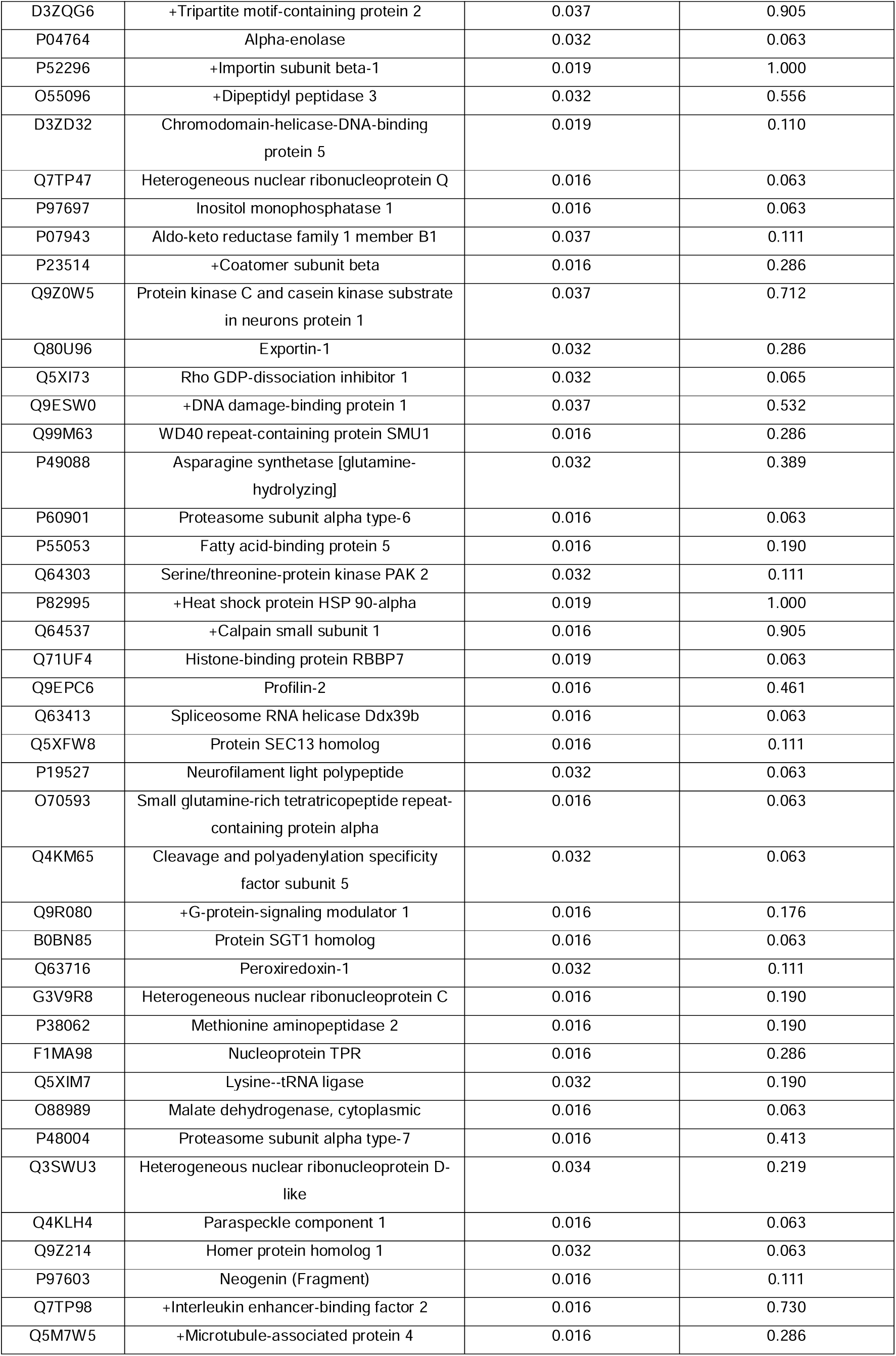
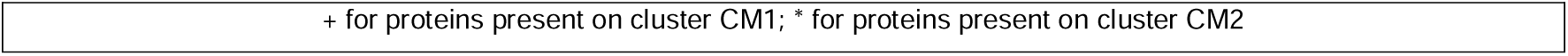
Proteins identified using a univariate approach comparing sham and OGD co-cultured with mechano-modulated UC-MSCs.

**Supplementary Table 5.**
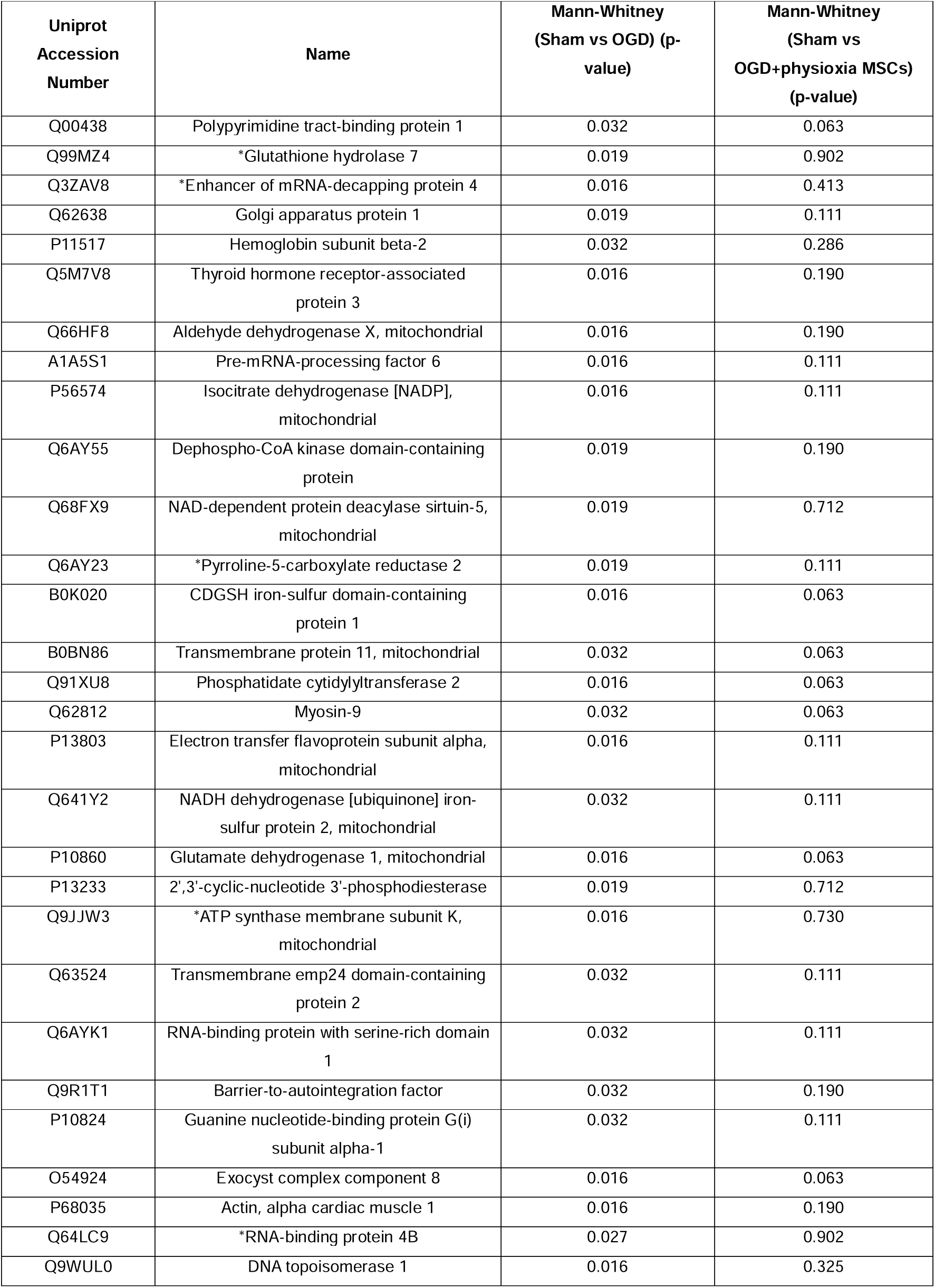
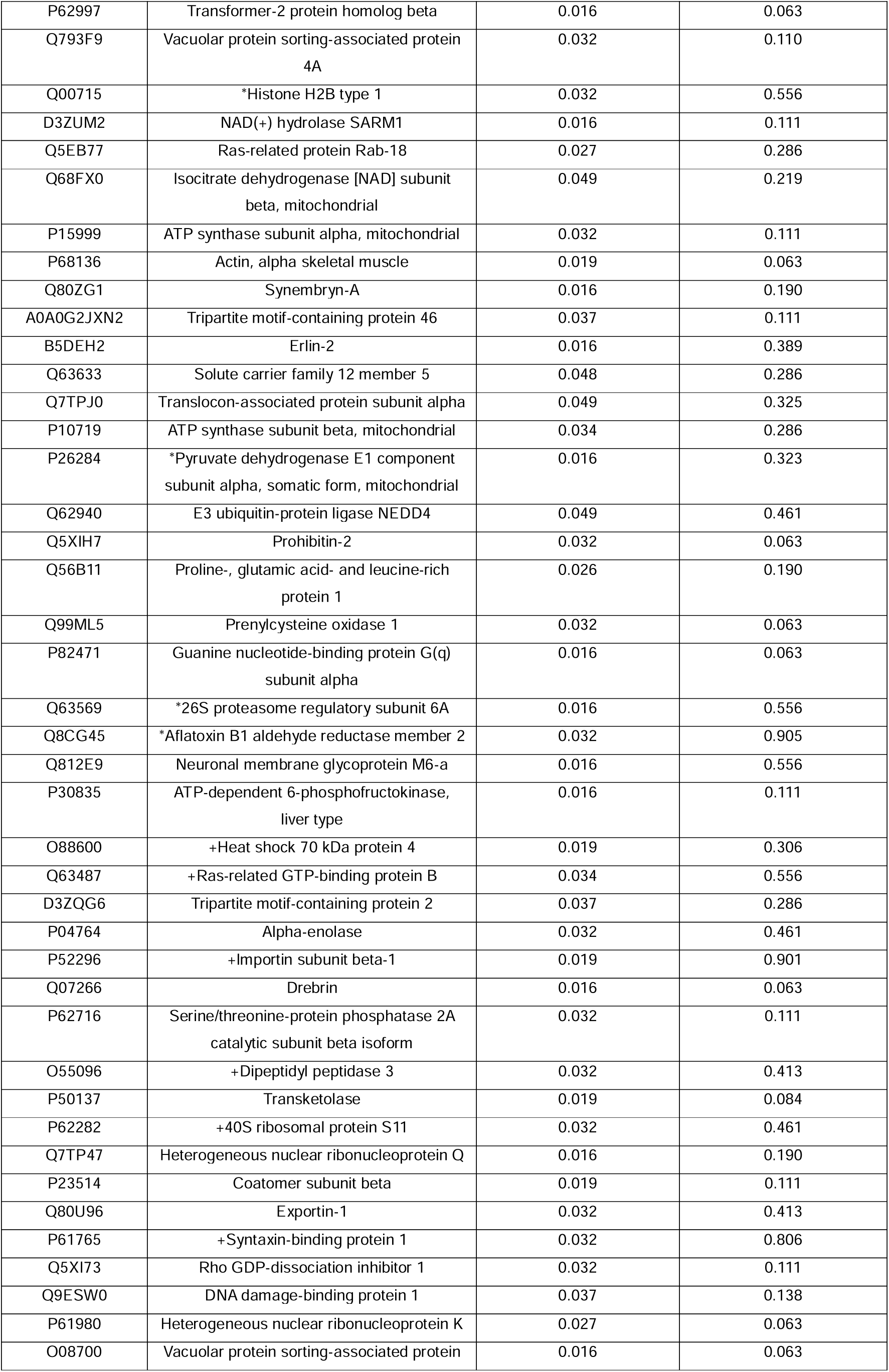
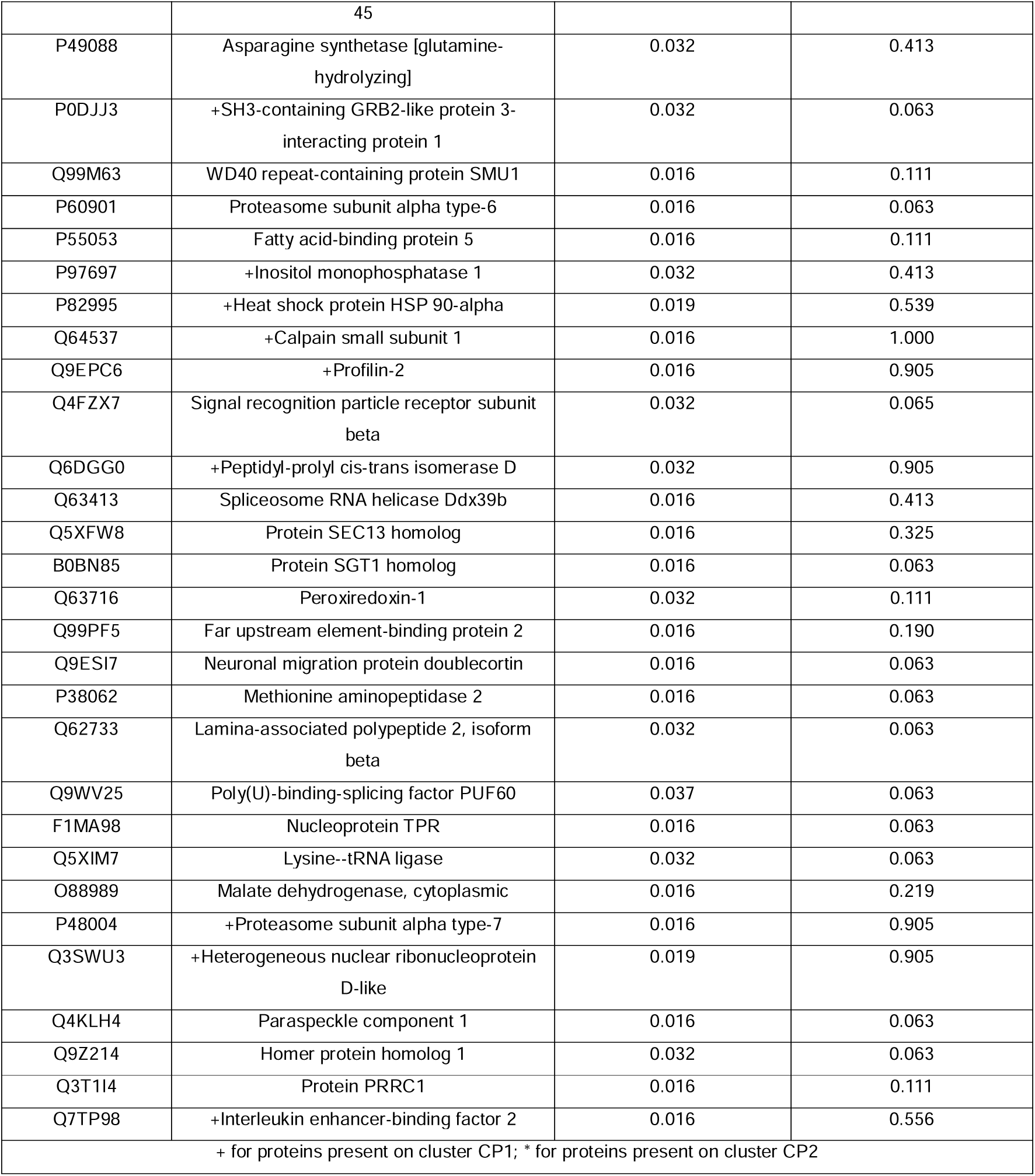
Proteins identified using a univariate approach comparing sham and OGD co-cultured with physioxia-modulated UC-MSCs.

**Supplementary Table 6.**
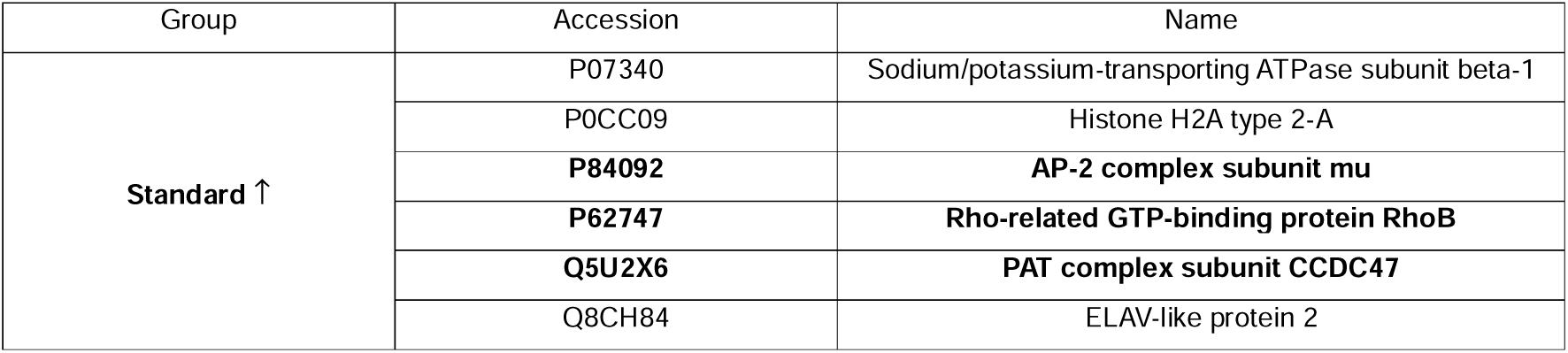
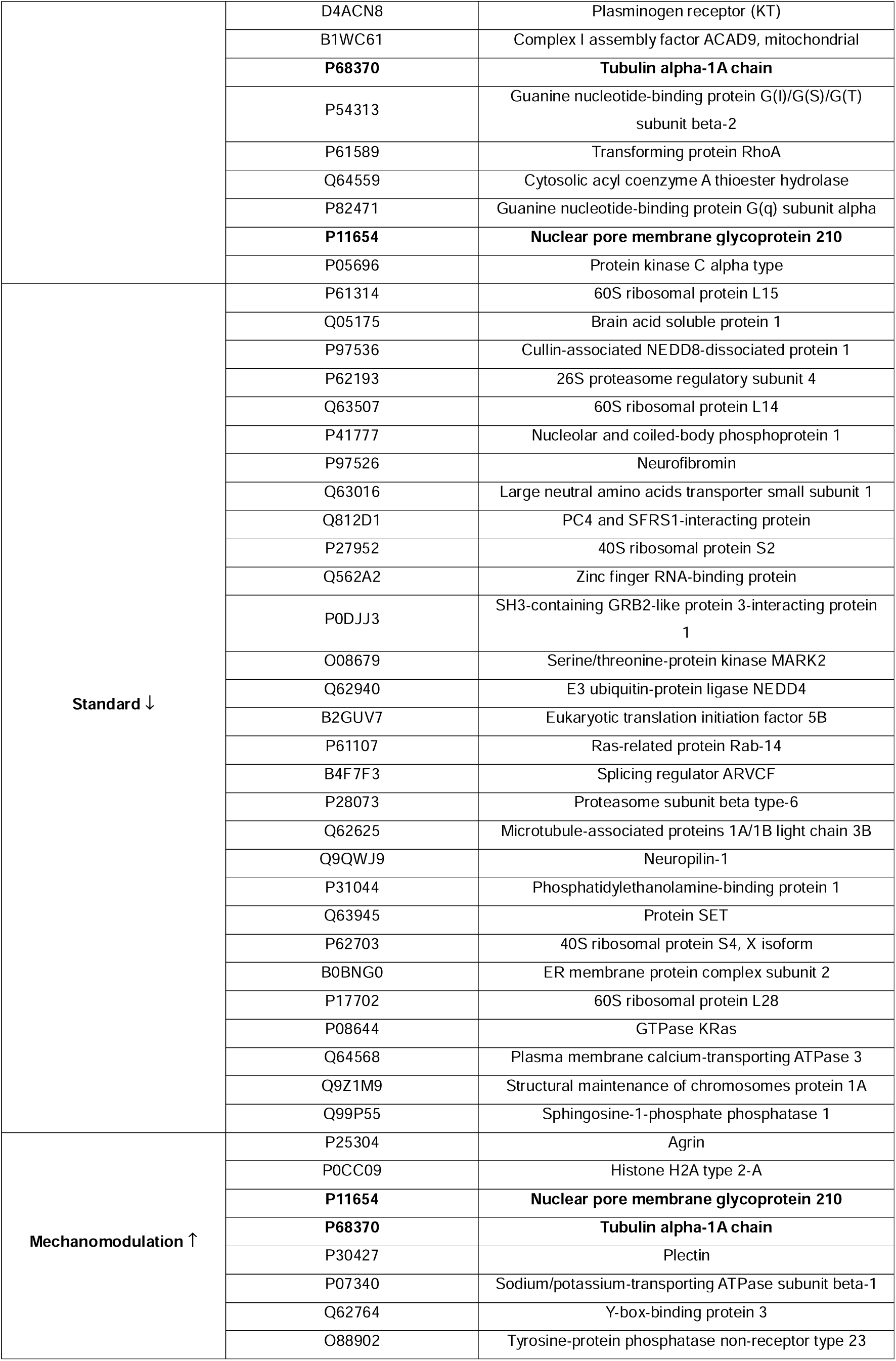
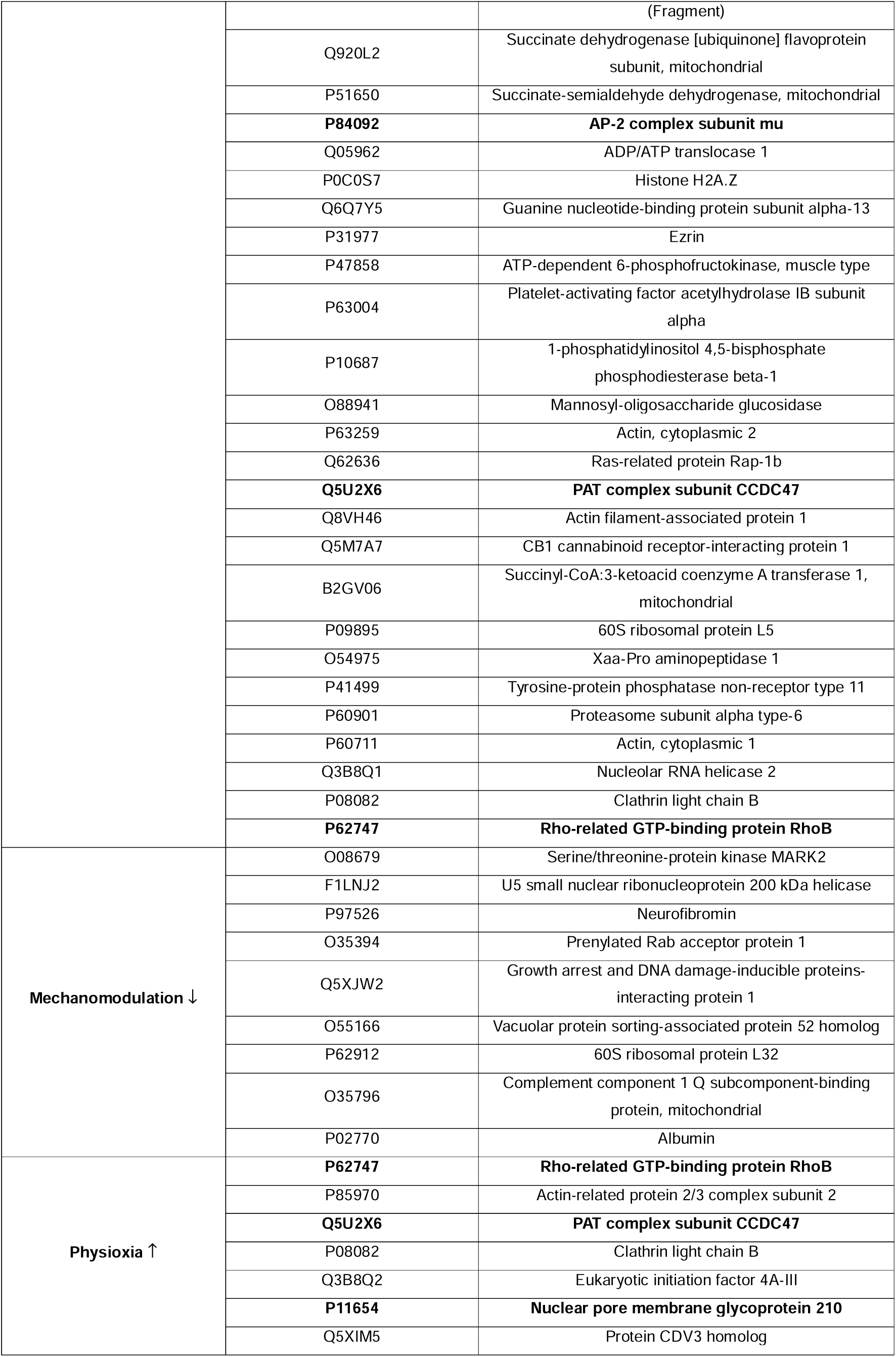
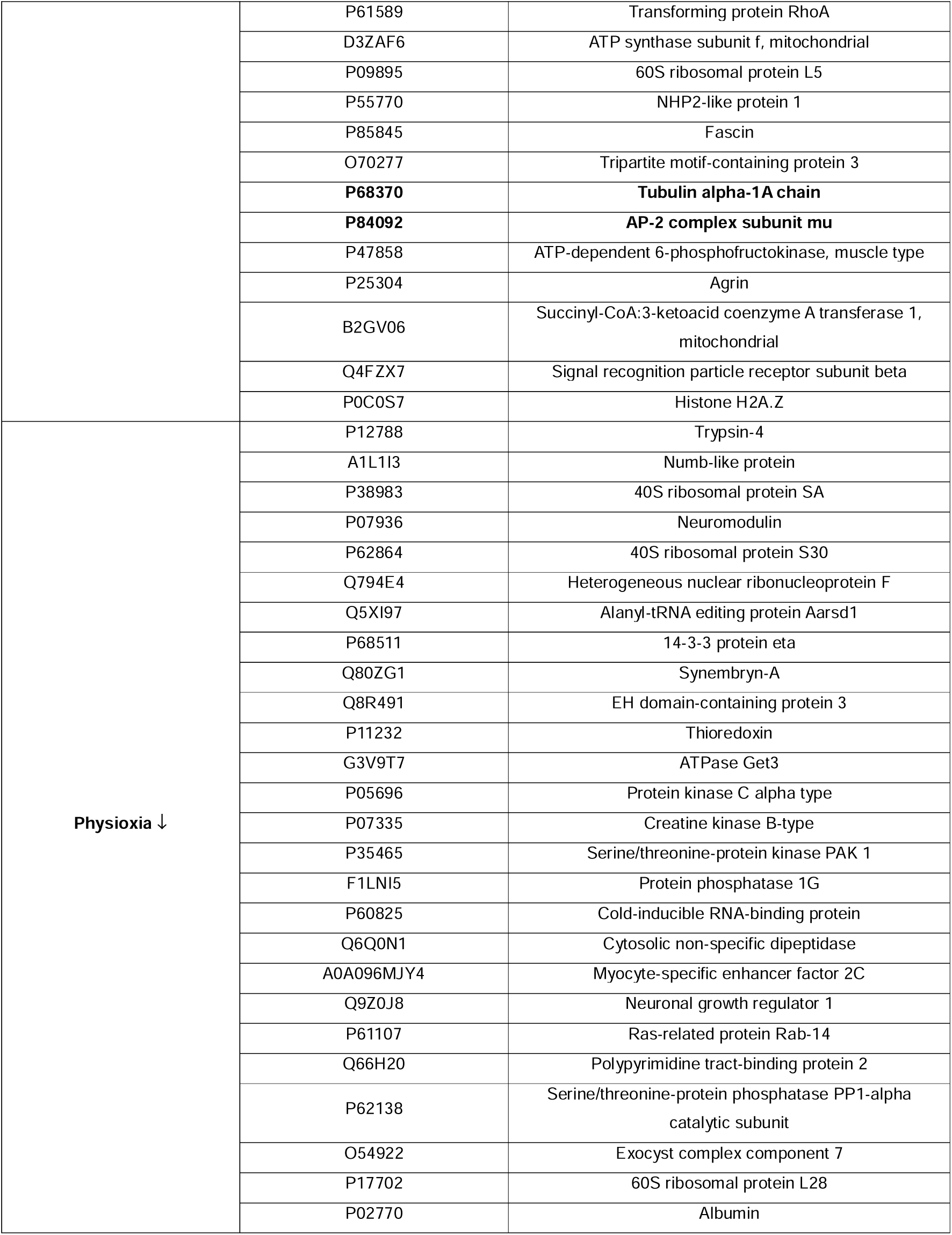
Proteins altered on neurons after the OGD insult and with the addition of modulated UC-MSCs’ secretome (represented on Venn, Supplementary Figure 5B).

**Supplementary Table 7.**
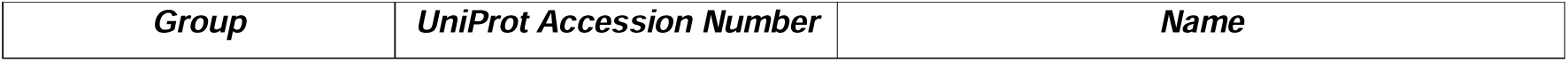
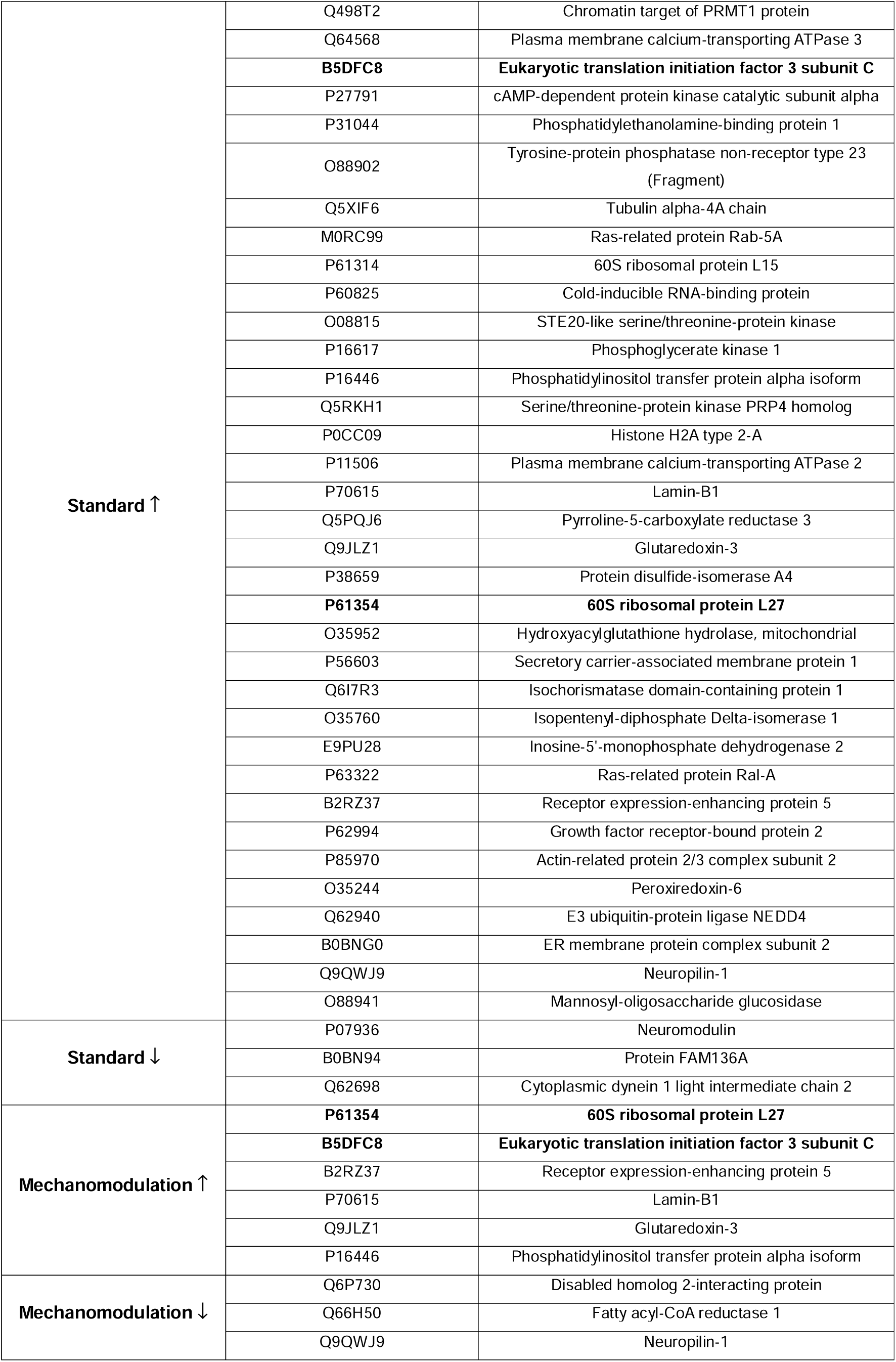
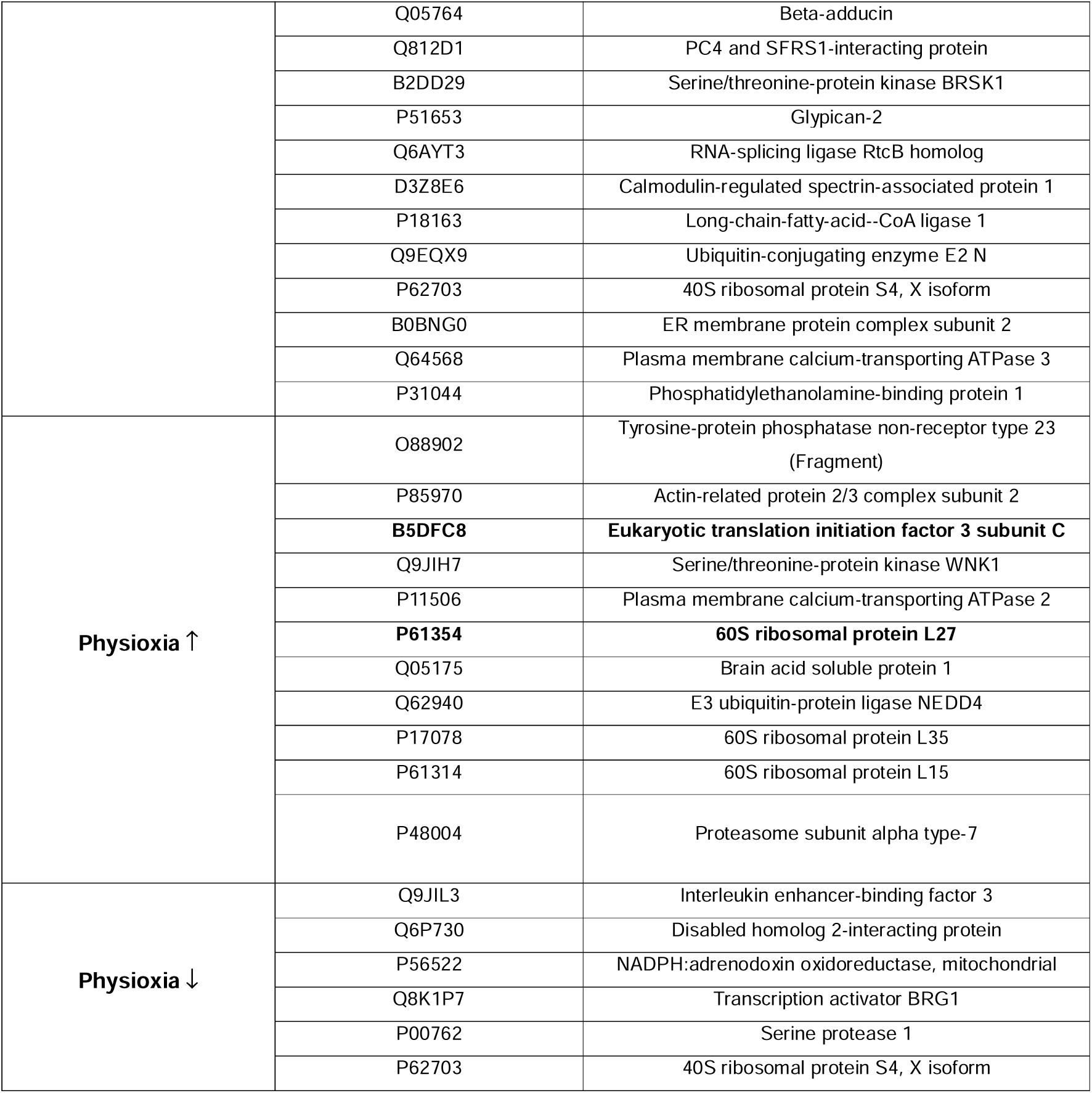
Proteins altered on sham neurons upon the addition of modulated UC-MSCs’ secretome (represented on Venn, Supplementary Figure 5F)

**Supplementary Table 8.**
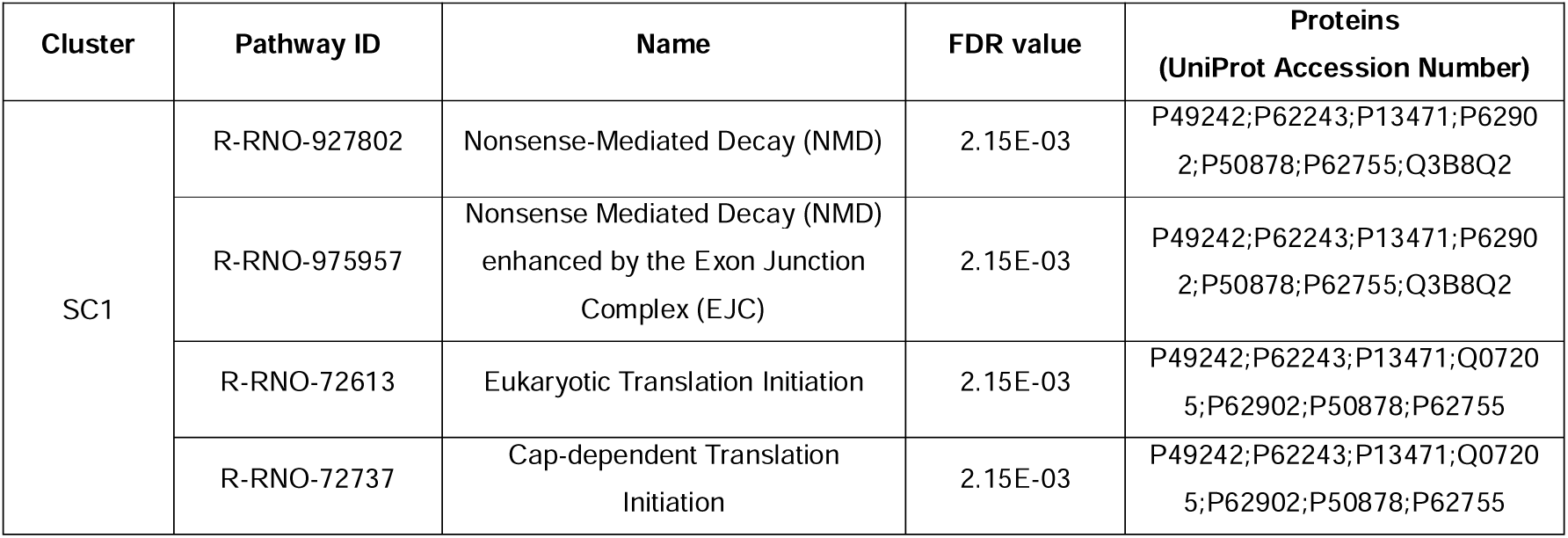
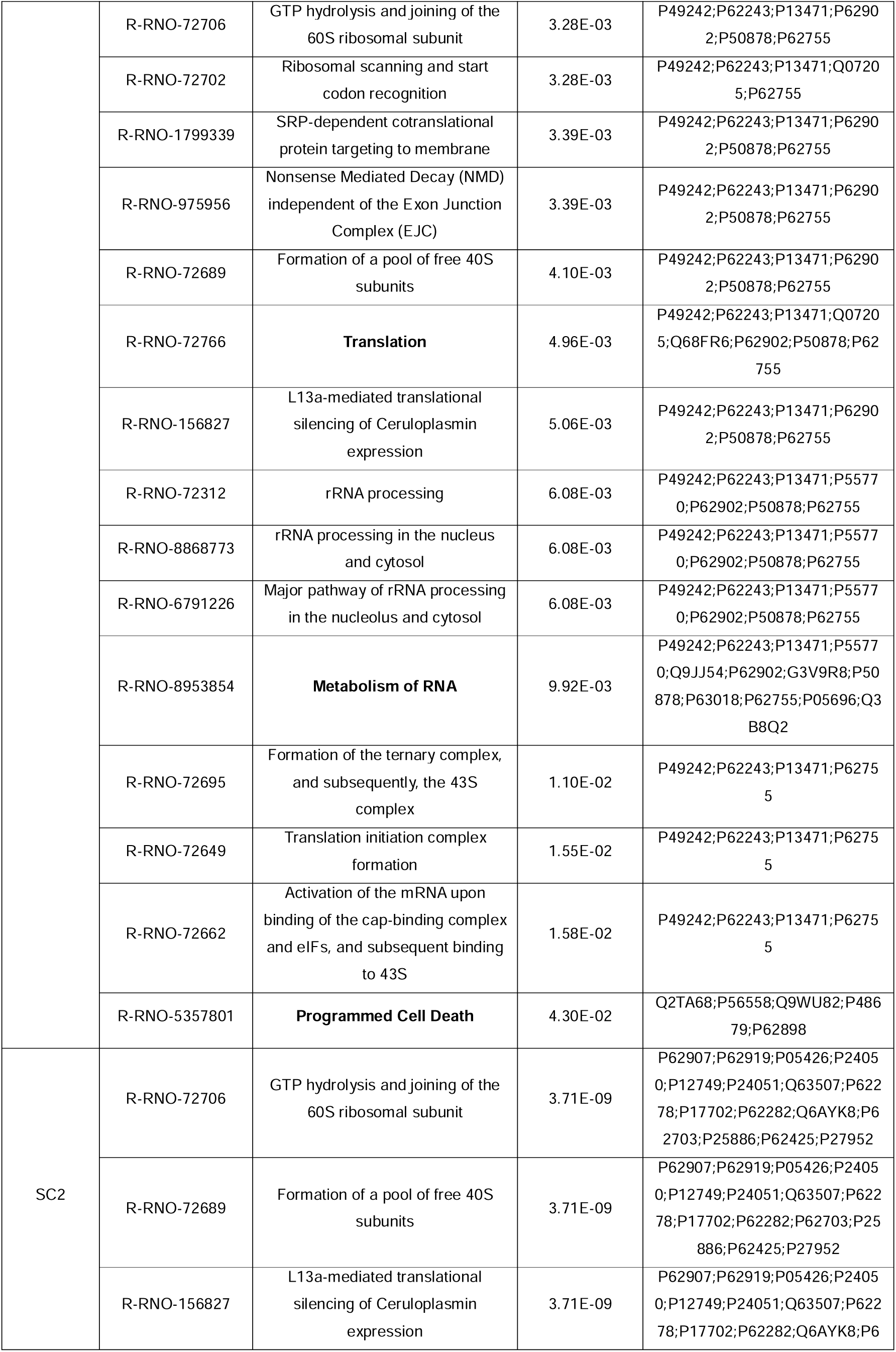
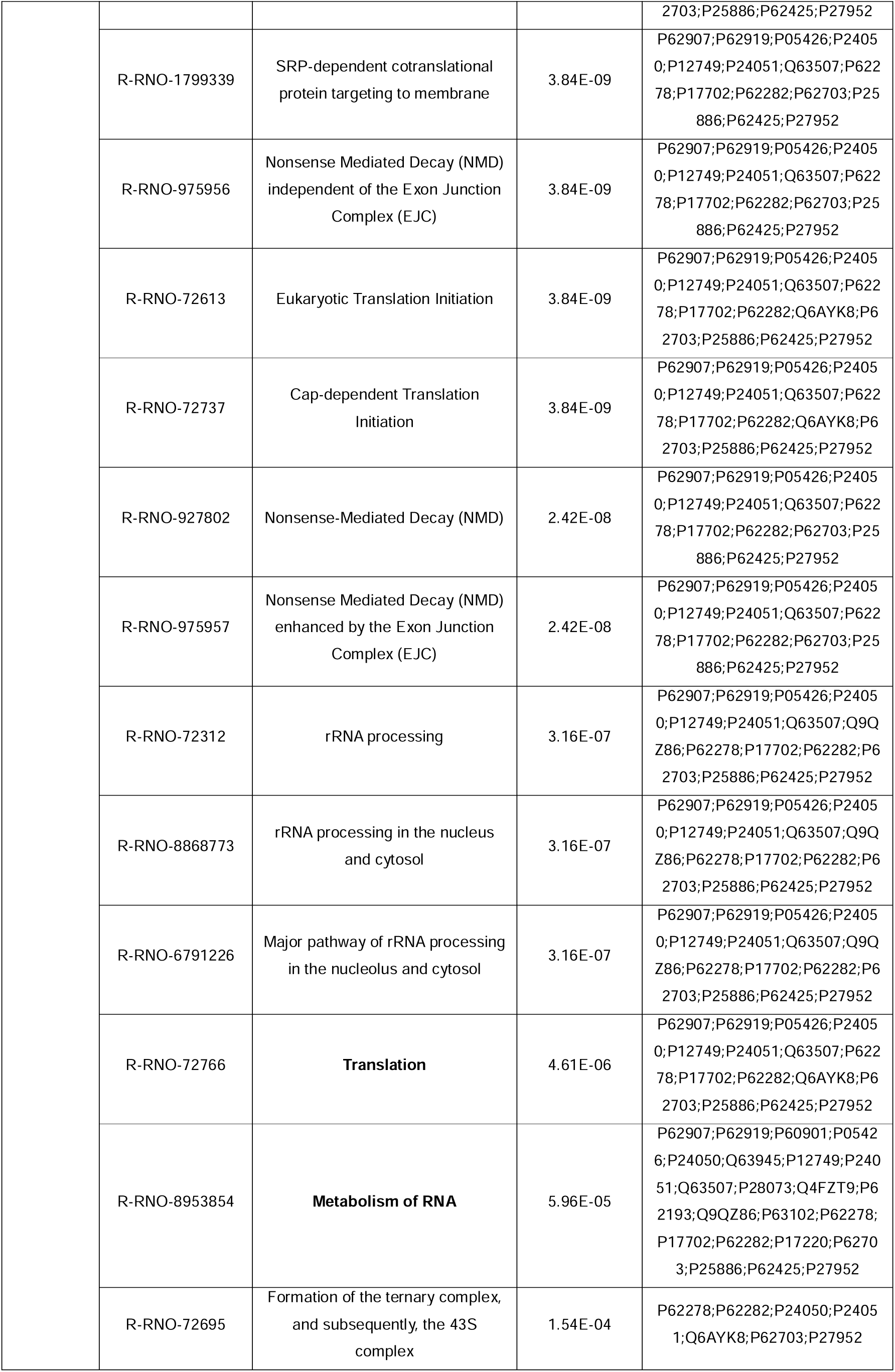
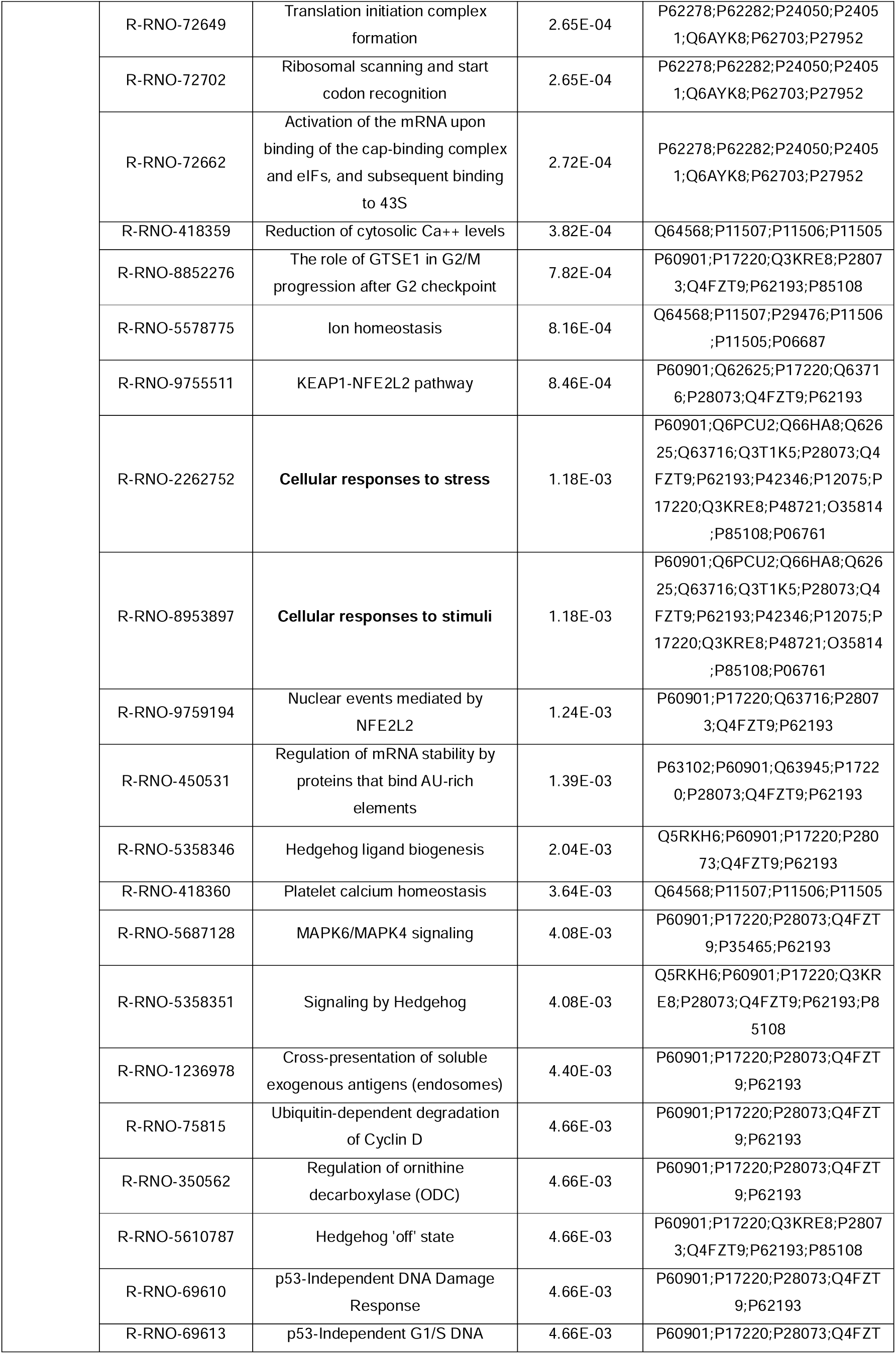
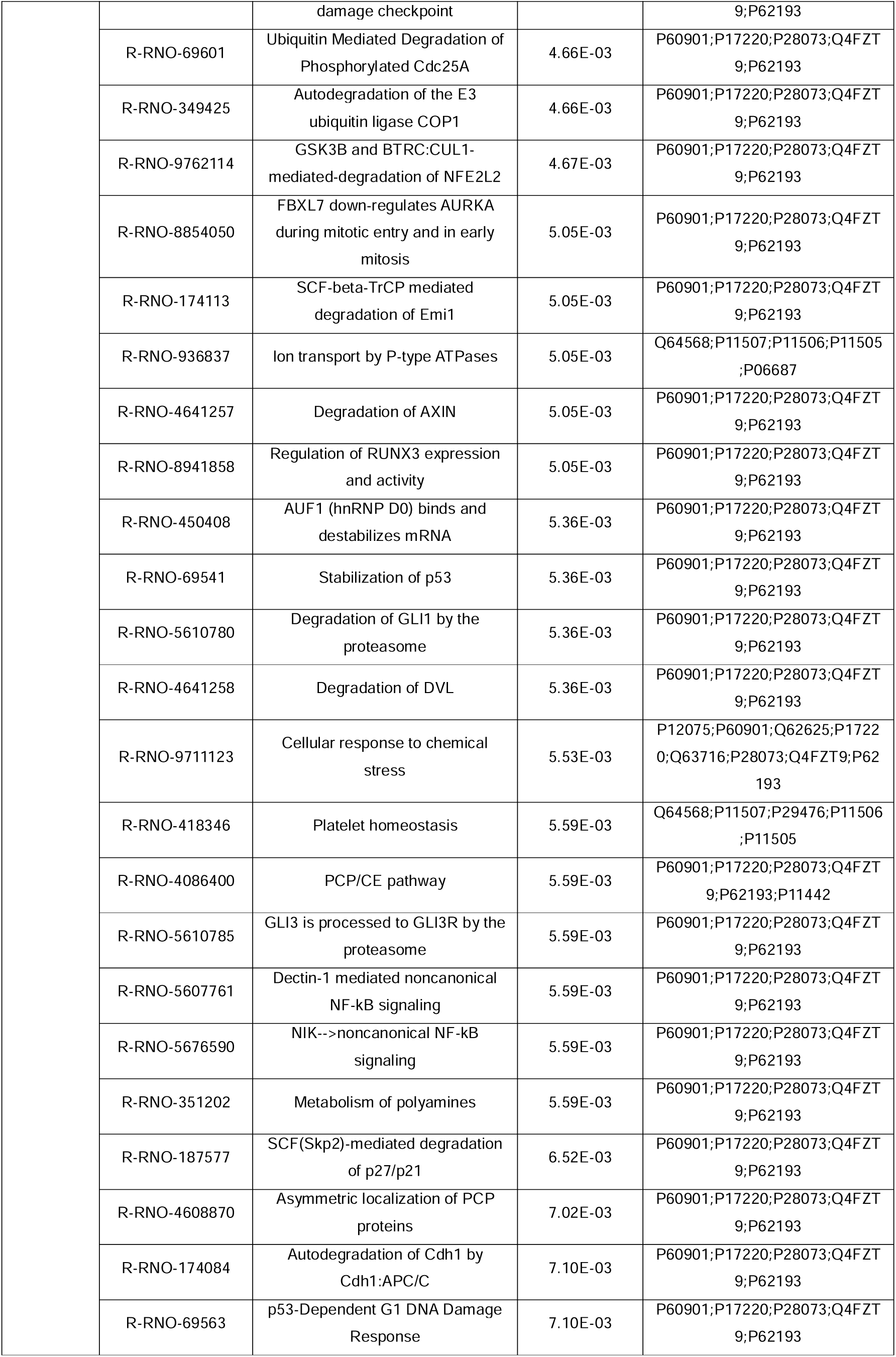
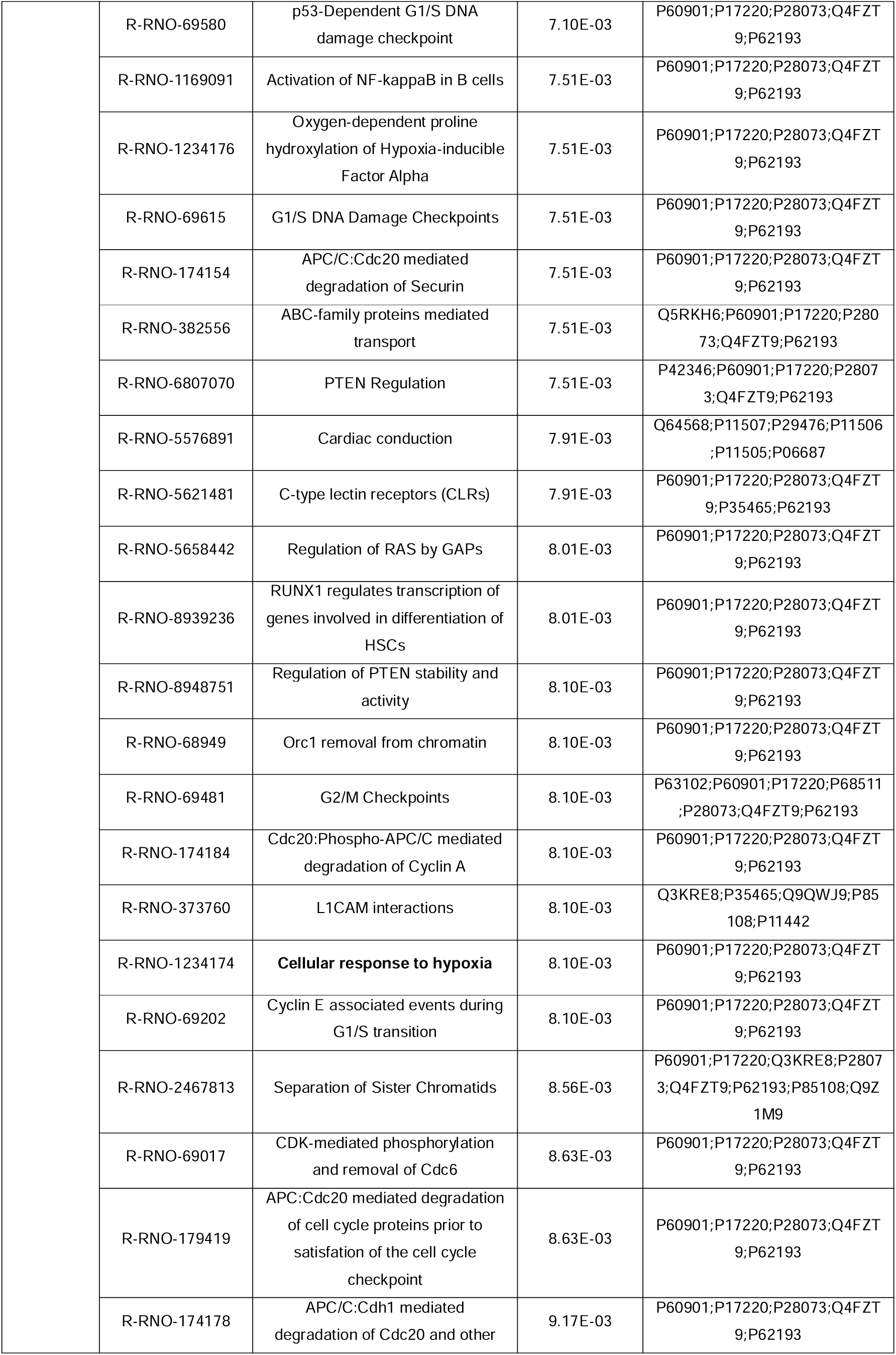
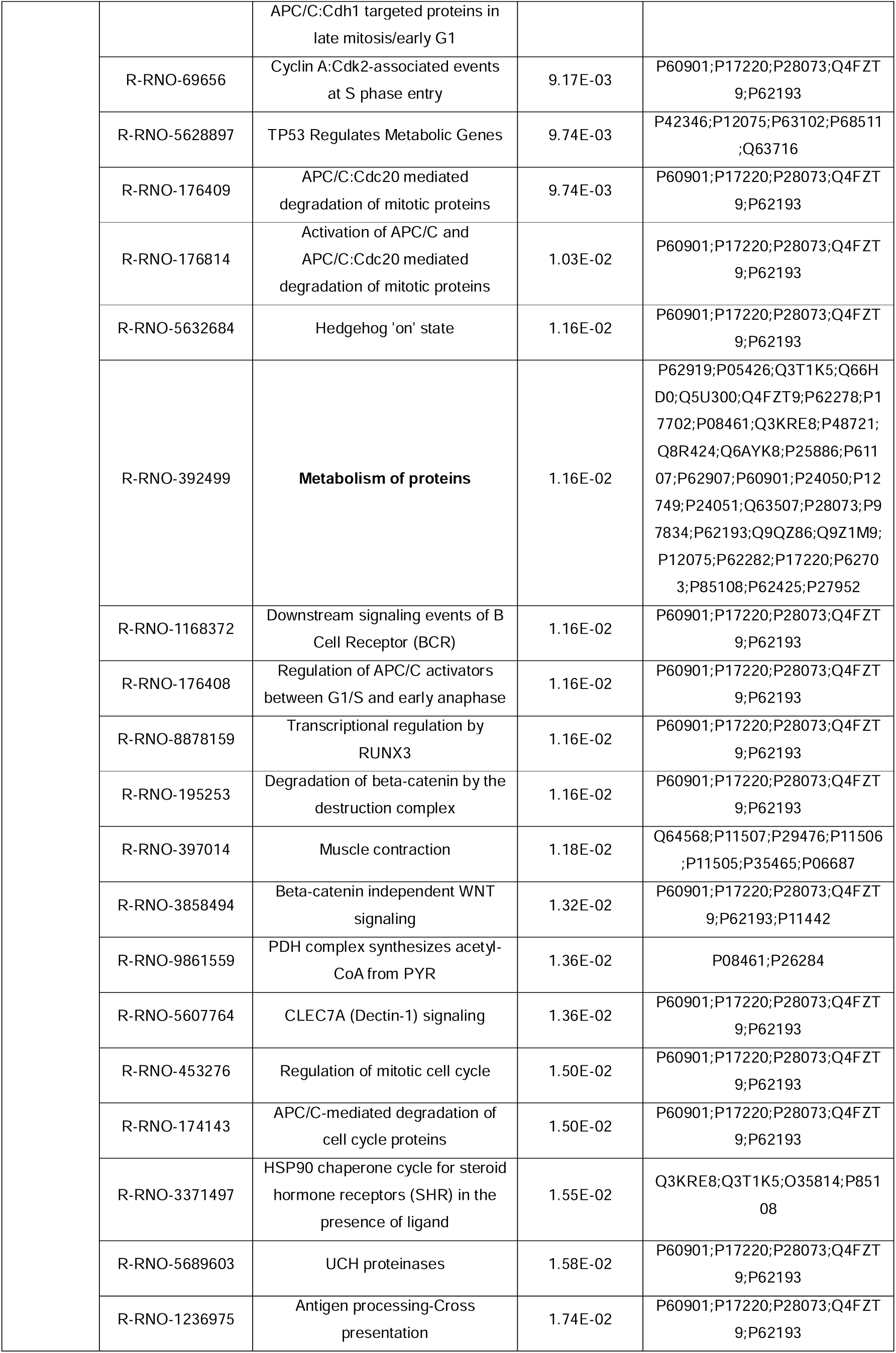
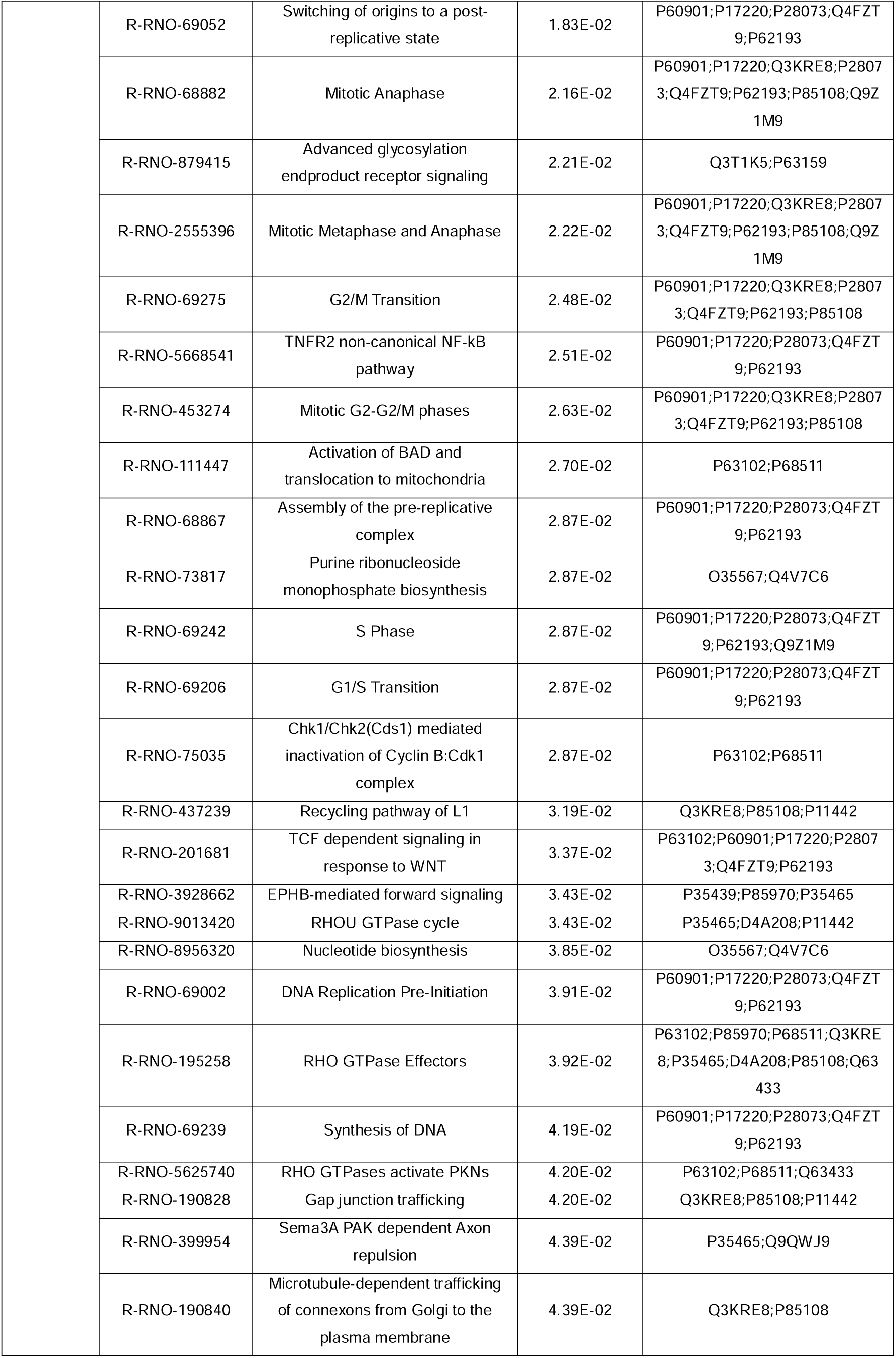
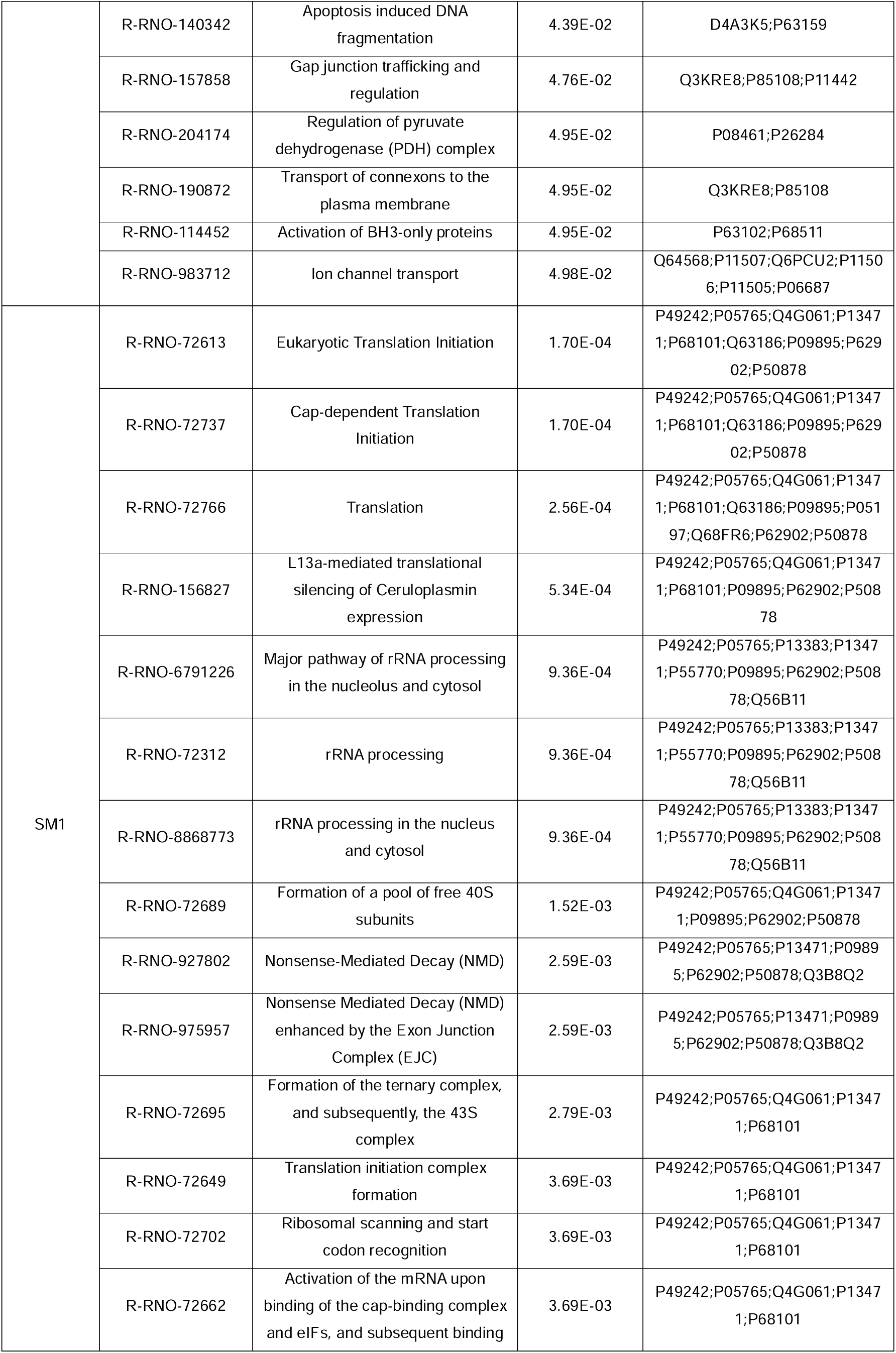
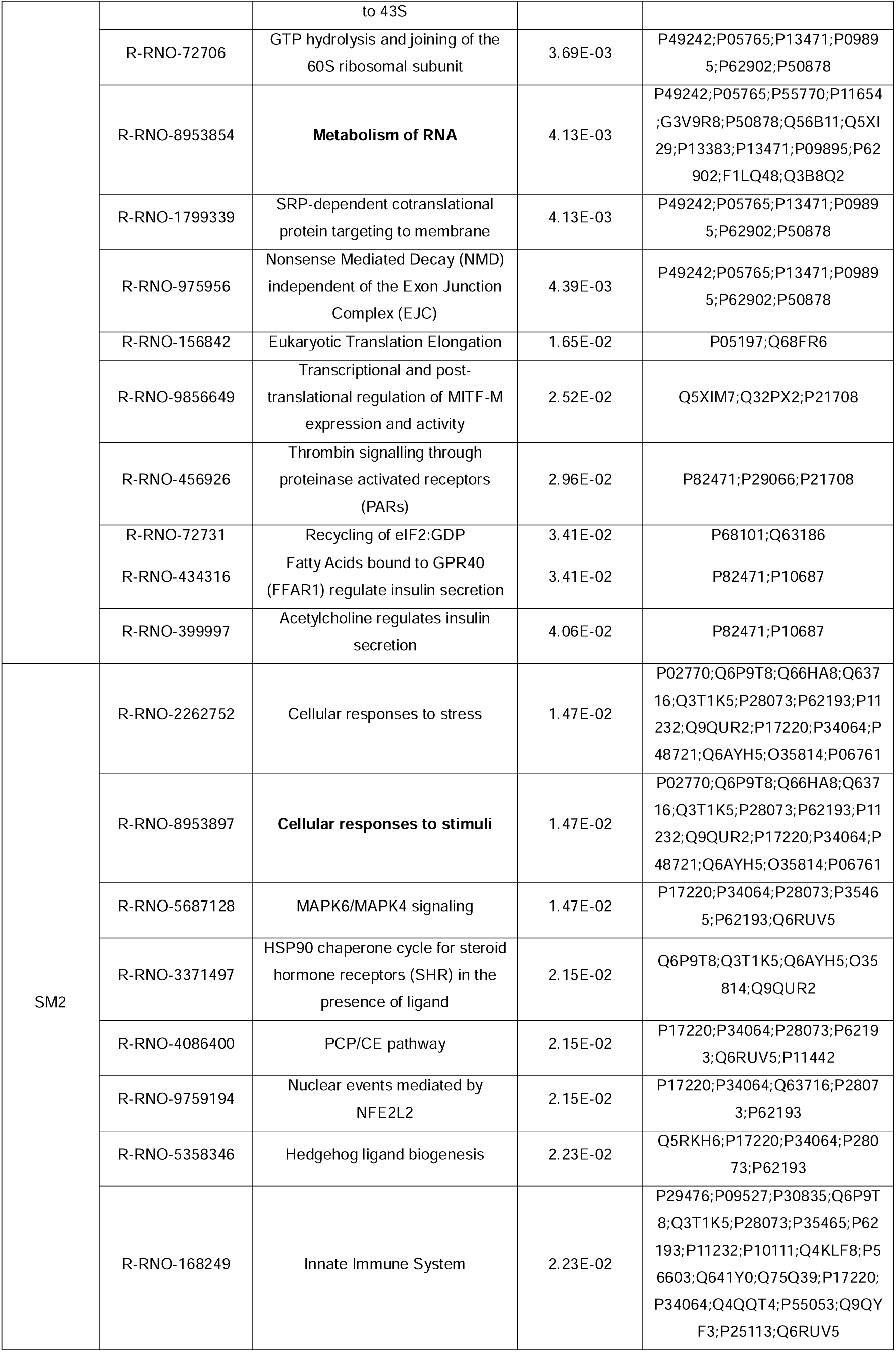
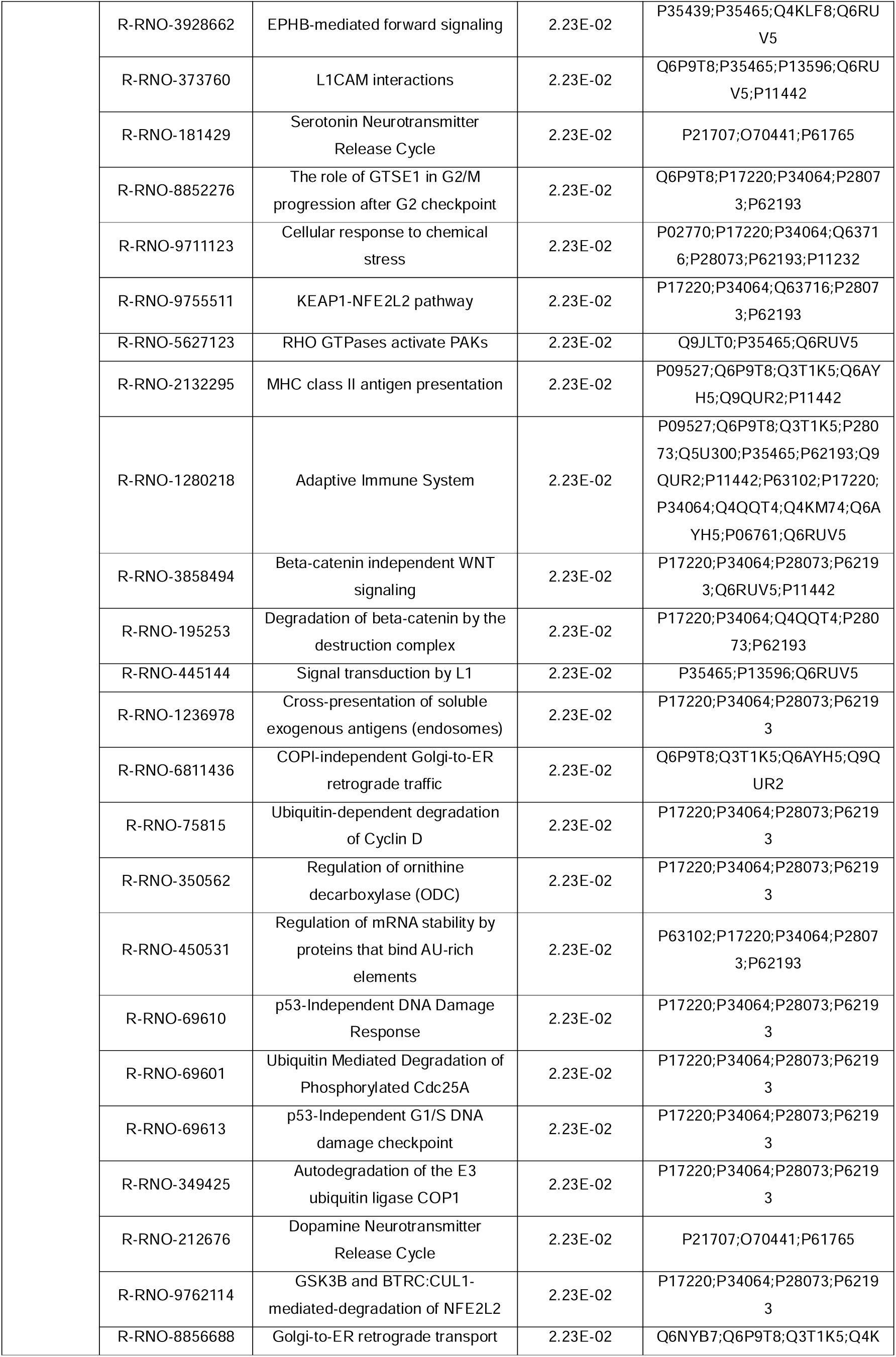
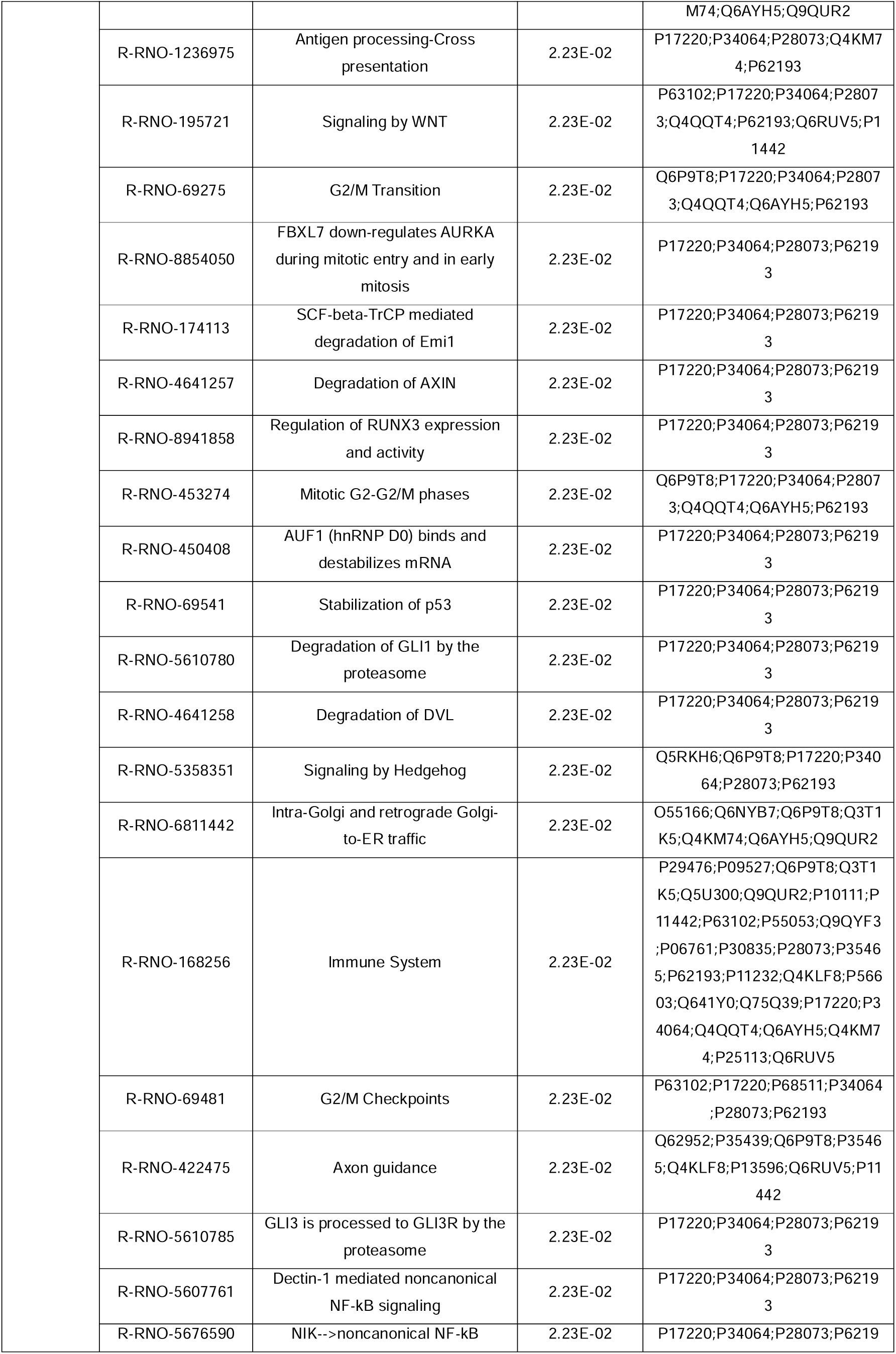
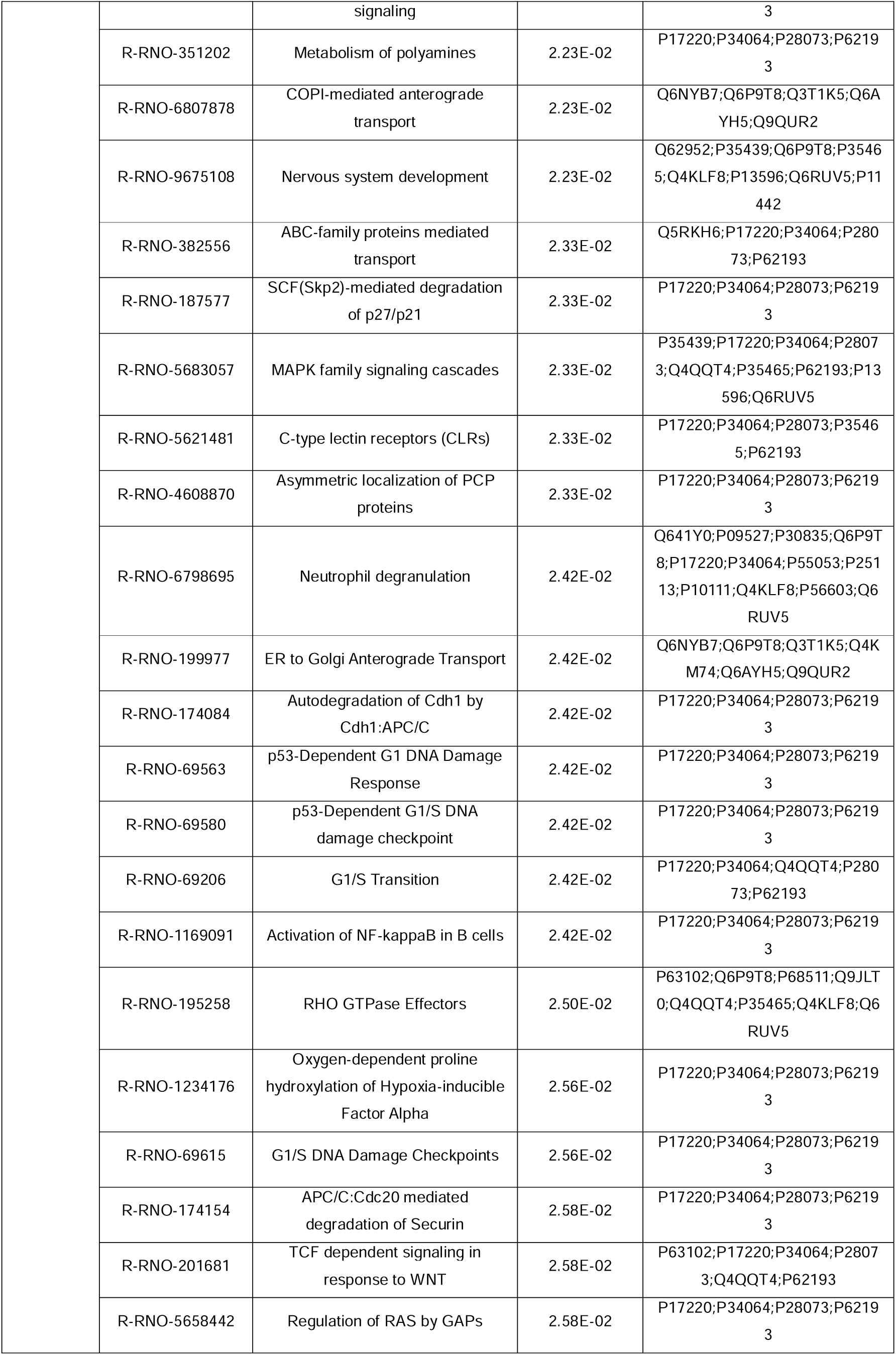
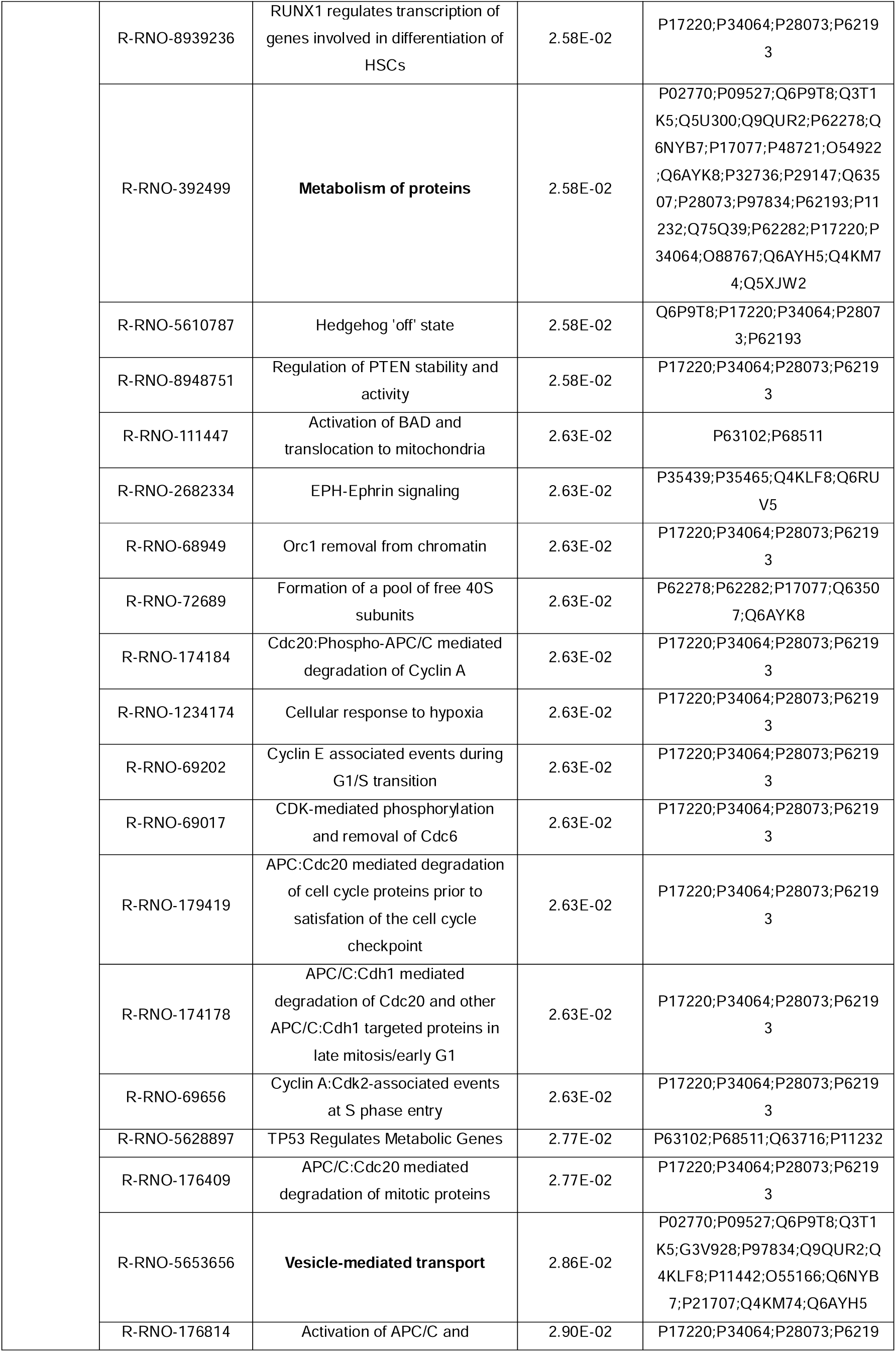
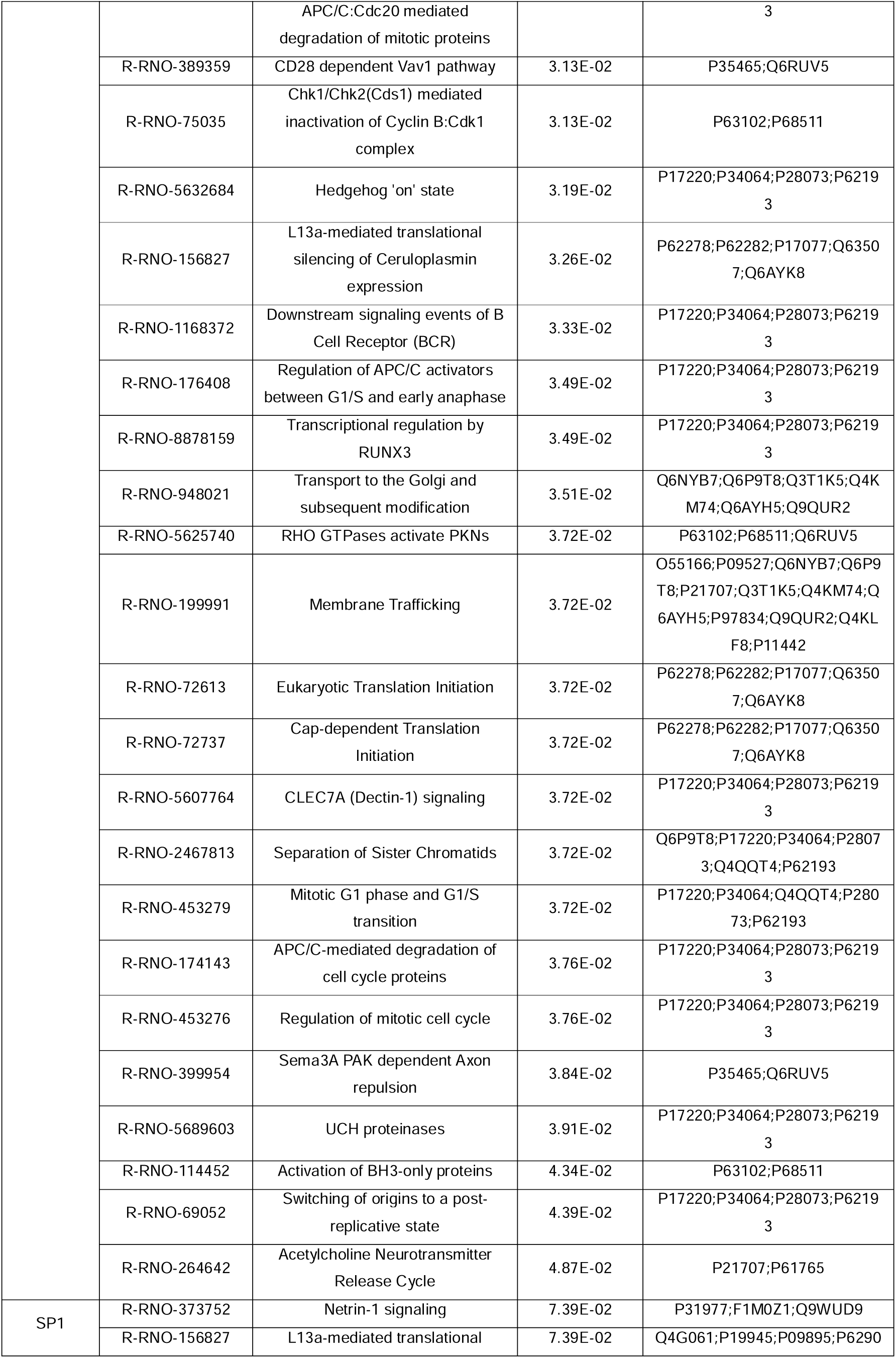
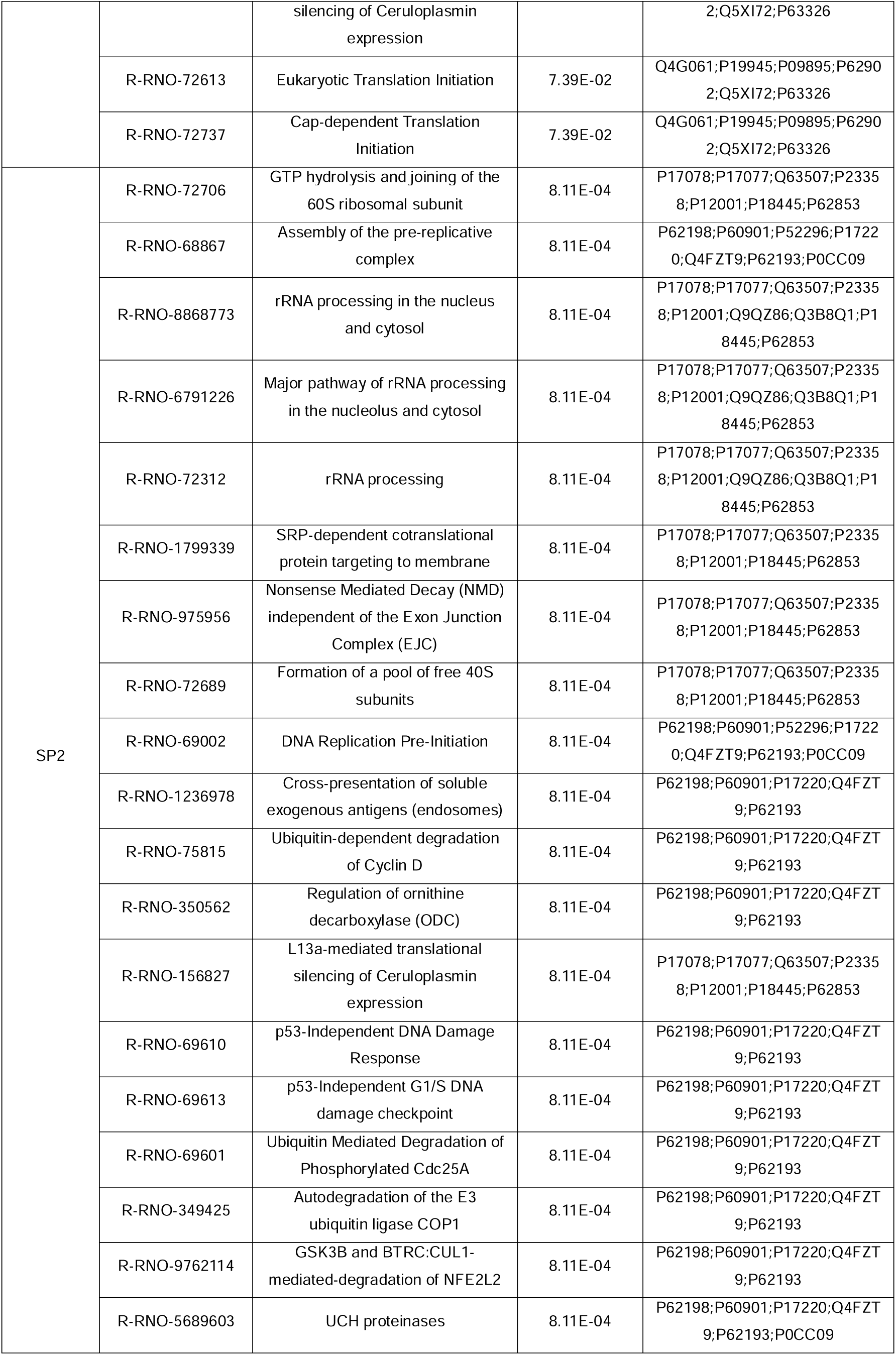
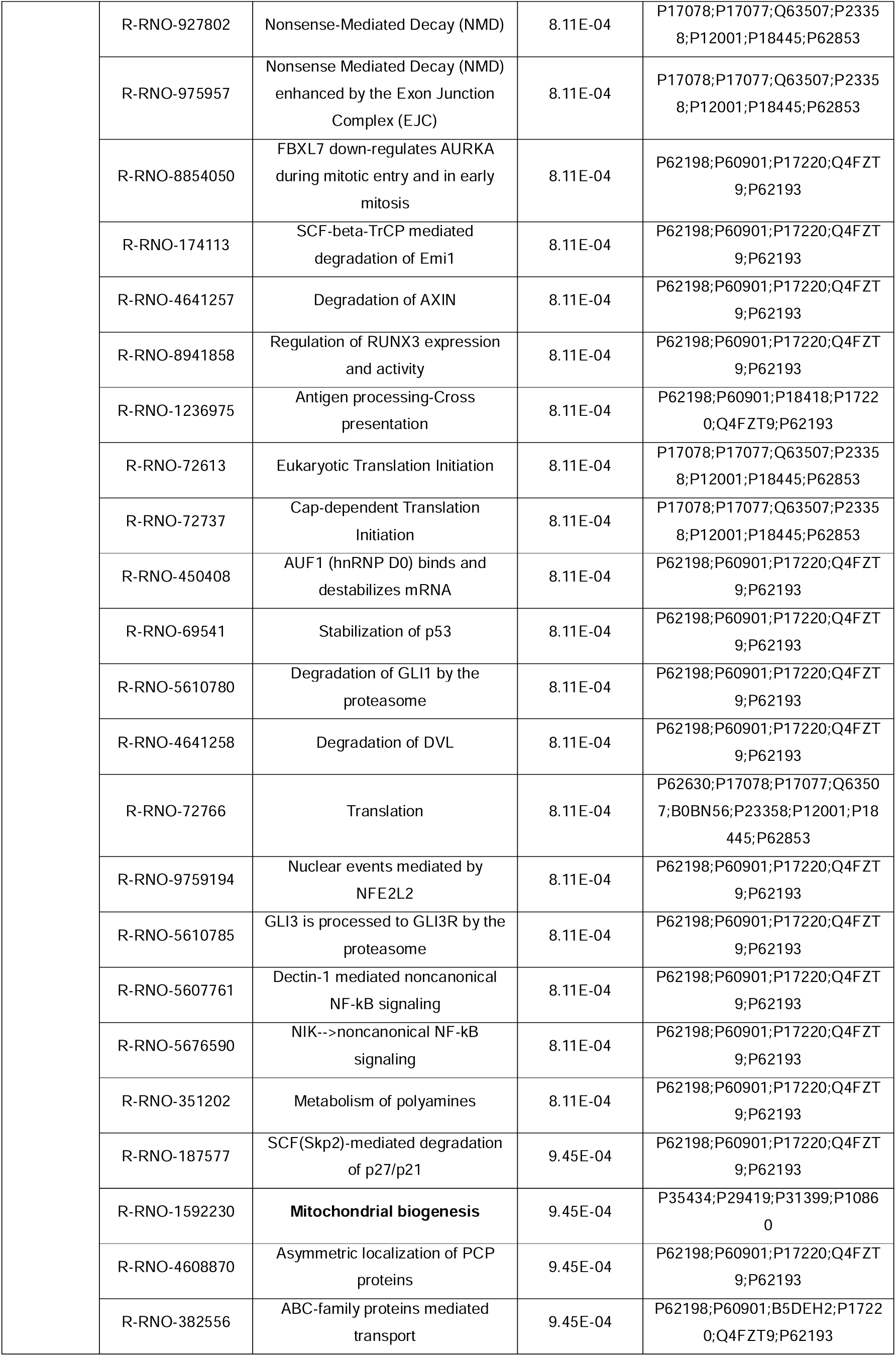
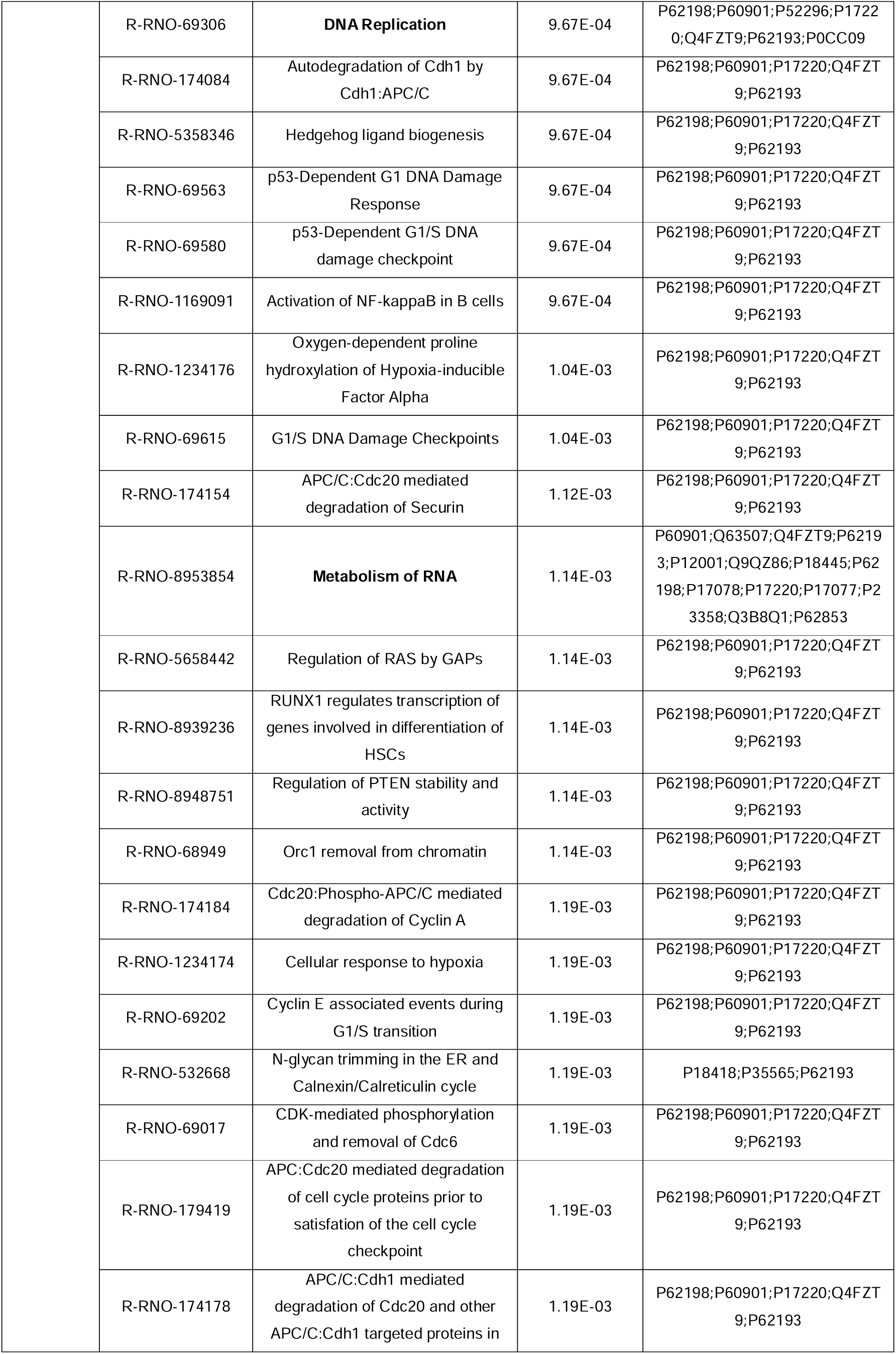
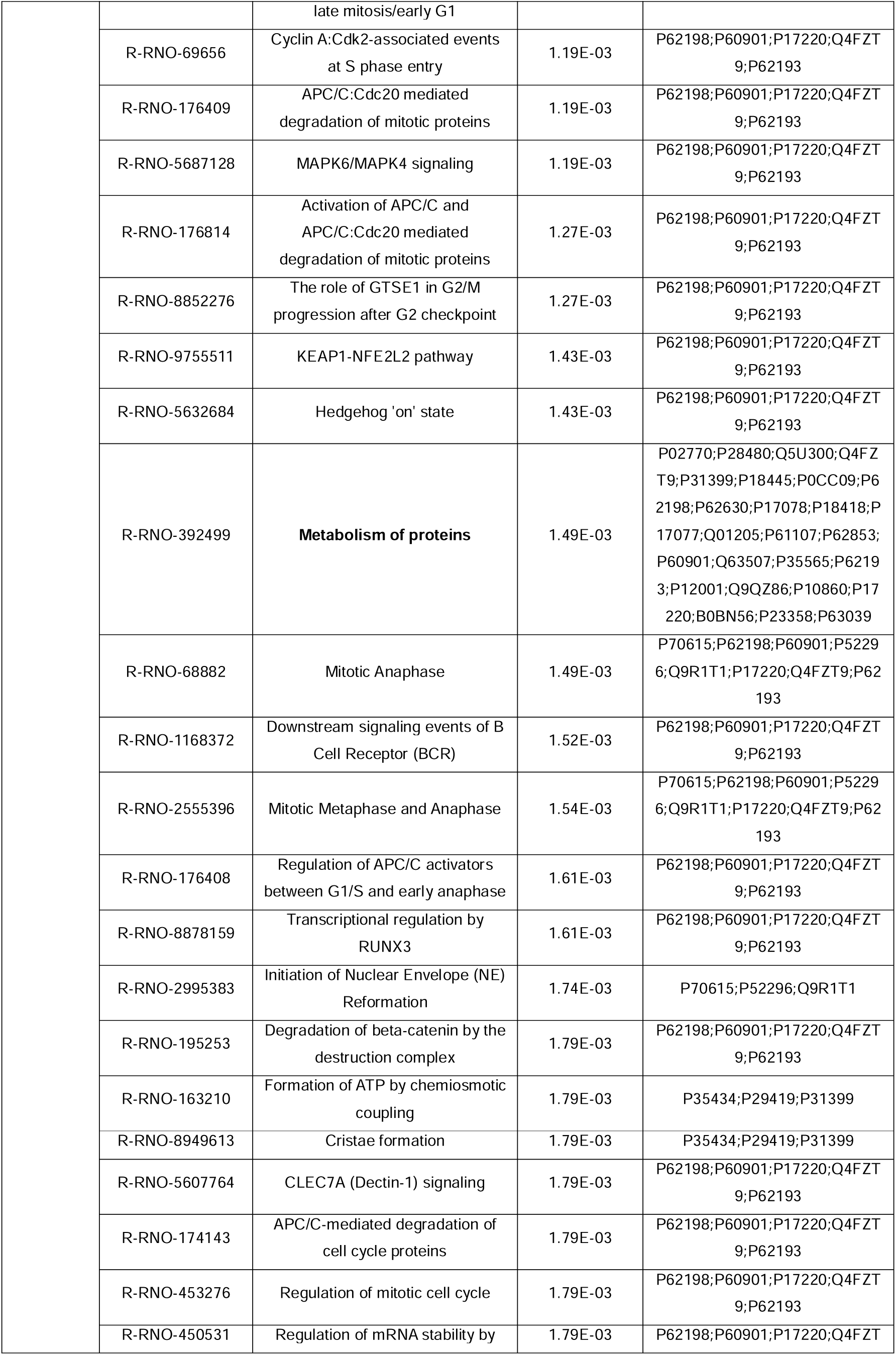
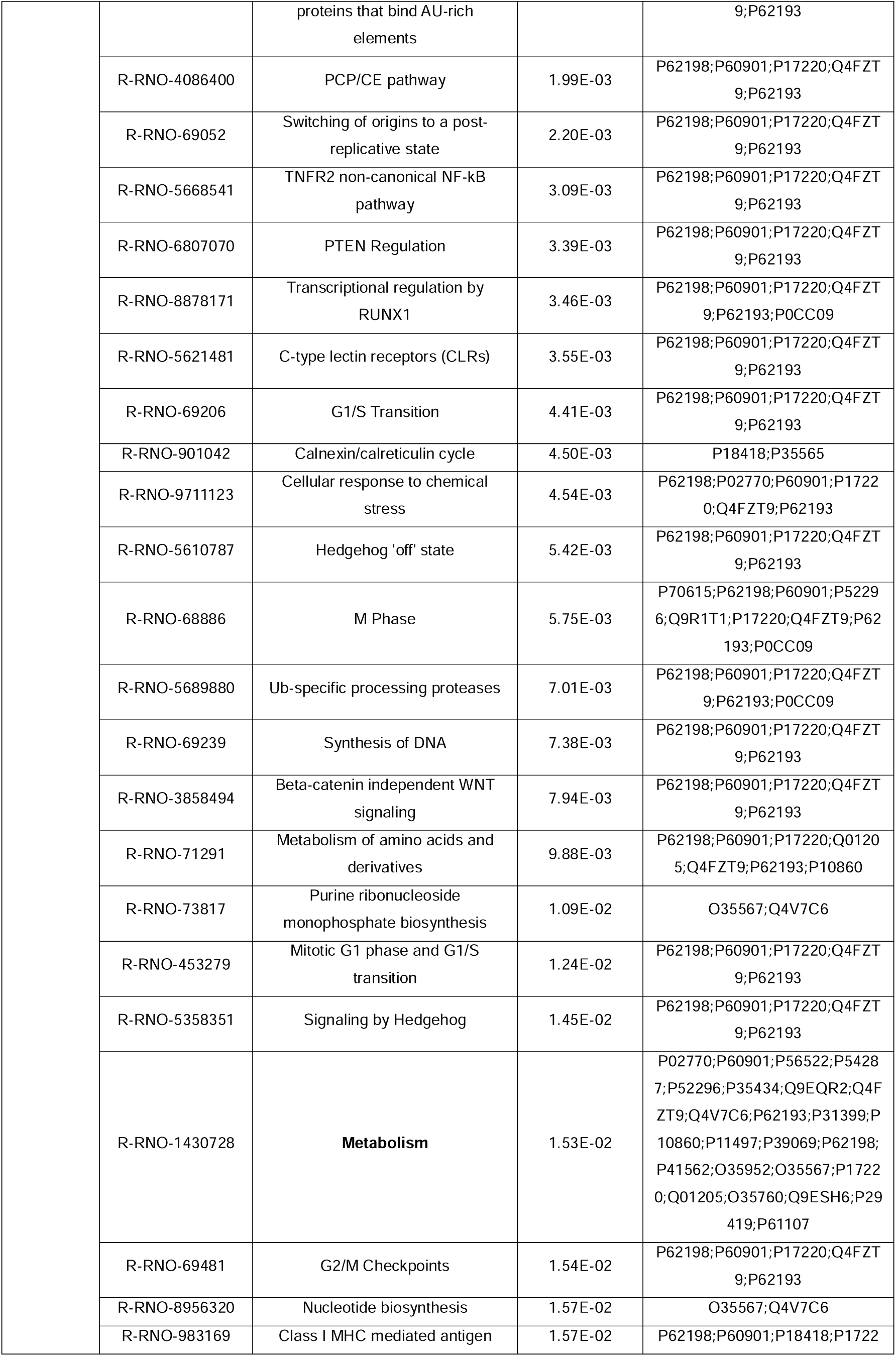
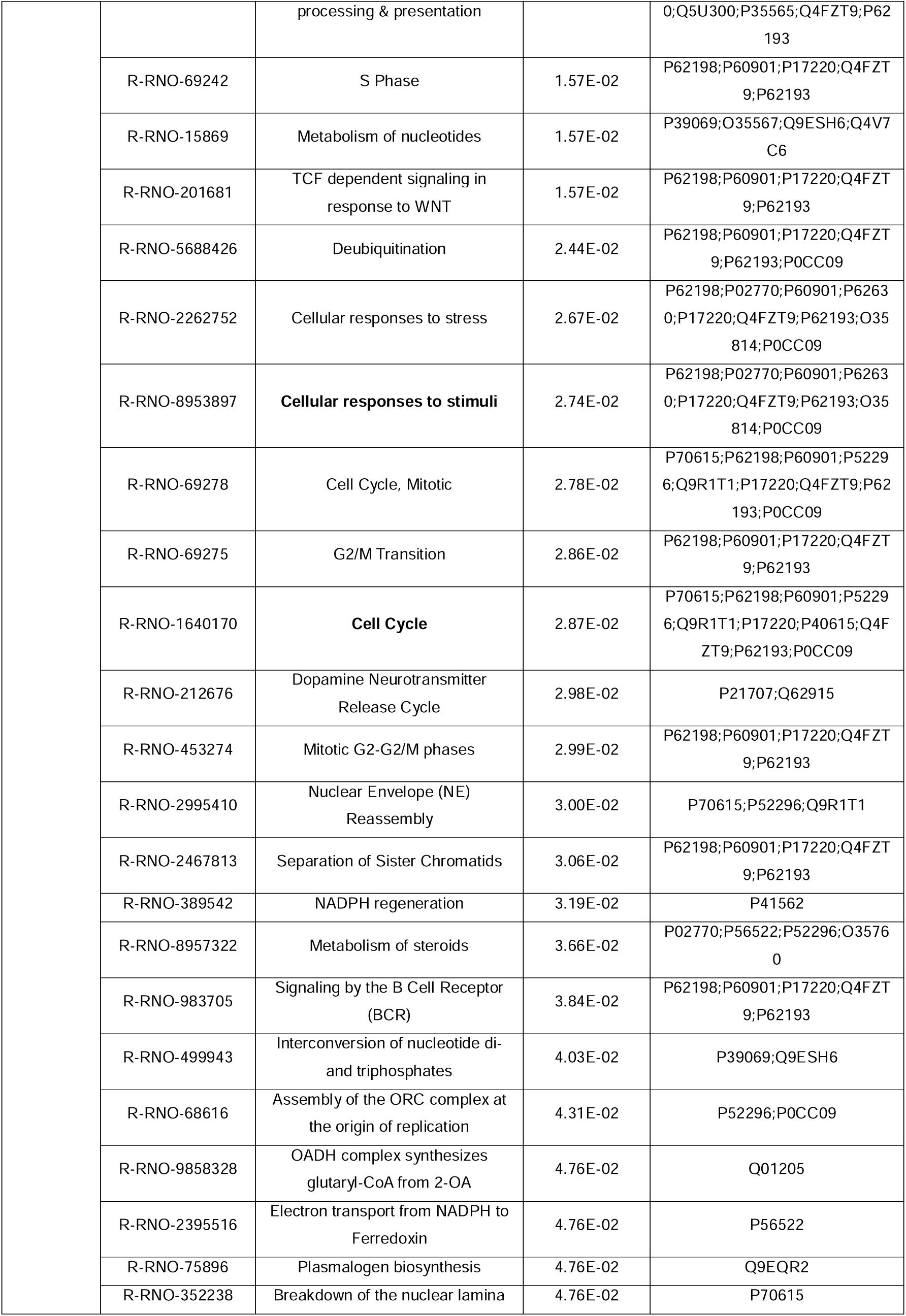
Reactome pathways of the selected clusters for neurons treated with UC-MSCs secretome (Figure 5D).

**Supplementary Table 9.** Proteins identified using a univariate approach comparing sham and OGD rescued neurons with conventional UC-MSCs’ secretome.

| Accession | Name | Mann-Whitney<br>(Sham vs OGD) (p-value) | Mann-Whitney<br>(Sham vs OGD+Standard) (p-value) |
| --- | --- | --- | --- |
| Q05175 | Brain acid soluble protein 1 | 0.015 | 0.078 |
| P16036 | Phosphate carrier protein, mitochondrial | 0.005 | 0.065 |
| **B0BNG0** | **\*ER membrane protein complex subunit 2** | **0.026** | **0.818** |
| P61314 | 60S ribosomal protein L15 | 0.030 | 0.485 |
| **P61107** | **\*Ras-related protein Rab-14** | **0.041** | **0.937** |
| P02770 | Albumin | 0.026 | 0.394 |
| P19511 | ATP synthase F(0) complex subunit B1, mitochondrial | 0.009 | 0.132 |
| P18445 | 60S ribosomal protein L27a | 0.015 | 0.065 |
| O88941 | Mannosyl-oligosaccharide glucosidase | 0.041 | 0.093 |
| P32822 | Trypsin V-B | 0.041 | 0.818 |
| P53767 | Vascular endothelial growth factor receptor 1 | 0.015 | 0.132 |
| Q6PDV7 | 60S ribosomal protein L10 | 0.015 | 0.132 |
| Q9JJM9 | Septin-5 | 0.045 | 0.065 |
| Q924S5 | Lon protease homolog, mitochondrial | 0.009 | 0.180 |
| P31399 | ATP synthase subunit d, mitochondrial | 0.009 | 0.132 |
| P09527 | Ras-related protein Rab-7a | 0.026 | 0.132 |
| P62914 | 60S ribosomal protein L11 | 0.009 | 0.093 |
| P49432 | Pyruvate dehydrogenase E1 component subunit beta, mitochondrial | 0.026 | 0.470 |
| P15865 | Histone H1.4 | 0.041 | 0.485 |
| F1LX07 | Electrogenic aspartate/glutamate antiporter SLC25A12, mitochondrial | 0.025 | 0.180 |
| P23358 | 60S ribosomal protein L12 | 0.008 | 0.378 |
| P56522 | NADPH:adrenodoxin oxidoreductase, mitochondrial | 0.019 | 0.180 |
| O08701 | Arginase-2, mitochondrial | 0.026 | 0.310 |
| P54311 | Guanine nucleotide-binding protein G(I)/G(S)/G(T) subunit beta-1 | 0.010 | 0.065 |
| P62853 | 40S ribosomal protein S25 | 0.041 | 0.485 |
| Q641Y0 | Dolichyl-diphosphooligosaccharide--protein glycosyltransferase 48 kDa subunit | 0.015 | 0.485 |
| Q5SGE0 | Leucine-rich PPR motif-containing protein, mitochondrial | 0.015 | 0.699 |
| **P62425** | **\*60S ribosomal protein L7a** | **0.016** | **0.937** |
| Q9QYF3 | Unconventional myosin-Va | 0.002 | 0.818 |
| O08875 | Serine/threonine-protein kinase DCLK1 | 0.041 | 0.078 |
| P10860 | Glutamate dehydrogenase 1, mitochondrial | 0.041 | 0.240 |
| P63039 | 60 kDa heat shock protein, mitochondrial | 0.004 | 0.180 |
| **P18163** | **\*Long-chain-fatty-acid--CoA ligase 1** | **0.045** | **0.575** |
| **P06687** | **\*Sodium/potassium-transporting ATPase subunit alpha-3** | **0.041** | **0.470** |
| **P05065** | **+Fructose-bisphosphate aldolase A** | **0.041** | **0.485** |
| F1LYQ8 | FERM, ARHGEF and pleckstrin domain-containing protein 1 | 0.026 | 0.093 |
| Q9Z2F5 | C-terminal-binding protein 1 | 0.045 | 0.297 |
| P35815 | Protein phosphatase 1B | 0.045 | 0.078 |
| F1MA98 | Nucleoprotein TPR | 0.026 | 0.485 |
| Q5BJT0 | Arginine and glutamate-rich protein 1 | 0.041 | 0.310 |
| Q32PX2 | Aminoacyl tRNA synthase complex-interacting multifunctional protein 2 | 0.015 | 0.589 |
| Q9ESI7 | Neuronal migration protein doublecortin | 0.041 | 0.180 |
| Q2PQA9 | Kinesin-1 heavy chain | 0.037 | 0.310 |
| P62815 | V-type proton ATPase subunit B, brain isoform | 0.004 | 0.065 |
| P68255 | 14-3-3 protein theta | 0.037 | 0.149 |
| **O88637** | **+Ethanolamine-phosphate cytidyltransferase** | **0.045** | **0.378** |
| **P00762** | **+Serine protease 1** | **0.041** | **0.394** |
| P84586 | RNA-binding motif protein, X chromosome retrogene-like | 0.015 | 0.699 |
| P09895 | 60S ribosomal protein L5 | 0.006 | 0.937 |
| Q63028 | Alpha-adducin | 0.026 | 0.180 |
| Q1LZ53 | DNA (cytosine-5)-methyltransferase 3A | 0.041 | 0.310 |
| P11654 | Nuclear pore membrane glycoprotein 210 | 0.026 | 0.180 |
| + for common proteins with cluster A; * for common proteins with cluster B |  |  |  |

**Supplementary Table 10.** Proteins identified using a univariate approach comparing sham and OGD treated neurons with mechano-modulated UC-MSCs’ secretome.

| **Accession** | **Name** | **Mann-Whitney**<br>**(Sham vs OGD) (p-value)** | **Mann-Whitney**<br>**(Sham vs**<br>**OGD+mechano) (p-value)** |
| --- | --- | --- | --- |
| Q05175 | Brain acid soluble protein 1 | 0.015 | 0.065 |
| Q5XIF6 | Tubulin alpha-4A chain | 0.041 | 0.310 |
| B0BNG0 | ER membrane protein complex subunit 2 | 0.026 | 0.394 |
| P61314 | 60S ribosomal protein L15 | 0.030 | 0.065 |
| P61107 | Ras-related protein Rab-14 | 0.041 | 0.423 |
| **P02770** | **+Albumin** | **0.026** | **0.818** |
| P18445 | 60S ribosomal protein L27a | 0.015 | 0.180 |
| P32822 | Trypsin V-B | 0.041 | 0.093 |
| Q6PDV7 | 60S ribosomal protein L10 | 0.015 | 0.240 |
| Q9JJM9 | Septin-5 | 0.045 | 0.336 |
| Q06647 | ATP synthase subunit | 0.041 | 0.180 |
| Q924S5 | Lon protease homolog, mitochondrial | 0.009 | 0.240 |
| P62914 | 60S ribosomal protein L11 | 0.009 | 0.310 |
| P31399 | ATP synthase subunit d, mitochondrial | 0.009 | 0.180 |
| **P09527** | **+Ras-related protein Rab-7a** | **0.026** | **0.937** |
| P54313 | Guanine nucleotide-binding protein G(I)/G(S)/G(T) subunit beta-2 | 0.015 | 0.065 |
| P49432 | Pyruvate dehydrogenase E1 component subunit beta, mitochondrial | 0.026 | 0.132 |
| P15865 | Histone H1.4 | 0.041 | 0.310 |
| F1LX07 | Electrogenic aspartate/glutamate antiporter<br>SLC25A12, mitochondrial | 0.025 | 0.132 |
| Q00715 | Histone H2B type 1 | 0.026 | 0.093 |
| P23358 | 60S ribosomal protein L12 | 0.008 | 0.173 |
| P04646 | 60S ribosomal protein L35a | 0.009 | 0.065 |
| P56522 | NADPH:adrenodoxin oxidoreductase,<br>mitochondrial | 0.019 | 0.423 |
| O08701 | Arginase-2, mitochondrial | 0.026 | 0.065 |
| P54311 | Guanine nucleotide-binding protein<br>G(I)/G(S)/G(T) subunit beta-1 | 0.010 | 0.065 |
| P62853 | 40S ribosomal protein S25 | 0.041 | 0.132 |
| **Q641Y0** | **+Dolichyl-diphosphooligosaccharide--**<br>**protein glycosyltransferase 48 kDa**<br>**subunit** | **0.015** | **0.485** |
| Q5SGE0 | Leucine-rich PPR motif-containing protein,<br>mitochondrial | 0.004 | 0.132 |
| P62425 | 60S ribosomal protein L7a | 0.016 | 0.065 |
| **Q9QYF3** | **+Unconventional myosin-Va** | **0.008** | **0.699** |
| O08875 | Serine/threonine-protein kinase DCLK1 | 0.041 | 0.132 |
| P10860 | Glutamate dehydrogenase 1, mitochondrial | 0.041 | 0.093 |
| **P18163** | **+Long-chain-fatty-acid--CoA ligase 1** | **0.045** | **0.818** |
| Q7TT49 | Serine/threonine-protein kinase MRCK beta | 0.041 | 0.054 |
| P50137 | Transketolase | 0.026 | 0.065 |
| Q64303 | Serine/threonine-protein kinase PAK 2 | 0.005 | 0.394 |
| Q9QXU8 | Cytoplasmic dynein 1 light intermediate chain<br>1 | 0.026 | 0.065 |
| Q05695 | Neural cell adhesion molecule L1 | 0.045 | 0.128 |
| P46462 | Transitional endoplasmic reticulum ATPase | 0.005 | 0.173 |
| **F1LYQ8** | **\*FERM, ARHGEF and pleckstrin domain-**<br>**containing protein 1** | **0.026** | **0.180** |
| **P13383** | **\*Nucleolin** | **0.015** | **0.818** |
| Q9Z2F5 | C-terminal-binding protein 1 | 0.045 | 0.090 |
| P63045 | Vesicle-associated membrane protein 2 | 0.026 | 0.093 |
| P35815 | Protein phosphatase 1B | 0.045 | 0.630 |
| Q5BJT0 | Arginine and glutamate-rich protein 1 | 0.041 | 0.065 |
| D3ZQG6 | Tripartite motif-containing protein 2 | 0.009 | 0.132 |
| **Q32PX2** | **\*Aminoacyl tRNA synthase complex-**<br>**interacting multifunctional protein 2** | **0.015** | **0.261** |
| P62815 | V-type proton ATPase subunit B, brain<br>isoform | 0.004 | 0.093 |
| P68255 | 14-3-3 protein theta | 0.037 | 0.078 |
| **O55165** | **\*Kinesin-like protein KIF3C** | **0.045** | **0.688** |
| **Q2PQA9** | **\*Kinesin-1 heavy chain** | **0.002** | **0.310** |
| Q8K3F3 | Protein phosphatase 1 regulatory subunit<br>14B | 0.009 | 0.109 |
| Q7TMB7 | Phospholipid phosphatase-related protein<br>type 4 | 0.009 | 0.093 |
| O88637 | Ethanolamine-phosphate cytidyltransferase | 0.045 | 0.229 |
| Q6AXU6 | Jupiter microtubule associated homolog 1 | 0.015 | 0.132 |
| **D3ZE55** | **\*Protocadherin-8** | **0.026** | **0.937** |
| P84586 | RNA-binding motif protein, X chromosome<br>retrogene-like | 0.015 | 0.065 |
| Q99376 | Transferrin receptor protein 1 | 0.005 | 0.065 |
| **P09895** | **\*60S ribosomal protein L5** | **0.006** | **0.589** |
| Q6MG08 | ATP-binding cassette sub-family F member 1 | 0.009 | 0.065 |
| Q4FZY0 | EF-hand domain-containing protein D2 | 0.026 | 0.132 |
| **Q66H80** | **\*Coatomer subunit delta** | **0.009** | **0.310** |
| Q2IBD4 | Cortactin-binding protein 2 | 0.009 | 0.065 |
| **P11654** | **\*Nuclear pore membrane glycoprotein 210** | **0.026** | **0.310** |
| + for common proteins with cluster C; * for common proteins with cluster D |  |  |  |

**Supplementary Table 11.** Proteins identified using a univariate approach comparing sham and OGD treated neurons with physioxia-modulated UC-MSCs’ secretome.

| Acession | Name | Mann-Whitney<br>(Sham vs OGD) (p-value) | Mann-Whitney<br>(Sham vs<br>OGD+physioxia) (p-value) |
| --- | --- | --- | --- |
| Q5XIF6 | Tubulin alpha-4A chain | 0.041 | 0.065 |
| P31044 | Phosphatidylethanolamine-binding protein 1 | 0.041 | 0.180 |
| B0BNG0 | ER membrane protein complex subunit 2 | 0.026 | 0.310 |
| P61314 | 60S ribosomal protein L15 | 0.030 | 0.065 |
| **P61107** | **\*Ras-related protein Rab-14** | **0.041** | **0.485** |
| **P02770** | **\*Albumin** | **0.026** | **0.937** |
| P19511 | ATP synthase F(0) complex subunit B1,<br>mitochondrial | 0.009 | 0.093 |
| **P18445** | **\*60S ribosomal protein L27a** | **0.015** | **0.262** |
| O88941 | Mannosyl-oligosaccharide glucosidase | 0.041 | 0.310 |
| **P32822** | **\*Trypsin V-B** | **0.041** | **0.336** |
| **P53767** | **\*Vascular endothelial growth factor**<br>**receptor 1** | **0.015** | **0.180** |
| Q6PDV7 | 60S ribosomal protein L10 | 0.015 | 0.132 |
| Q9JJM9 | Septin-5 | 0.045 | 0.132 |
| Q924S5 | Lon protease homolog, mitochondrial | 0.009 | 0.310 |
| P62914 | 60S ribosomal protein L11 | 0.009 | 0.310 |
| **P31399** | **\*ATP synthase subunit d, mitochondrial** | **0.009** | **0.699** |
| P09527 | Ras-related protein Rab-7a | 0.026 | 0.065 |
| P15865 | Histone H1.4 | 0.041 | 0.180 |
| F1LX07 | Electrogenic aspartate/glutamate antiporter<br>SLC25A12, mitochondrial | 0.025 | 0.229 |
| P12075 | Cytochrome c oxidase subunit 5B,<br>mitochondrial | 0.026 | 0.394 |
| **P23358** | **\*60S ribosomal protein L12** | **0.008** | **0.575** |
| P04646 | 60S ribosomal protein L35a | 0.009 | 0.093 |
| **P56522** | **\*NADPH:adrenodoxin oxidoreductase,**<br>**mitochondrial** | **0.019** | **0.937** |
| O08701 | Arginase-2, mitochondrial | 0.026 | 0.180 |
| **P62853** | **\*40S ribosomal protein S25** | **0.041** | **0.818** |
| **Q641Y0** | **\*Dolichyl-diphosphooligosaccharide--**<br>**protein glycosyltransferase 48 kDa**<br>**subunit** | **0.015** | **0.093** |
| **P18418** | **\*Calreticulin** | **0.041** | **0.589** |
| Q8VHF5 | Citrate synthase, mitochondrial | 0.005 | 0.065 |
| Q64428 | Trifunctional enzyme subunit alpha,<br>mitochondrial | 0.026 | 0.336 |
| Q9QYF3 | Unconventional myosin-Va | 0.004 | 0.132 |
| **P10860** | **\*Glutamate dehydrogenase 1,**<br>**mitochondrial** | **0.041** | **0.937** |
| P18163 | Long-chain-fatty-acid--CoA ligase 1 | 0.045 | 0.699 |
| **P63039** | **\*60 kDa heat shock protein, mitochondrial** | **0.013** | **0.378** |
| Q7TT49 | Serine/threonine-protein kinase MRCK beta | 0.041 | 0.394 |
| **P05065** | **+Fructose-bisphosphate aldolase A** | **0.041** | **0.470** |
| **Q7M767** | **+Ubiquitin-conjugating enzyme E2 variant**<br>**2** | **0.045** | **0.485** |
| P46462 | Transitional endoplasmic reticulum ATPase | 0.030 | 0.173 |
| Q5RKH0 | Cytokine-like nuclear factor N-PAC | 0.026 | 0.149 |
| Q05695 | Neural cell adhesion molecule L1 | 0.045 | 0.092 |
| **F1LYQ8** | **+FERM, ARHGEF and pleckstrin domain-**<br>**containing protein 1** | **0.026** | **0.873** |
| **Q9Z2F5** | **+C-terminal-binding protein 1** | **0.045** | **0.378** |
| **P13383** | **+Nucleolin** | **0.015** | **0.335** |
| P35815 | Protein phosphatase 1B | 0.045 | 0.065 |
| **Q5BJT0** | **+Arginine and glutamate-rich protein 1** | **0.041** | **0.937** |
| D3ZQG6 | Tripartite motif-containing protein 2 | 0.009 | 0.065 |
| **Q32PX2** | **+Aminoacyl tRNA synthase complex-**<br>**interacting multifunctional protein 2** | **0.015** | **0.240** |
| **Q2PQA9** | **+Kinesin-1 heavy chain** | **0.041** | **1.000** |
| Q9ESI7 | Neuronal migration protein doublecortin | 0.041 | 0.132 |
| Q7TMB7 | Phospholipid phosphatase-related protein<br>type 4 | 0.009 | 0.093 |
| **P68255** | **+14-3-3 protein theta** | **0.037** | **0.699** |
| O55165 | Kinesin-like protein KIF3C | 0.045 | 0.297 |
| Q8K3F3 | Protein phosphatase 1 regulatory subunit<br>14B | 0.009 | 0.093 |
| P00762 | Serine protease 1 | 0.041 | 0.262 |
| P17425 | Hydroxymethylglutaryl-CoA synthase,<br>cytoplasmic | 0.009 | 0.065 |
| **P09895** | **+60S ribosomal protein L5** | **0.006** | **0.937** |
| Q9QUH6 | Ras/Rap GTPase-activating protein SynGAP | 0.026 | 0.065 |
| Q71UE8 | NEDD8 | 0.009 | 0.228 |
| Q1LZ53 | DNA (cytosine-5)-methyltransferase 3A | 0.041 | 0.093 |
| **P11654** | **+Nuclear pore membrane glycoprotein**<br>**210** | **0.026** | **0.485** |
| Q9ES73 | Melanoma-associated antigen D1 | 0.026 | 0.132 |
| Q5XIM5 | Protein CDV3 homolog | 0.009 | 0.132 |

## Abbreviations

BSA: Bovine serum albumin
CM: Conditioned medium
DEPs: Differentially expressed proteins
DIV: Days in vitro
eIF2_α_: GTP-linked eukaryotic initiation factor
FDR: False Discovery Rate
HIE: Hypoxic ischemic encephalopathy
ICC: Immunocytochemistry
LC-MS/MS: Liquid Chromatography coupled to tandem mass spectrometry
OCR: Oxygen consumption rate
OGD: Oxygen Glucose Deprivation
PA: Perinatal asphyxia
PERK: PKR-like endoplasmic reticulum eIF2_α_ kinase
PVDF: Low fluorescence polyvinylidene fluoride
TCPS: Tissue culture polystyrene
TH: Therapeutical hypothermia
UC-MSCs: Umbilical cord Mesenchymal Stem Cells
VEGFR1: Vascular endothelial growth factor receptor 1
WB: Western Blot

